# Mosaic sarbecovirus nanoparticles elicit cross-reactive responses in pre-vaccinated animals

**DOI:** 10.1101/2024.02.08.576722

**Authors:** Alexander A. Cohen, Jennifer R. Keeffe, Ariën Schiepers, Sandra E. Dross, Allison J. Greaney, Annie V. Rorick, Han Gao, Priyanthi N.P. Gnanapragasam, Chengcheng Fan, Anthony P. West, Arlene I. Ramsingh, Jesse H. Erasmus, Janice D. Pata, Hiromi Muramatsu, Norbert Pardi, Paulo J.C. Lin, Scott Baxter, Rita Cruz, Martina Quintanar-Audelo, Ellis Robb, Cristina Serrano-Amatriain, Leonardo Magneschi, Ian G. Fotheringham, Deborah H. Fuller, Gabriel D. Victora, Pamela J. Bjorkman

## Abstract

Immunization with mosaic-8b [60-mer nanoparticles presenting 8 SARS-like betacoronavirus (sarbecovirus) receptor-binding domains (RBDs)] elicits more broadly cross-reactive antibodies than homotypic SARS-CoV-2 RBD-only nanoparticles and protects against sarbecoviruses. To investigate original antigenic sin (OAS) effects on mosaic-8b efficacy, we evaluated effects of prior COVID-19 vaccinations in non-human primates and mice on anti-sarbecovirus responses elicited by mosaic-8b, admix-8b (8 homotypics), or homotypic SARS-CoV-2 immunizations, finding greatest cross-reactivity for mosaic-8b. As demonstrated by molecular fate-mapping in which antibodies from specific cohorts of B cells are differentially detected, B cells primed by WA1 spike mRNA-LNP dominated antibody responses after RBD-nanoparticle boosting. While mosaic-8b- and homotypic-nanoparticles boosted cross-reactive antibodies, de novo antibodies were predominantly induced by mosaic-8b, and these were specific for variant RBDs with increased identity to RBDs on mosaic-8b. These results inform OAS mechanisms and support using mosaic-8b to protect COVID-19 vaccinated/infected humans against as-yet-unknown SARS-CoV-2 variants and animal sarbecoviruses with human spillover potential.

## Introduction

Spillover of animal SARS-like betacoronaviruses (sarbecoviruses) resulted in two health crises in the past 20 years: the SARS-CoV (SARS-1) epidemic in the early 2000s and the recent COVID-19 pandemic caused by SARS-CoV-2 (SARS-2). Cross-species transmission of a new sarbecovirus that is easily spread between humans from reservoirs in bats or other animals could result in another pandemic.^1–3^ In addition, COVID-19 continues to be a health concern due to long COVID complications^4^ as well as SARS-2 variants of concern (VOCs) that show apparently increased transmissibility and/or resistance to neutralizing antibodies (Abs) elicited by infection or vaccination.^5–8^ For example, in Omicron VOCs, substitutions in the SARS-2 spike protein receptor-binding domains (RBDs), the major target of neutralizing Abs and detectable cross-variant neutralization,^9,10^ have reduced the efficacies of vaccines and therapeutic monoclonal Abs (mAbs).^8,11^ One strategy to increase protective responses to SARS-2 VOCs involves novel variant vaccine boosters.^12^ Since repeated updating of COVID vaccines is impractical and expensive, a more optimal strategy would be a vaccine that does not require changing to protect against both emerging sarbecoviruses and SARS-2 VOCs.

To develop a vaccine that would protect against unknown sarbecoviruses and new SARS-2 variants, we used an approach involving simultaneous display of eight different sarbecovirus RBDs arranged randomly on protein-based 60-mer nanoparticles (mosaic-8b RBD-nanoparticles) (Figure 1A) and evaluated Ab responses against RBDs representing both matched (RBD on the nanoparticle) and mismatched (RBD not on the nanoparticle) viruses.^13^ We found that mosaic-8b nanoparticles showed enhanced heterologous binding, neutralization, and protection from sarbecovirus challenges compared with homotypic (SARS-2 RBD only) nanoparticles in animal models.^13,14^ For example, mosaic-8b immunizations showed protection in K18-hACE2 transgenic mice, a stringent model of coronavirus infection,^15^ from both SARS-2 and a mismatched SARS-1 challenge, whereas homotypic SARS-2 immunized mice were protected only from SARS-2 challenge.^14^

**Figure 1.**
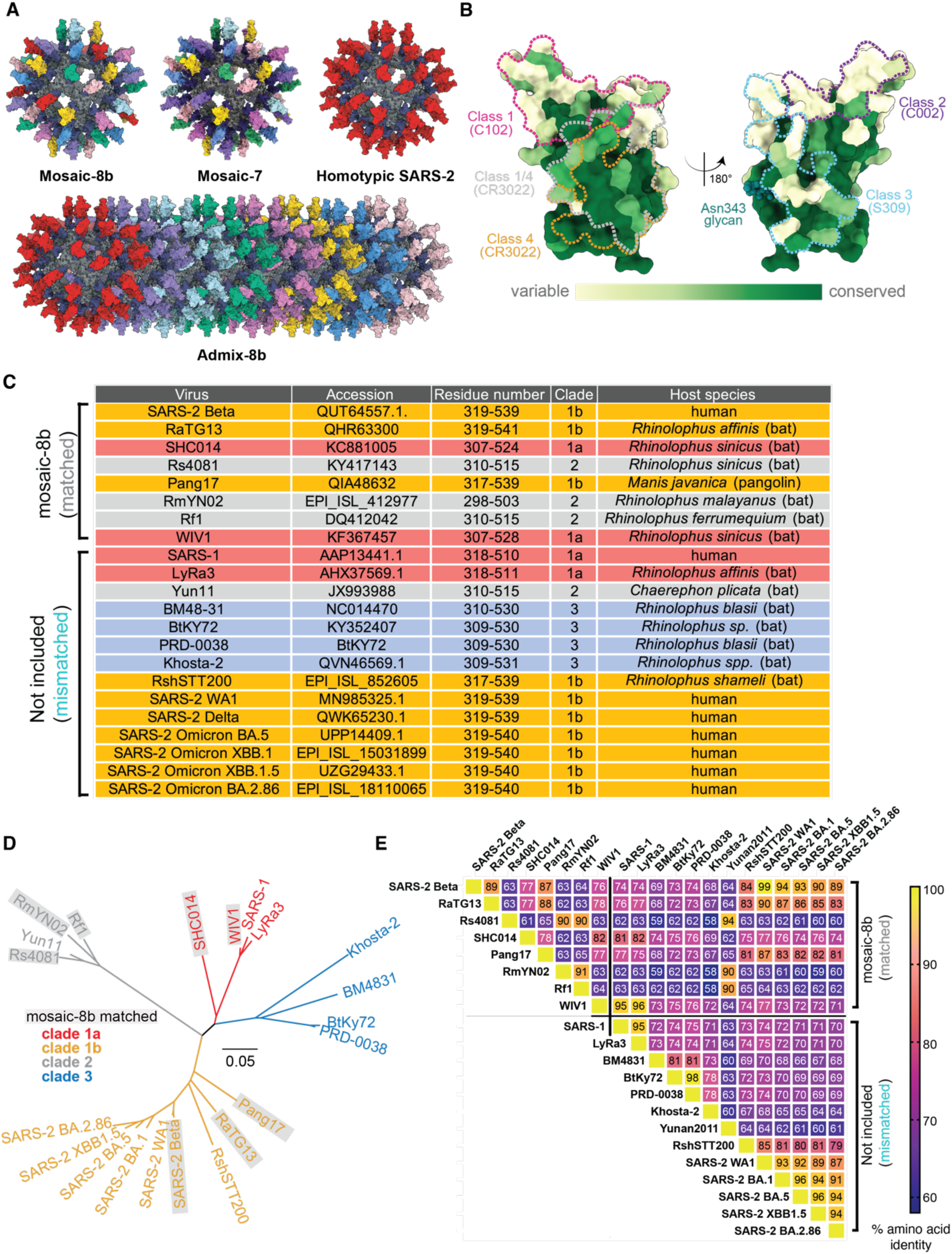
RBDs used to make nanoparticles and for assays, related to Figure S1, Table S1. (A) Models of mosaic-8b, mosaic-7 (mosaic-8b without SARS-2 RBD), homotypic SARS-2, and admix-8b RBD-nanoparticles constructed using coordinates of an RBD (PDB 7BZ5), SpyCatcher (PDB 4MLI), and i3-01 nanoparticle (PDB 7B3Y). (B) Sequence conservation determined using the ConSurf Database^95^ of the 16 sarbecovirus RBDs used to make nanoparticles and/or for assays shown on two views of an RBD surface (PDB 7BZ5). Class 1, 2, 3, 4, and 1/4 epitopes are outlined in different colors using information from Fab-RBD or Fab-spike trimer structures (C102, PDB 7K8M; C002, PDB 7K8T; S309, PDB 7JX3; CR3022, PDB 7LOP; and C118, PDB 7RKV). (C) List of sarbecoviruses from which the RBDs in mosaic-8b and admix-8b were included (matched) or not included (mismatched). Clades were defined as described.^96^ (D) Phylogenetic tree of selected sarbecoviruses calculated using a Jukes-Cantor generic distance model using Geneious Prime® 2023.1.2 based on amino acid sequences of RBDs aligned using Clustal Omega.^97^ Viruses with RBDs included in mosaic-8b are highlighted in gray rectangles. (E) Amino acid sequence identity matrix of RBDs based on alignments using Clustal Omega.^97^

To explain the increased cross-reactivity of Abs elicited by mosaic-8b, we hypothesized that B cells with B cell receptors (BCRs) that can crosslink using both of their antigen-binding Fab arms between adjacent non-identical RBDs to allow recognition of conserved epitopes would be preferentially stimulated to produce cross-reactive Abs, as compared with B cells presenting BCRs that bind to variable epitopes, which could rarely, if ever, crosslink between non-identical RBDs arranged randomly on a nanoparticle^14^ (Figure S1A). By contrast, homotypic RBD-nanoparticles presenting identical RBDs are predicted to bind BCRs against immunodominant strain-specific epitopes presented on adjacent identical RBDs. Epitope mapping of polyclonal antisera elicited by mosaic-8b versus homotypic SARS-2 RBD-nanoparticles using deep mutational scanning (DMS)^16^ provided evidence supporting this model: Abs from mosaic-8b antisera primarily targeted more conserved class 4 and class 1/4 RBD epitopes (Figure 1B) (RBD epitope nomenclature from ref.^17^) that show less variability in sarbecoviruses and SARS-2 VOCs because they contact other portions of the spike trimer,^14^ whereas homotypic antiserum Abs primarily targeted variable class 1 and 2 RBD epitope regions that are more accessible and are not involved in contacts with non-RBD portions of spike.^14^ These results, combined with animal experiments showing promising cross-reactive binding and neutralization data for elicited polyclonal antisera^13,14^ and mAbs,^18^ suggested that mosaic-8b RBD-nanoparticles represent a promising vaccine strategy to protect against current and future SARS-2 VOCs as well as additional animal sarbecoviruses that could spill over into humans.

The mosaic-8b immunized animals in our previous studies were naïve with respect to SARS-2 exposure. By contrast, the majority of humans receiving a mosaic-8b vaccine would have already been infected with SARS-2, vaccinated, or both. An important question, therefore, is whether mosaic-8b immunization would elicit broadly cross-reactive recognition of conserved sarbecovirus RBD epitopes in animals that were pre-vaccinated with spike-based COVID-19 vaccines; e.g., mRNA-lipid nanoparticle (LNP) vaccines. Specifically, we aimed to understand the role of immune imprinting, also known as original antigenic sin (OAS), as it relates to eliciting breadth in non-naïve individuals. Immune imprinting was initially postulated to explain why influenza virus infections with new and distinct viral strains resulted in preferential boosting of Ab responses against epitopes shared with the original strain.^19,20^ Recently, experiments in Ab fate-mapping (*Igk*^Tag^) mice demonstrated that, upon repeated immunization, serum Abs continue to derive from B cells that were initially activated in the primary response.^21^ Abs induced de novo in subsequent responses were suppressed, consistent with OAS. This reliance on primary cohort B cells (primary addiction) decreased with antigenic distance, resulting in partial alleviation of OAS upon boosting with BA.1 spike mRNA-LNP after single-dose WA1 mRNA priming.^21^ To investigate basic mechanisms in B cell activation in animals with previous antigenic exposure and to inform future mosaic-8b immunizations in humans with previous SARS-2 exposure, we designed experiments to compare immune responses to RBD-nanoparticles and additional COVID-19 immunizations in pre-vaccinated animals.

Here, we evaluated immune responses to mosaic-8b, mosaic-7 (mosaic-8b without a SARS-2 RBD), homotypic SARS-2, and admix-8b (mixture of 8 homotypic nanoparticles) (Figure 1A) in non-human primates (NHPs) and in mice that were previously vaccinated with DNA, mRNA, a self-amplifying RNA (repRNA), or adenovirus-vectored COVID-19 vaccines (Table S1). As previously observed in SARS-2 naïve animals,^13,14^ we found that mosaic-8b immunization in three independent pre-vaccinated animal models elicited broadly cross-reactive binding and neutralizing Ab responses against viral strains that were both matched and mismatched. As also observed in naïve animals, we found increased cross-reactivity of Abs elicited by mosaic-8b compared with homotypic SARS-2 RBD-nanoparticles in pre-vaccinated animals, and also demonstrated increased cross-reactivity for mosaic-8b versus admix-8b sera in mice that were pre-vaccinated with a Pfizer-like mRNA-LNP. Molecular fate-mapping^21^ studies in mice showed that de novo Abs were predominantly induced in pre-vaccinated mice only with mosaic-8b boosting, and that these Abs were specific for variant RBDs with increased identity to RBDs on mosaic-8b. Results from these studies contribute to increased understanding of basic immune phenomena involving B cell responses under potential OAS conditions. In addition, they support the use of a mosaic-8b RBD nanoparticle vaccine in humans with previous SARS-2 exposure to protect against future SARS-2 variants, and also of critical importance, to prevent another pandemic caused by spillover of an animal sarbecovirus with the potential for human-to-human transmission.

## RESULTS

We used the SpyCatcher-SpyTag system^22,23^ to covalently attach RBDs with C-terminal SpyTag sequences to a 60-mer nanoparticle (SpyCatcher-mi3)^24^ to make RBD-nanoparticles designated as either mosaic-8b (each nanoparticle presenting the SARS-2 Beta RBD and seven other sarbecovirus RBDs attached randomly to 60 sites),^14^ mosaic-7 (mosaic-8b without the SARS-2 Beta RBD), homotypic SARS-2 (each nanoparticle presenting 60 copies of SARS-2 Beta RBD),^14^ or admix-8b (mixture of 8 homotypic RBD-nanoparticles corresponding to the RBDs in mosaic-8b) (Figure 1; Figure S1). We used RBD-nanoparticle doses based upon RBD molarity; thus, animal cohorts received a consistent RBD dose but homotypic SARS-2 RBD-nanoparticles have 8 times as many SARS-2 Beta RBDs as either mosaic-8b or admix-8b.

Throughout the paper, we refer to COVID-19 pre-vaccinations in NHPs and mice as “vaccinations,” whereas we describe subsequent injections of additional immunogens (either RBD-nanoparticles or a genetically-encoded immunogen) in the non-naïve animals as “immunizations,” with the first injection after pre-vaccination referred to as a prime and subsequent immunizations as boosts. For each immunogen evaluated in a pre-vaccinated animal model, we assessed serum Ab binding to spike antigens (different sarbecovirus RBDs or WA1 spike) by ELISA and using an in vitro neutralization assay. Of potential relevance to results in humans, both binding and neutralizing Ab levels correlate with vaccine efficacy.^25^ We present binding and neutralization data against sarbecovirus antigens (ELISAs) or pseudoviruses (neutralization assays) for each pre-vaccinated and then immunized cohort in the form of line plots with determinations of statistically significant differences between cohorts evaluated using pairwise comparisons for each viral strain.

### Mosaic-8b immunizations in previously-vaccinated NHPs elicits cross-reactive Ab responses

To address the immunogenicity of a mosaic-8b immunization in a pre-vaccinated animal model, we immunized 14 NHPs that had previously received nucleic acid-based SARS-2 vaccines. These animals received four doses of DNA or repRNA vaccines between weeks -64 and -30 as described in the methods, Figure 2A, and Table S1.^26–28^ Eight weeks before immunization, we re-stratified the animals into three groups (n=4-5) , each of which was similar in mean and range of weights, ages, sex, and Ab neutralization titers against SARS-2 D614G (Figure S2A), without considering vaccine history. The mixed immune history in these groups of pre-vaccinated NHPs is representative of a complex immune history in people who have been vaccinated and/or infected multiple times. The NHPs were immunized at week 0 with two doses (8 weeks apart) of either mosaic-8b or homotypic SARS-2 RBD-nanoparticles or with a bivalent repRNA-LION^28^ vaccine encoding the WA1 and the Omicron BA.1 spike proteins, which mimicked immunogens used in commercial bivalent mRNA-LNP vaccines (Figure 2A). Serum was then sampled every two weeks for 22 weeks after the first nanoparticle or bivalent repRNA boost to measure the peak and subsequent contraction of serum binding and neutralizing Ab responses (Figure 2A).

**Figure 2.**
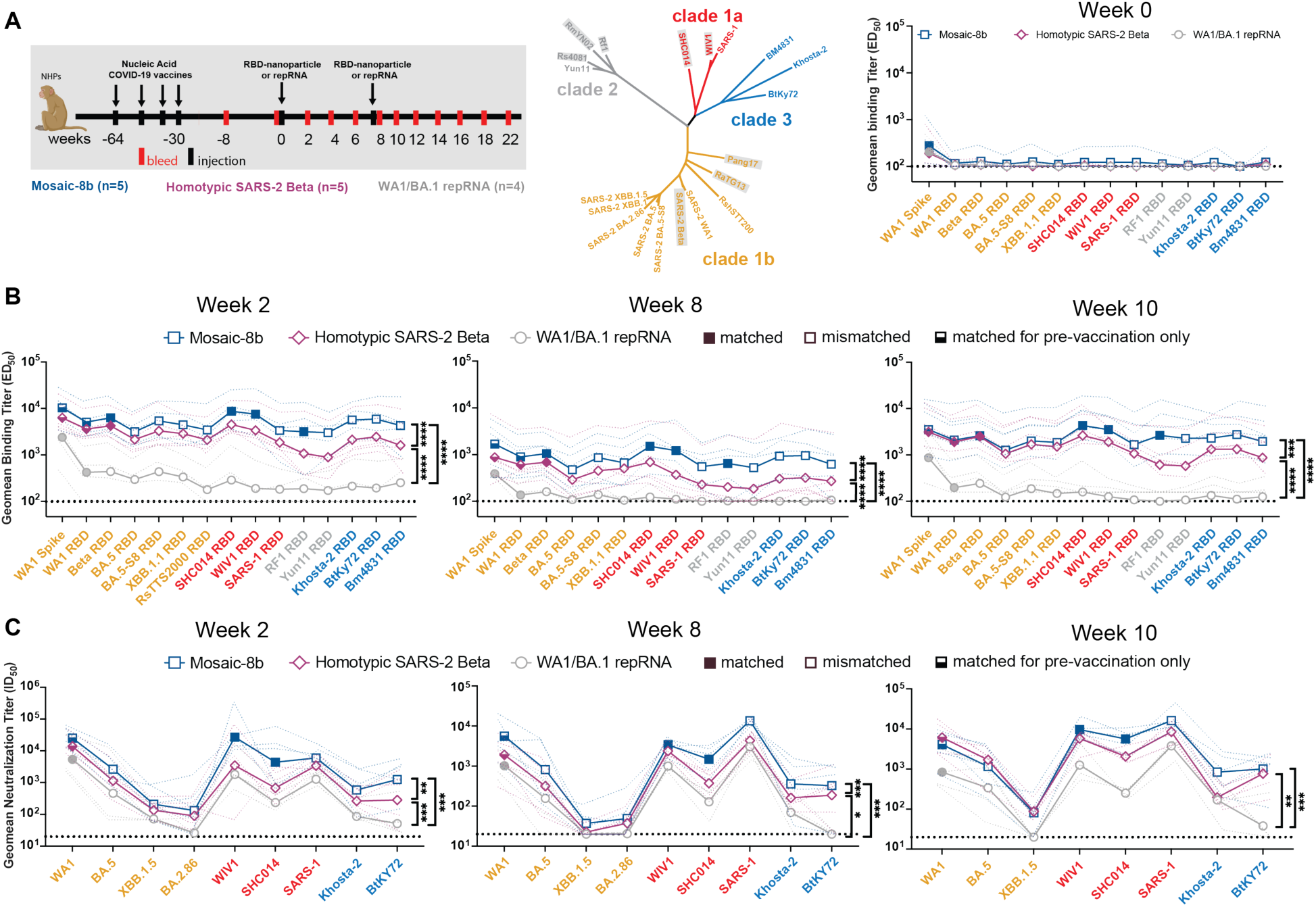
Mosaic-8b immunizations in previously-vaccinated NHPs elicits cross-reactive Ab responses, related to Figure S2. Data are shown for ELISA and neutralization analyses for serum samples from weeks 0, 2, 8, and 10. (Data for samples from weeks 4, 12, and 22 are in Figure S2.) Geometric means of ED_50_ or ID_50_ values for all animals in each cohort are indicated by symbols connected by thick colored lines; results for individual animals are indicated by dotted colored lines. Mean titers against indicated viral antigens or pseudoviruses were compared pairwise across immunization cohorts by Tukey’s multiple comparison test with the Geisser-Greenhouse correction (as calculated by GraphPad Prism). Significant differences between cohorts linked by vertical lines are indicated by asterisks: p<0.05 = *, p<0.01 = **, p<0.001 = ***, p<0.0001 = ****. (A) Left: Schematic of vaccination/immunization regimen. NHPs were vaccinated at the indicated weeks prior to RBD-nanoparticle or repRNA prime and boost immunizations at week 0 and week 8. Middle: Phylogenetic tree of selected sarbecoviruses calculated using a Jukes-Cantor generic distance model using Geneious Prime® 2023.1.2 based on amino acid sequences of RBDs aligned using Clustal Omega.^97^ RBDs included in mosaic-8b are highlighted in gray rectangles. Right: Geometric mean ELISA binding titers at week 0 (after pre-vaccinations but prior to RBD-nanoparticle or repRNA immunizations) against indicated viral antigens. (B) Geometric mean ELISA binding titers at the indicated weeks after immunization with mosaic-8b, homotypic SARS-2, or bivalent WA1/BA.1 repRNA against indicated viral antigens. (C) Geometric mean neutralization titers at the indicated weeks after immunization with mosaic-8b, homotypic SARS-2, or bivalent WA1/BA.1 repRNA against indicated sarbecovirus pseudoviruses.

We performed ELISAs against a panel of RBD and spike proteins to assess serum Ab binding titers and pseudovirus assays against available viral strains to determine Ab neutralizing titers (Figure 2; Figure S2B-C). Responses at week 0 (immediately prior to nanoparticle or repRNA-LION immunizations) showed equivalent titers across the three groups (Figure 2A). The mosaic-8b and homotypic SARS-2 nanoparticle-immunized NHPs elicited significantly higher levels of binding (Figure 2B; Figure S2B) and neutralization (Figure 2C; Figure S2C) than the bivalent repRNA-LION immunization after week 0. Notably, RBD-nanoparticle immunizations in pre-immunized NHPs elicited high overall ELISA and neutralization titers against a diverse array of both matched and mismatched sarbecoviruses (Figure 2B-C; Figure S2B-C). As generally observed for vaccines,^29–31^ we found that elicited Ab binding responses contracted after both RBD-nanoparticle and bivalent repRNA immunizations (5-8 fold from week 2 to week 8) and after boosting (∼10 fold from week 10 to week 22). Contractions in binding Ab levels largely correlated with drops in neutralizing Ab titers (Figure 2B-C; Figure S2B-C).

Mean serum binding titers for all RBD variants evaluated by ELISA were significantly higher at 5 of 6 time points evaluated after priming pre-vaccinated NHPs for mosaic-8b compared with homotypic SARS-2 samples (Figure 2B; Figure S2B). In addition, at weeks 2, 4, and 8 (after only a single nanoparticle immunization), mosaic-8b elicited significantly higher neutralization titers compared to immunization with homotypic SARS-2 (Figure 2C; Figure S2C). At every time point, either mean binding titers, mean neutralization titers, or both, were significantly higher for mosaic-8b than for homotypic samples across divergent sarbecoviruses, a required result for potentially broad protection since both Ab binding titer and neutralization correlate with vaccine efficacy in licensed SARS-2 vaccines.^31^ In summary, critical for its potential use in non-naïve humans, mosaic-8b elicited cross-reactive Ab responses even in a background of four prior COVID-19 vaccinations, and mosaic-8b immunizations consistently elicited a broader Ab response than homotypic SARS-2 and bivalent repRNA immunizations in the pre-vaccinated NHP model (Figure 2; Figure S2).

### Mosaic-8b immunizations outperform genetically-encoded vaccine and non-mosaic RBD-nanoparticle immunizations in pre-vaccinated mice

To generalize our findings in pre-vaccinated NHPs to a different animal model and to other vaccine modalities, we conducted experiments in mice that had been pre-immunized with genetically-encoded vaccines, either an mRNA-LNP formulation equivalent to Pfizer-BioNTech’s WA1 spike-encoding BNT162b2 vaccine (Figure 3; Figure S3; Table S1) or with an adenovirus-vectored vaccine (research-grade ChAdOx1 nCoV-19 produced by the Viral Vector Core Facility, Oxford University) also encoding the WA1 spike (Figure 4; Figure S4; Table S1). For mouse experiments, we could increase the number of animals per cohort and also add new cohorts (admix-8b and mosaic-7) to comparisons of mosaic-8b, homotypic SARS-2, and mRNA-LNP or viral vector immunizations after pre-vaccinations (Figure 3A,D; Figure 4A).

**Figure 3.**
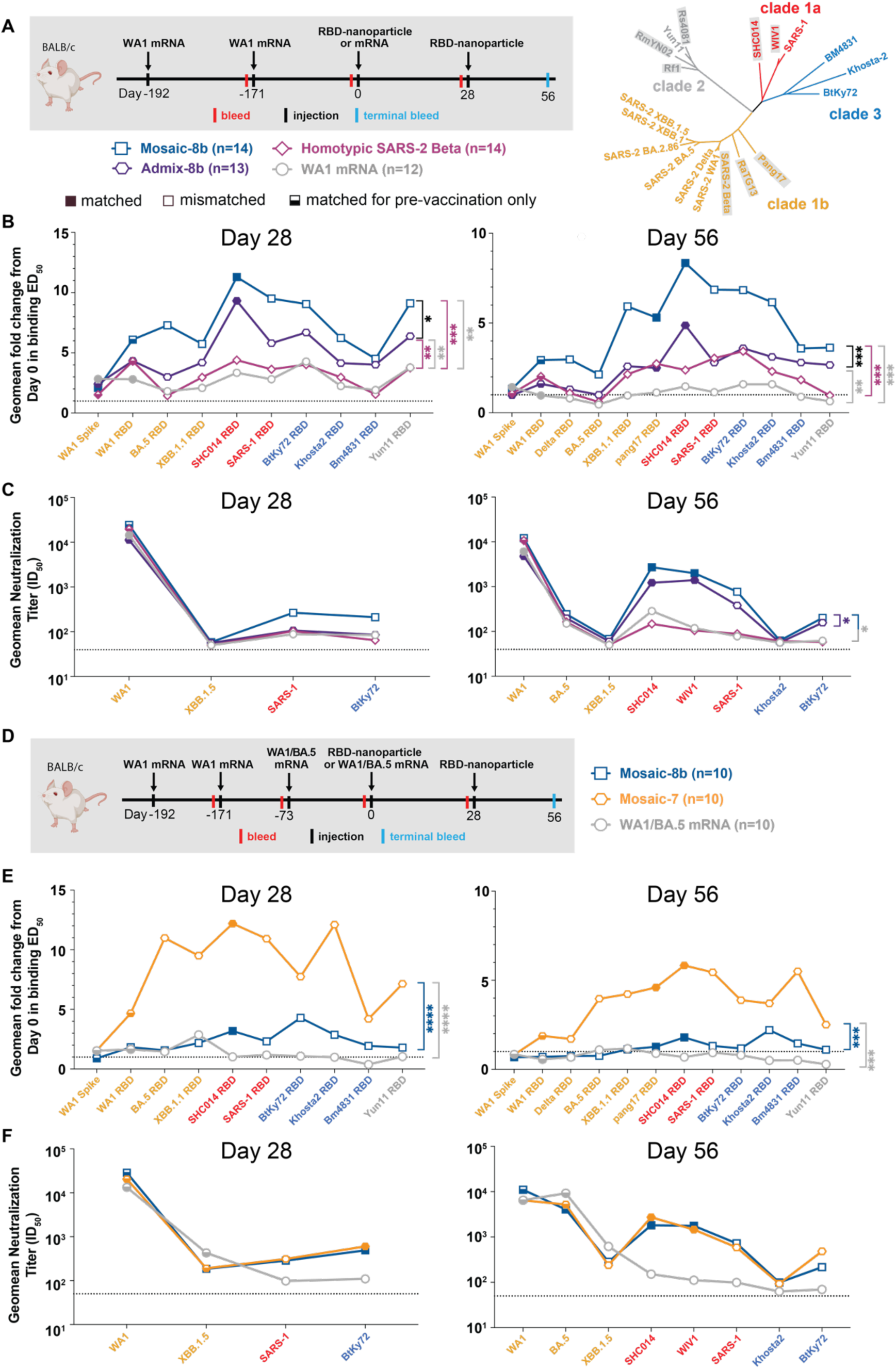
Mosaic-8b immunizations in mice previously immunized with an mRNA-LNP vaccine elicit cross-reactive Ab responses, related to Figure S3. For experiments shown schematically in panels A and D, binding responses at day 0 (immediately prior to nanoparticle or other vaccine immunizations) showed significant differences across cohorts in titers elicited by the pre-vaccinations. We therefore applied baseline corrections to account for different mean responses at day 0 prior to immunizations in each of the four groups for data in panel B and each of the three groups for data in panel E (see Methods). Non-baseline corrected binding data for panel E are shown in Figure S3G. Geometric means of fold change in ED_50_ or geometric means of ID_50_ values for all animals in each cohort are indicated by symbols connected by thick colored lines. Mean titers against indicated viral antigens or pseudoviruses were compared pairwise across immunization cohorts by Tukey’s multiple comparison test with the Geisser-Greenhouse correction (as calculated by GraphPad Prism). Significant differences between cohorts linked by vertical lines are indicated by asterisks: p<0.05 = *, p<0.01 = **, p<0.001 = ***, p<0.0001 = ****. (A) Left: Schematic of vaccination/immunization regimen for panels B and C. Mice were vaccinated at the indicated days prior to RBD-nanoparticle prime and boost immunizations at days 0 and 28 or mRNA-LNP prime immunization at day 0. Right: Phylogenetic tree of selected sarbecoviruses calculated using a Jukes-Cantor generic distance model using Geneious Prime® 2023.1.2 based on amino acid sequences of RBDs aligned using Clustal Omega.^97^ RBDs included in mosaic-8b are highlighted in gray rectangles. (B) Geometric mean fold change ELISA binding titers at the indicated days after immunization with mosaic-8b, admix-8b, homotypic SARS-2, or WA1 mRNA-LNP against indicated viral antigens. (C) Geometric mean neutralization titers at the indicated weeks after immunization with mosaic-8b, admix-8b, homotypic SARS-2, or WA1 mRNA-LNP against indicated sarbecovirus pseudoviruses. (D) Schematic of vaccination/immunization regimen for panels E and F. Mice were vaccinated at the indicated days prior to RBD-nanoparticle prime and boost immunizations at days 0 and 28 or mRNA-LNP prime immunization at day 0. (E) Geometric mean fold change ELISA binding titers at the indicated days after immunization with mosaic-8b, mosaic-7, or WA1/BA.5 mRNA-LNP against indicated viral antigens. Compare with Figure S3G(non-baseline corrected geometric mean ELISA binding titers). (F) Geometric mean neutralization titers at the indicated weeks after immunization with mosaic-8b, mosaic-7, or WA1/BA.5 mRNA-LNP against indicated sarbecovirus pseudoviruses.

**Figure 4.**
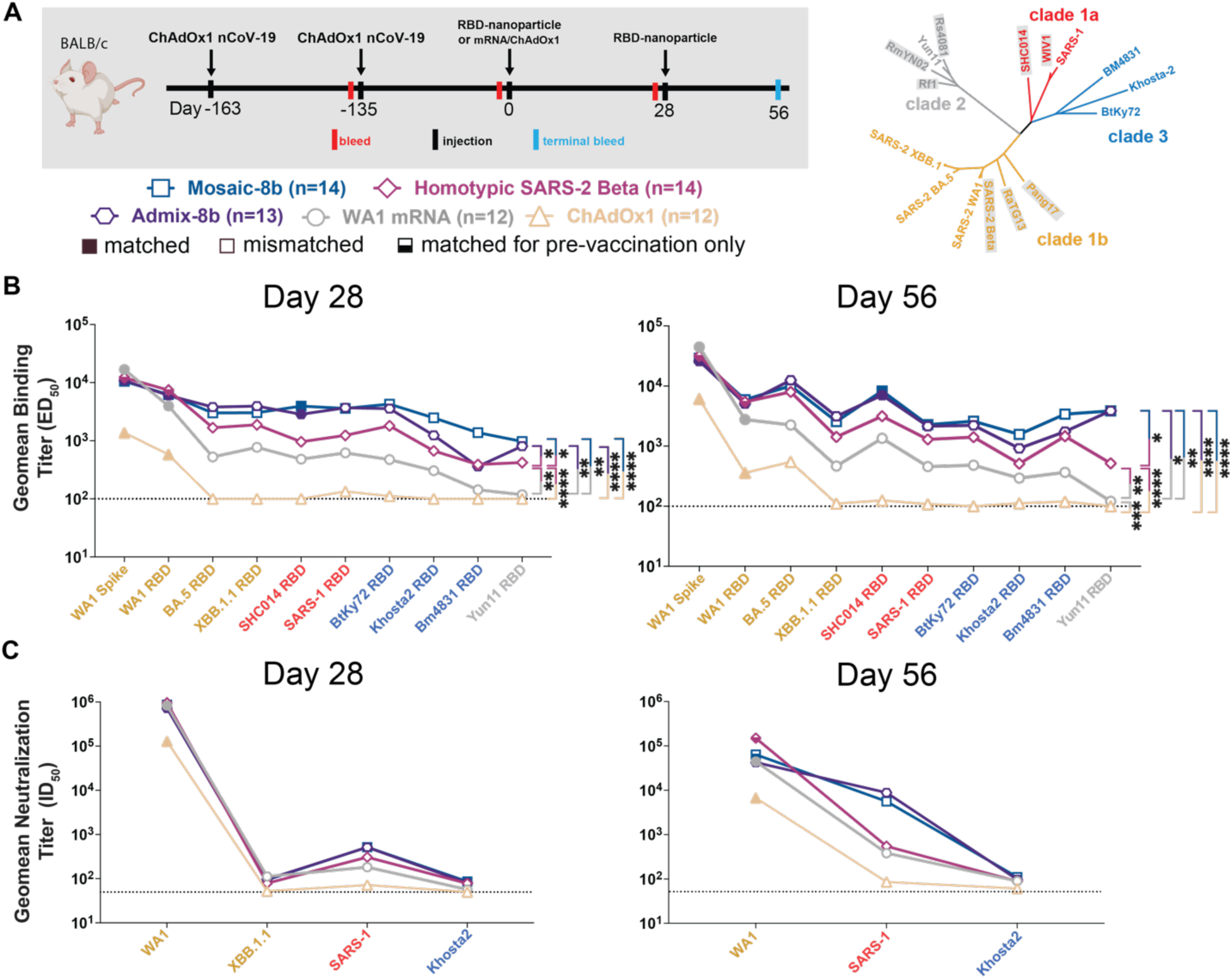
Mosaic-8b immunizations in ChAdOx1-vaccinated mice elicit cross-reactive Ab responses, related to Figure S4. Geometric means of ED_50_ or ID_50_ values for all animals in each cohort are indicated by symbols connected by thick colored lines. Mean titers against indicated viral antigens or pseudoviruses were compared pairwise across immunization cohorts by Tukey’s multiple comparison test with the Geisser-Greenhouse correction (as calculated by GraphPad Prism). Significant differences between cohorts linked by vertical lines are indicated by asterisks: p<0.05 = *, p<0.01 = **, p<0.001 = ***, p<0.0001 = ****. (A) Left: Schematic of vaccination/immunization regimen. Mice were vaccinated at the indicated days prior to RBD-nanoparticle prime and boost immunizations at days 0 and 28 or mRNA-LNP or ChAdOx1 prime immunizations at day 0. Right: Phylogenetic tree of selected sarbecoviruses calculated using a Jukes-Cantor generic distance model using Geneious Prime® 2023.1.2 based on amino acid sequences of RBDs aligned using Clustal Omega.^97^ RBDs included in mosaic-8b are highlighted in gray rectangles. (B) Geometric mean ELISA binding titers at the indicated days after immunization with mosaic-8b, admix-8b, homotypic SARS-2, WA1 mRNA-LNP, or ChAdOx1 against indicated viral antigens. (C) Geometric mean neutralization titers at the indicated weeks after immunization with mosaic-8b, admix-8b, homotypic SARS-2, WA1 mRNA-LNP, or ChAdOx1 against indicated sarbecovirus pseudoviruses.

Pre-vaccinated mice were bled prior to immunizations at day 0 and after immunizations at days 28 and 56. As in the NHP experiments, we performed ELISAs against a panel of RBD and spike proteins and pseudovirus assays against available viral strains (Figure 3, 4; Figure S3, S4). In the WA1 mRNA-LNP pre-vaccination experiments, responses at day 0 (after all animals had received the same course of mRNA-LNP vaccines) showed significant differences in binding titers elicited by the pre-vaccinations across the cohorts (Figure S3B,E), possibly because of differences in initial vaccine responses or in Ab contraction. Thus, we applied baseline corrections to account for different mean responses at day 0 in each of the four groups (see Methods). Neutralization potencies at day 0 were similar for all cohorts (Figure S3C,F) and therefore were not baseline-corrected. Baseline corrections were not required for the ChAdOx1 pre-vaccination experiments (Figure S4).

#### mRNA-LNP pre-vaccinated mice

In the WA1 Pfizer-like mRNA-LNP pre-vaccinated animals (Figure 3A; Table S1), the increase in Ab binding titers was significantly higher at days 28 and 56 for mosaic-8b than for the other cohorts when evaluated against sarbecovirus RBDs (especially notable for day 56) (Figure 3B). In general, while RBD-nanoparticle immunizations boosted Ab binding against RBDs, immunization with an additional dose of WA1 mRNA-LNP did not appreciably boost Ab responses, especially at day 56 (Figure 3B). When comparing responses to different protein nanoparticles, mosaic-8b immunizations induced significantly higher increases in mean binding titers than either admix-8b or homotypic SARS-2 immunizations. However, mean neutralization titers did not show as pronounced differences between cohorts, although the titers for mosaic-8b were higher than titers for admix-8b or homotypic SARS-2 against some viral strains (e.g., SHC014, WIV1, SARS-1) (Figure 3C).

To evaluate whether a mosaic RBD-nanoparticle without a SARS-2 RBD would also overcome potential OAS effects, we compared Ab responses raised by mosaic-7 (includes all mosaic-8b RBDs except for SARS-2; Figure 1A) to mosaic-8b responses (Figure 3D-F). Mice were pre-vaccinated with two doses of WA1 mRNA-LNP and one dose of WA1/BA.5 bivalent mRNA-LNP and then immunized with either two doses of an RBD-nanoparticle (mosaic-8b or mosaic-7) or an additional dose of WA1/BA.5 mRNA-LNP, which was included to mimic a more up-to-date immune history in the human population) (Figure 3D). We found consistently greater increases in mean Ab binding titers at day 28 and day 56 with respect to day 0 for mosaic-7 compared with mosaic-8b immunized animals across all RBDs (Figure 3E). Although increases in binding titers with respect to day 0 values were higher for mosaic-7 than for mosaic-8b, differences in mean titers for the corresponding non-baseline corrected data were only marginal between the two groups at day 28, dropping to no significance at day 56 (Figure S3G), suggesting that immunization by these RBD-nanoparticles had elicited a maximum binding anti-RBD Ab response. In addition, there were no neutralization differences for mosaic-8b and mosaic-7 antisera across strains (Figure 3F), as also observed for the non-baseline corrected mean binding titers. In this pre-vaccination/immunization regimen, both mosaic-8b and mosaic-7 were significantly better in inducing binding Abs than an additional dose of WA1/BA.5 (Figure S3G).

#### ChAdOx1 pre-vaccinated mice

We also examined the responses to mosaic-8b, homotypic SARS-2, and admix-8b in ChAdOx1 pre-vaccinated animals (Figure 4; Figure S4; Table S1). In these experiments, we compared RBD-nanoparticle immunization with immunization of an additional dose of ChAdOx1 or of WA1 mRNA-LNP (Figure 4A). Overall Ab binding responses across RBDs were generally higher for mosaic-8b and admix-8b than for homotypic SARS-2, and the three nanoparticle immunizations produced higher binding titers than an additional immunization with WA1 mRNA-LNP or ChAdOx1, as reflected in some cases with significant differences in mean ELISA binding titers (Figure 4B). Neutralization titers against SARS-1 were higher for mosaic-8b and admix-8b than for homotypic SARS-2 at day 56, but differences in mean of mean neutralization titers were not significant across cohorts (Figure 4C). Overall, mosaic-8b and admix-8b prime/boost immunizations in ChAdOx1 pre-vaccinated mice elicited the broadest binding and neutralization responses.

To determine which RBD epitopes were targeted in pre-vaccinated and then immunized mice, we performed DMS^16^ using yeast display libraries derived from WA1 and XBB.1.5 RBDs for ChAdOx1 pre-vaccinated mice that were immunized with either mosaic-8b, admix-8b, homotypic SARS-2, or WA1 mRNA-LNP (Figure 4A; Figure 5). To aid in interpreting epitope mapping of polyclonal antisera, we first conducted control experiments by comparing DMS of individual mAbs with known epitopes (class 1, 2, 3, 4, and 1/4; Figure 1B) to DMS using different mixtures of the same mAbs (Figure S5). As previously shown for mapping of mAb epitopes,^32,33^ DMS revealed the expected escape profiles matching the epitopes of the control mAbs (Figure S5). However, when mAbs were mixed in various ratios, DMS signals were obscured, with little or no highlighting of epitopes in the equimolar mix and only partial highlighting when the mixtures contained either more class 3, class 1/4, and class 4 mAbs or more class 1/class 2 mAbs (Figure S5). Thus, a polyclonal serum including anti-RBD Abs that are equally distributed across all classes (i.e., a “polyclass” antiserum) will likely display low DMS escape fractions across all RBD residues, whereas a polyclonal serum response that is more focused on particular epitope(s) will show more defined escape peaks.

**Figure 5.**
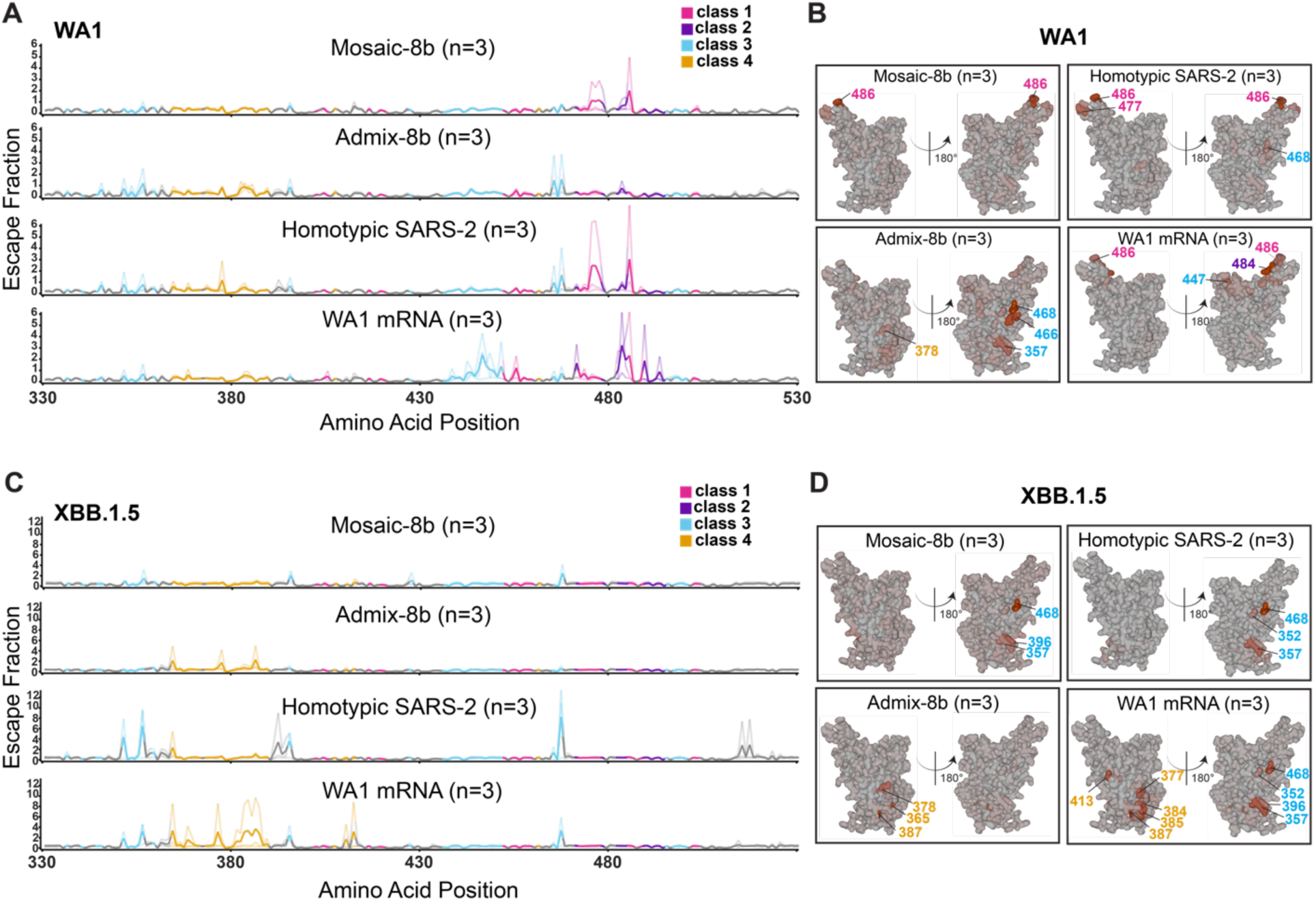
Differences in epitope targeting of Abs elicited in ChAdOx1 pre-vaccinated mice by mosaic-8b and admix-8b compared with homotypic SARS-2 and WA1 mRNA-LNP, related to Figure S5. Sera were derived from ChAdOx1 pre-vaccinated mice that were immunized with either two doses of an RBD-nanoparticle immunogen or one dose of WA1 mRNA-LNP. (A) Line plots for DMS results from sera from ChAdOx1-vaccinated mice that were immunized with the indicated immunogens (immunization schedule in Figure 4A). DMS was conducted using a WA1 RBD library. The x-axis shows the RBD residue number and the y-axis shows the sum of the Ab escape of all mutations at a site (larger numbers indicating more Ab escape). Each line represents one antiserum with heavy lines showing the average across the n=3 sera in each group. Lines are colored differently for RBD epitopes within different classes^17^ (color definitions in upper right legend; epitopes defined in Figure 1E; gray for residues not assigned to an epitope). (B) The average site-total Ab escape for the WA1 RBD library for mice immunized with the indicated immunogens after ChAdOx1 vaccination mapped to the surface of the WA1 RBD (PDB 6M0J), with gray indicating no escape, and epitope-specific colors indicating sites with the most escape. (C) Line plots for DMS results from sera from ChAdOx1-vaccinated mice that were immunized with the indicated immunogens (immunization schedule in Figure 4A). DMS was conducted using a XBB.1.5 RBD library. The x-axis shows the RBD residue number and the y-axis shows the sum of the Ab escape of all mutations at a site (larger numbers indicating more Ab escape). Each line represents one antiserum with heavy lines showing the average across the n=3 sera in each group. Lines are colored differently for RBD epitopes within different classes^17^ (color definitions in upper right legend; epitopes defined in Figure 1E; gray for residues not assigned to an epitope). (D) The average site-total Ab escape for the XBB.1.5 RBD library for mice immunized with the indicated immunogens after ChAdOx1 vaccination mapped to the surface of the WA1 RBD (PDB 6M0J), with gray indicating no escape, and epitope-specific colors indicating sites with the most escape.

We next analyzed differences in epitope targeting of Abs elicited by mosaic-8b, admix-8b, homotypic SARS-2, and WA1 mRNA-LNP in ChAdOx1 pre-vaccinated mice (Figure 5), keeping in mind (*i*) the possibility of inducing polyclass Abs, and (*ii*) that DMS results can depend on the particular RBD library used in an experiment. For example, class 1 and class 2 anti-RBD Abs from WA1 mRNA-LNP vaccinated animals do not bind to XBB.1.5, which would emphasize detection of responses to the more conserved class 3 and class 4 epitopes.

We found that both homotypic SARS-2 and WA1 mRNA-LNP immunizations resulted in relatively defined class 1, 2, and 3 DMS profiles against the WA1 library (Figure 5A-B) and defined class 3 and 4 profiles against the XBB.1.5 library (Figure 5C-D). By contrast, the mosaic-8b and admix-8b DMS profiles were consistent with polyclass Ab induction (Figure 5A-D). Although sometimes difficult to discern, the majority of responses against the WA1 library mapped to class 1, 2, or 3 epitopes (Figure 5B), whereas most responses against the XBB.1.5 library mapped to class 3 or class 4 epitopes (Figure 5D). We conclude that immunizing pre-vaccinated ChAdOx1 mice with a mosaic or admix RBD-nanoparticle immunogen induces a more polyclass Ab response than immunization with a homotypic protein or mRNA-LNP immunogen. In addition, we note that serum boosting responses after mosaic-8b and admix-8b immunizations in the pre-vaccinated mice showed only weak signals for the more conserved class 3 and class 4 epitopes (Figure 5A,B), similar to results for the mAb mix 3 experiment, in which class 1 and class 2 mAbs were in excess over other mAbs (Figure S5). These results imply a diversified response to RBDs after mosaic or admix immunization; i.e., class 1 and class 2 anti-RBD Abs originally induced by ChAdOx1 vaccination dominated serum responses and subsequently masked DMS signals from class 3 and class 4 Abs induced by mosaic-8b and admix-8b.

### Molecular fate-mapping of mosaic-8b boosting in previously vaccinated mice reveals efficient recall of cross-reactive Abs and generation of strain-specific de novo responses

DMS comparisons in pre-vaccinated mice (Figure 5; Figure S5) involved analyses of mixtures of de novo and recall Ab responses. Thus, we could not determine whether the improved breadth elicited by mosaic-8b immunization was due to de novo responses or recall of cross-reactive Abs elicited by the primary vaccination series. To address this issue, we used recently developed molecular fate-mapping S1pr2-Cre^ERT2^*.Igk*^Tag/Tag^ (S1pr2-*Igk*^Tag^) mice that enable antigen-specific serum Abs to be traced to their cellular and temporal origins.^21^ In these mice, the C-terminus of the immunoglobulin kappa light chain (Igκ) is extended to encode a LoxP-flanked Flag tag and stop codon followed by a downstream Strep tag. B cells bearing this *Igk*^Tag^ allele produce Abs that are Flag-tagged unless they are exposed to Cre recombinase, after which they permanently switch the Flag tag for a Strep tag.^21^ In S1pr2-*Igk*^Tag^ mice, tamoxifen-inducible Cre is expressed in B cells as they undergo affinity maturation in germinal centers (GCs).^34^ Abs derived from the cohort of B cells engaged in GCs at the time of tamoxifen administration, including the cohort’s memory B cell and plasma cell progeny, will be fate-mapped as Strep^+^. Boost-induced de novo GCs that are mainly formed by new cohorts of naïve B cells^35^ remain Flag^+^ in the absence of tamoxifen administration, and therefore de novo Abs derived from these B cells are reverse fate-mapped with Flag.^21^ Thus, by using tamoxifen to initiate fate-mapping of the B cells and Abs elicited by primary mRNA-LNP vaccinations, the Flag^+^ Abs induced de novo by RBD-nanoparticle immunizations can be distinguished from the recalled primary-cohort derived Strep^+^ Abs (Figure 6; Figure S6, Figure S7).

**Figure 6.**
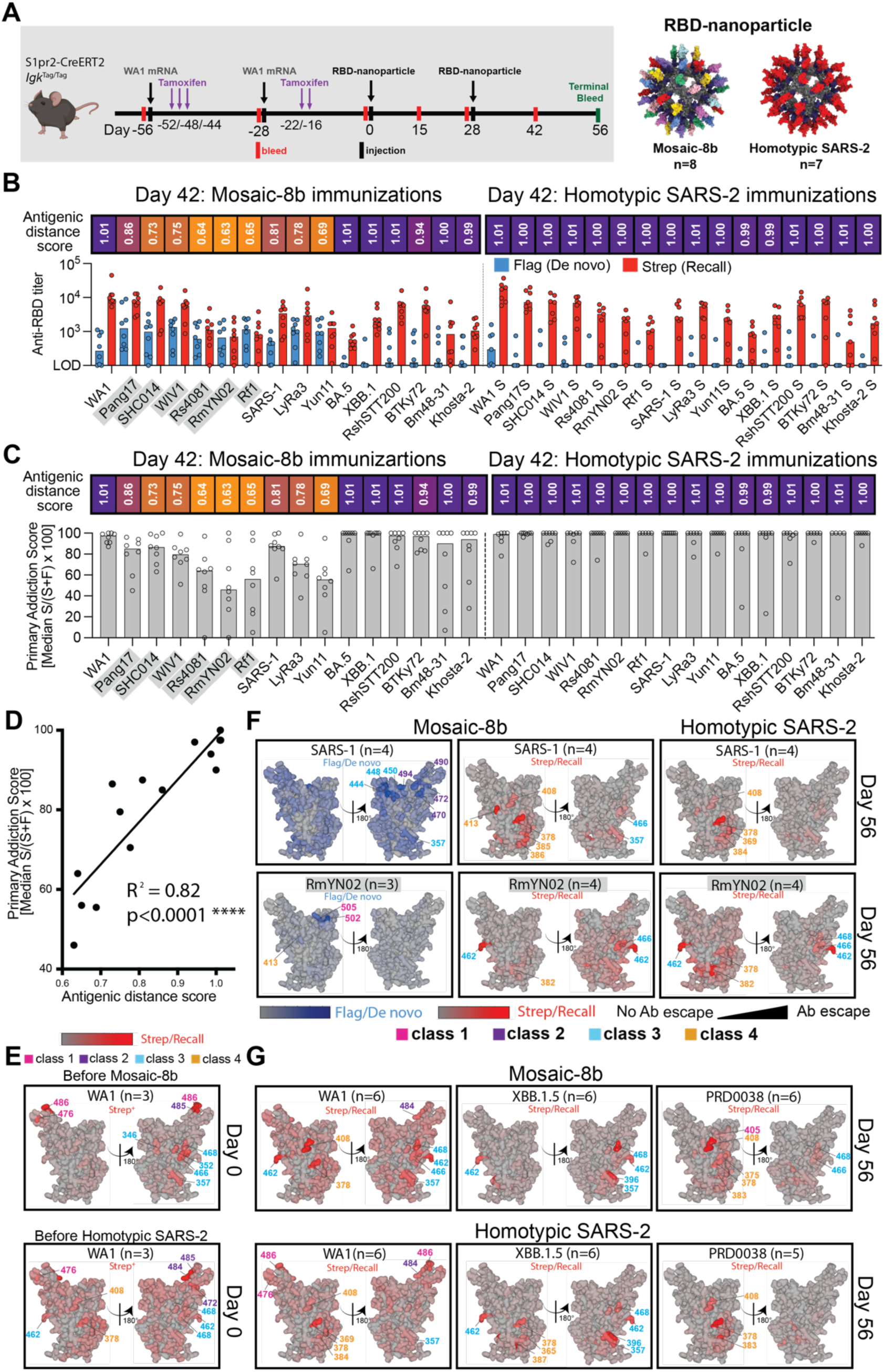
Fate mapping of serum antibodies reveals elicitation of cross-reactive recall responses and strain-specific de novo responses elicited by mosaic-8b nanoparticles in previously vaccinated animals, related to Figure S6. Antigenic distance score is defined as the ratio of % amino acid sequence identity of a strain to WA1 RBD divided by % sequence identity of that strain to the closest non-self RBD relative on the nanoparticle. The primary addiction index^21^ is defined as Strep/(Flag + Strep) × 100. See Figure 1B for RBD epitope classifications. (A) (Left) Schematic of vaccine regimen in S1pr2-*Igk*^Tag/Tag^ immunized mice. Mice were vaccinated with an mRNA-LNP vaccine encoding WA1 spike at the indicated days prior to RBD-nanoparticle immunizations at day 0 and day 28. After each dose of mRNA-LNP vaccine, mice were treated with tamoxifen to switch epitope tags on the Abs produced by B cells activated by the mRNA-LNP vaccinations from Flag to Strep, so that Ab responses elicited by RBD-nanoparticle boosting can be separated into recall (Strep^+^) and de novo (Flag^+^) responses. (Right) Models of RBD-nanoparticles that were used for immunizations at days 0 and 28. (B) Comparison of the Flag^+^/de novo (blue) and Strep^+^/recall (red) anti-RBD titers shown at day 42, two weeks after a second immunization with mosaic-8b or homotypic SARS-2 nanoparticles. For each strain, the antigenic distance score is shown above the plots and colored with a gradient from purple (score=1) to yellow (score=0). Gray shaded strains are matched to mosaic-8b. (C) The primary addiction index^21^ for each RBD is plotted for day 42 serum after two doses of mosaic-8b or homotypic SARS-2. The antigenic distance score is shown above the plots and colored with a gradient from purple (score=1) to yellow (score=0). Gray shaded strains are matched to mosaic-8b. (D) Correlation of the primary addiction index^21^ with the antigenic distance score. The best-fit line, R^2^, and p value were calculated by linear regression using Prism. (E) DMS analysis of Strep^+^ Abs from day 0 (prior to RBD-nanoparticle immunization) serum from n=3 pre-vaccinated mice using an RBD mutant library derived from WA1. Ab binding sites are shaded red according to degree of Strep^+^ Ab escape on the surface of the WA1 RBD (PDB 6M0J). Locations of residues with high escape scores are indicated on RBD surfaces with gray indicating no escape and shades of red indicating sites with most escape. Residue numbers are color coded according to their RBD epitope classification. (F) DMS analysis using SARS-1 (antigenic distance score = 0.81) and RmYN02 (antigenic distance score = 0.63) RBD mutant libraries of day 56 serum from n=4 or n=3 RBD-nanoparticle immunized mice. Ab binding sites are shaded according to degree of Ab escape, with blue for Flag/de novo responses and red for Strep/recall responses, on the surface of the WA1 RBD (PDB 6M0J). Comparisons are made for Flag/de novo and Strep/recall elicited by mosaic-8b and for Strep/recall elicited by homotypic SARS-2 (Flag/de novo responses after homotypic SARS-2 immunization were weak to undetectable). Gray shaded virus names represent strains that are matched to mosaic-8b. Locations of residues with high escape scores are indicated on RBD surfaces with gray indicating no escape and darker shades indicating sites with most escape. Residue numbers are color coded according to their RBD epitope classification. (G) DMS analysis of the Strep/recall compartment of day 56 serum from n=5 or n=6 mosaic-8b or homotypic SARS-2 immunized mice using RBD mutant libraries derived from strains with antigenic distance scores of 1.01 (WA1), 1.01 (XBB.1.5), and 0.94 (PRD008). Ab binding sites are shaded red according to degree of Strep/Recall Ab escape on the surface of the WA1 RBD (PDB 6M0J). Locations of residues with high escape scores are indicated on RBD surfaces with gray indicating no escape and darker shades indicating sites with most escape. Residue numbers are color coded according to their RBD epitope classification.

S1pr2-*Igk*^Tag^ mice were vaccinated twice with WA1 spike mRNA-LNP and then immunized twice with either mosaic-8b or homotypic RBD-nanoparticles (Figure 6A; Table S1). We analyzed binding of total IgG, Strep^+^ (recall), or Flag^+^ (de novo) Abs as a function of time after vaccination and immunization (Figure S6), obtaining data for the pre-vaccination timeframe that were consistent with published results.^21^ Based on these results, we focused on day 42 serum (two weeks after final immunization) for assessment of Flag/Strep Ab reactivity against a panel of RBDs. While both groups of mice recalled cross-reactive Strep^+^ Abs derived from primary responses, only the mosaic-8b boosted mice induced de novo Flag^+^ Abs (Figure 6B). The dominance of primary-derived Abs against most RBDs in both groups (Figure 6C) confirmed the strong OAS-like “primary addiction” previously observed with homologous mRNA-LNP boosting.^21^ In mRNA-LNP pre-vaccinated mosaic-8b immunized mice, de novo Ab reactivity was strongest against RBDs antigenically divergent from the priming WA1 RBD and from RBDs that were matched or antigenically highly similar to those on mosaic-8b: the lower the antigenic distance score (ratio between % amino acid identity to WA1 divided by the % identity to the most closely related non-self RBD on the immunizing RBD-nanoparticle), the higher the de novo Flag Ab response against the tested RBD (Figure 6B-C). This resulted in a lower reliance on primary cohort B cells, as evaluated by the primary addiction score (Strep/(Flag+Strep)*100) (Figure 6C), which correlated with the antigenic distance score (R^2^ = 0.82, p<0.0001; Figure 6D). Thus, we find that RBD-nanoparticle immunizations in pre-vaccinated mice elicited mostly cross-reactive recall responses first induced by mRNA-LNP vaccination, while subsequent immunization with mosaic-8b additionally induced de novo Ab responses against RBDs closely related to those present on the nanoparticle.

To determine the epitope specificity of de novo versus recall Abs in pre-vaccinated mice that were immunized with mosaic-8b, we performed DMS^16^ using libraries derived from WA1, XBB.1.5, SARS-1, RmYN02, and PRD-0038 RBDs (Figure 6E-G; Figure S7). As expected, day 0 responses against the WA1 library revealed a predominant class 1/class 2 response (Figure 6E). We compared Flag^+^/de novo and Strep^+^/recall Ab epitopes at day 56 mapped onto the SARS-1 and RmYN02 RBDs (Figure 6F): For SARS-1, we found that de novo responses to mosaic-8b immunizations after WA1 mRNA-LNP pre-vaccination targeted more variable portions of class 2 and class 3 RBD epitopes, whereas recall responses targeted the more conserved class 4 epitope and conserved portions of the class 3 epitope (Figure 6F; Figure S7A). For RmYN02, de novo responses were mainly class 1, and recall responses were class 3 and class 4 (Figure 6F; Figure S7B). In each individual mouse, de novo Flag^+^ Abs targeted epitopes that were distinct from those targeted by primary-derived Abs (Figure S7), as observed previously for WA1 priming followed by BA.1 boosting.^21^ The Strep/recall response elicited by homotypic SARS-2 targeted similar conserved epitopes on SARS-1, RmYN02, XBB.1.5 and PRD-0038 as responses to mosaic-8b (Figure 6F-G), as expected since Abs that bind to SARS-1, RmYN02, XBB.1.5 or PRD-0038 are necessarily cross-reactive. Accordingly, when comparing responses against WA1, mosaic-8b elicited a more conserved class 3/class 4 response, whereas homotypic SARS-2 elicited primarily class 1 responses combined with a minority class 4 responses, suggesting that mosaic-8b preferentially boosted cross-reactive recall Abs. These results are consistent with Ab-mediated feedback in which de novo Ab responses form against available epitopes that are not blocked by pre-existing high affinity Abs^21,36–39^ and with a model in which primary vaccine-derived Abs shift recall GC B cells away from previously targeted sites, thereby directing de novo Ab responses specifically towards other epitopes.^40^

## DISCUSSION

Much of the human population has already mounted immune responses to SARS-2, either from vaccination, infection, or both.^41,42^ Thus, new vaccines, especially pan-sarbecovirus vaccines designed to protect from future zoonotic spillovers and from SARS-2 VOCs, should be evaluated in non-naïve animal models. Here, we demonstrate that the advantages of mosaic-8b RBD-nanoparticles in eliciting cross-reactive and broad Ab responses in naïve animals^13,14^ are maintained in experiments in animals previously vaccinated with two to four doses of multiple types of COVID-19 vaccines: DNA or repRNA vaccines in NHPs, a Pfizer-equivalent mRNA-LNP in mice, or adenovirus-vectored AstraZeneca ChAdOx1 in mice.

We observed broadly cross-reactive Ab responses, as assessed by binding and neutralization assays, for mosaic-8b immunizations in pre-vaccinated animals, an important result because both binding and neutralization correlate with protection conferred by genetically-encoded and protein-based vaccines in humans and animals.^25,31,43–45^ We found that RBD-nanoparticle immunizations (mosaic-8b, admix-8b, mosaic-7, or homotypic SARS-2) produced higher binding and neutralizing titers than additional booster immunizations with mRNA-LNP, repRNA-LION, or adenovirus-vectored vaccines in pre-vaccinated mice or NHPs. When comparing RBD-nanoparticle immunogens, we observed higher homologous and heterologous neutralization titers in NHPs after only one dose of mosaic-8b versus one dose of homotypic SARS-2; the latter being a more typical protein-based COVID-19 vaccine akin to other homotypic RBD- and spike-based nanoparticle vaccines.^46–57^ This result suggests that a single mosaic RBD-nanoparticle immunization could be more broadly protective in non-immunologically naïve humans than one dose of a SARS-2 homotypic vaccine. We also note that our pre-vaccination experimental setups mimicked likely vaccination scenarios in humans; e.g., in which mosaic-8b vaccinees would have received one or more COVID-19 vaccination(s) months to year(s) before a mosaic-8b immunization (as in our mouse and NHP experiments in which mosaic-8b was injected 0.4–1.2 years after COVID-19 vaccinations) and may have also previously received different vaccine modalities. In summary, our results in three pre-vaccinated animal models, taken together and individually, support the premise that a mosaic-8b vaccine would elicit broad Ab responses in humans.

In a previous study in naïve mice,^14^ we used DMS to show that immunizations of mosaic-8b and homotypic SARS-2 elicited different types of Abs: for mosaic-8b, mainly Abs against the more conserved class 3, class 4, and class 1/4 RBD epitopes that exhibit increased cross-reactivity across sarbecoviruses and SARS-2 VOCs,^58–63^ versus for homotypic SARS-2, mostly class 1 and class 2 epitopes that rapidly evolve and vary between zoonotic and human sarbecoviruses.^32,64,65^ Here, DMS analyses of recall anti-RBD Abs from molecular fate-mapping studies demonstrated that mosaic-8b immunization boosted class 3 and 4 anti-RBD Abs originally elicited by primary vaccination with mRNA-LNP. Since analogous Abs have been identified in human donors,^58,65–71^ our DMS analyses of elicited Abs suggest that mosaic-8b vaccination in humans would boost cross-reactive Abs that broadly recognize mismatched zoonotic sarbecoviruses, including more distantly related clade 3 sarbecoviruses that could initiate another pandemic, and SARS-2 VOCs. Critical for the efficacy of mosaic-8b vaccination in humans, boosted responses in pre-vaccinated mice mirrored the responses of mosaic-8b in naïve mice.^14^

Molecular fate-mapping^21^ allowed us to separately characterize de novo and recall Ab responses to RBD-nanoparticles after mRNA-LNP prime and boosting vaccinations, demonstrating that Abs were derived mostly from B cells that were first engaged by mRNA-LNP vaccination (an example of primary addiction^21^), and that mosaic-8b, but not homotypic SARS-2, induced substantial de novo Ab responses. In general, the recall Abs were broader and more cross-reactive than the more strain-specific de novo responses. We infer that Abs induced de novo after mosaic-8b immunization in pre-vaccinated mice targeted epitopes that were not blocked by mRNA vaccine-induced Abs, supporting a model in which feedback from originally induced Abs suppresses induction of de novo Abs against overlapping epitopes.^21,37–40^ Our results also suggest that immunizing with mosaic-8b after COVID-19 vaccination elicits de novo strain-specific Abs that recognize largely variable epitopes on non-SARS-2 strains that are completely or closely matched to RBDs on mosaic-8b. Indeed, eliciting Abs against the seven non-SARS-2 RBDs on mosaic-8b could provide additional protection from future zoonotic spillovers of sarbecoviruses that are closely related to those represented by RBDs on mosaic-8b. Thus, it could be advantageous to introduce a clade 3 RBD to a mosaic RBD-nanoparticle to promote further cross-reactive recognition. This could be achieved by replacing the SARS-2 Beta RBD on mosaic-8b with a clade 3 RBD since we found improved Ab responses for mosaic-7, a mosaic RBD-nanoparticle without SARS-2 Beta, compared with mosaic-8b, which includes SARS-2 Beta. Additional RBD compositions on a mosaic RBD-nanoparticle could also be explored: for example, recent results show that mosaic-7_COM_ (computationally chosen from natural sarbecoviruses) outperformed mosaic-8b and mosaic-7 in both naïve and pre-vaccinated animals.^72^

In addition to informing future mosaic RBD vaccine design, our results are relevant to understanding OAS, first described with reference to influenza infections and more recently extended to studies of SARS-CoV-2 and other viruses.^19,20,73,74^ Our studies showed that OAS did not entirely prevent mosaic-8b’s induction of potent and cross-reactive Abs after vaccination with monovalent or bivalent genetically-encoded vaccines in three independent pre-vaccinated animal models, reproducing our findings of mosaic-8b advantages over homotypic SARS-2 immunogens in naïve animals.^13,14^ In our pre-vaccinated animals, we observed that RBD-nanoparticle immunizations mainly produced recall Abs but that de novo responses could be elicited by including antigenically-distant antigens, consistent with previous molecular fate-mapping studies^21^ and the strong correlation (R^2^ = 0.82; p<0.0001) observed in the current study between primary addiction and antigenic distance scores. This is relevant to use of mosaic-8b as a vaccine because it includes RBDs that are both closely (∼91% sequence identity) and more distantly (62%) related to the WA1 RBD in spike-containing vaccinations; thus, we could use OAS to our advantage to both boost recall responses and to elicit de novo responses. By contrast, when immunizing pre-vaccinated animals with homotypic SARS-2, only recall responses were elicited. Taken together, our results are consistent with the general premise that the immune system evolved to respond to new pathogens.

We also note that recall Abs that were boosted by mosaic-8b after monovalent or bivalent SARS-2 spike immunizations should include broadly cross-reactive Abs, as demonstrated by the isolation of potent and broadly cross-reactive monoclonal Abs from human convalescent COVID-19 donors and vaccinees despite the immunodominance of the more strain-specific class 1 and class 2 anti-RBD Abs.^58,65–71^ Thus, mosaic-8b immunization in, e.g., an mRNA-LNP vaccinated person, would prepare that person’s humoral immune system for a novel sarbecovirus, akin to the idea of using an influenza vaccine encoding 20 different hemagglutinins to prime humans for the next influenza epidemic.^75^ Finally, mosaic-8b immunization of the non-naïve human population would produce de novo responses to the 7 other RBDs that could then be recalled or boosted after an infection by a zoonotic sarbecovirus in a new spillover or by a novel SARS-2 VOC. Taken together, our results support a model in which OAS influences responses to immunogens but that the immune system can also make new responses. Thus, we advocate using maximally multivalent sarbecovirus next-generation vaccines (i.e., with as much antigenic distance as possible) that do not require updating in a single immunization rather than continuously updating with new monovalent or bivalent vaccines.

### Limitations of the study

We performed three independent experiments in pre-vaccinated animal models to compensate for not being able to compare responses to RBD-nanoparticle immunogens in vaccinated versus non-vaccinated humans. We used different types of vaccines encoding SARS-2 spike to pre-vaccinate animals: DNA-based, mRNA-LNP, and an adenovirus-vectored vaccine. However, time and financial limitations prevented evaluating other types of COVID-19 vaccines; e.g., a protein-based vaccine such as Novavax,^45^ in animal models. Because we were unable to obtain licensed Pfizer-BioNTech or Moderna mRNA-LNP vaccines for research purposes from the respective companies, we used mRNA-LNP formulations from other sources that may not perform identically to clinically-available vaccines. In addition, potential improvements to mosaic-8b (e.g., mosaic-7 or a mosaic-8 with clade 3 RBD instead of SARS-2 RBD) could not be evaluated in the time frame of these experiments; nor could we include more recent SARS-2 VOCs in assays due to limited remaining quantities of immune serum. We note that our assays to compare immune responses included serum Ab binding and neutralization but not protection assessment in a challenge model. We believe this concern is mitigated by our previous demonstration of mosaic-8b protection from matched and mismatched challenges in K18-hACE2 transgenic mice,^14^ a stringent model of coronavirus infection,^15^ and by the observation that protection has been widely reported for most, if not all, COVID-19 vaccine candidates in less stringent animal challenge models. A K18-hACE2 challenge experiment involving a mismatched sarbecovirus in pre-vaccinated and then immunized animals could be informative with respect to OAS effects but was outside the scope of this study. Finally, it is not possible to find analogous doses for genetically-encoded vaccines versus protein-based vaccines; thus, it is hard to interpret differences in elicited Ab quantities and neutralization potencies when comparing the effects of boosting vaccinated animals with an mRNA-LNP or adenovirus-vectored vaccine to boosting with RBD-nanoparticles.

Future studies of interest in pre-vaccinated animals immunized with mosaic-8b include isolating monoclonal Abs and investigating how the memory B cell compartment is altered.

## ACKNOWLEDGEMENTS

We thank Jesse Bloom (Fred Hutchinson) and Tyler Starr (University of Utah) for RBD libraries and help setting up DMS at Caltech, Jost Vielmetter, Luisa Segovia, Annie Lam, and the Caltech Beckman Institute Protein Expression Center for protein production, Igor Antoshechkin and the Caltech Millard and Muriel Jacobs Genetics and Genomics Laboratory for Illumina sequencing, S. Alison Arnold, Connor Bowen, Sharon Versteeg, Jack E. Kay, Joseph Moores-Poole, and Vasilis Tsakonitis (Ingenza, Ltd.) for expressing and purifying Bacillus mi3 and *Pichia* RBDs, Marta Murphy for help making figures, Kenneth Matreyek (Case Western Reserve University) for the high-expressing hACE2 cell line used for SHC014 neutralization assays, the Viral Vector Core Facility (Oxford University) for ChAdOx1 nCoV-19, Helix Biotech for Pfizer-like WA1 and BA.5 mRNA-LNP, Jeffrey Ravetch (Rockefeller University) for the Moderna WA1 mRNA-LNP vaccine, Labcorp Drug Development (Denver, PA) for performing BALB/c mouse immunizations for the mRNA-LNP and ChAdOx1 nCoV-19 experiments, T. Kurosaki and T. Okada (U. Osaka, RIKEN-Yokohama) for S1pr2-Cre^ERT2^ BAC-transgenic mice, and Alain Townsend and Jack T. K. Tan (Oxford), Andrew DeLaitsch and Magnus Hoffmann (Caltech), and Bjorkman lab members for helpful discussions. These studies were funded by Wellcome Leap (P.J.B.), the National Institutes of Health: [P01-AI165075 (P.J.B.), R01AI119006 and R01AI173086 (G.D.V.), R01AI153064 (N.P.)], Bill and Melinda Gates Foundation INV-034638 (P.J.B.), the Coalition for Epidemic Preparedness Innovations (CEPI) (P.J.B., I.F.), the Merkin Institute for Translational Research (Caltech), and the Stavros Niarchos Foundation Institute (Rockefeller).

## AUTHOR CONTRIBUTIONS

Conceptualization: A.A.C., J.R.K., D.H.F., P.J.B.; Methodology: A.A.C., J.R.K., A.S., S.E.D., A.J.G., D.H.F., G.D.V., P.J.B.; Software, A.J.G., A.P.W.; Investigation: A.A.C., J.R.K., A.S., S.E.D., A.V.R., H.G., P.N.P.G., Resources: A.I.R., J.H.E., J.D.P., H.M., N.P., P.J.C.L., R.C., C.S.-, E.R., S.B., M.Q.-A.; Writing – original draft: A.A.C., J.R.K., A.S., P.J.B.; Writing – review and editing: A.A.C., J.R.K., A.C., A.P.W., S.E.D., D.H.F., G.D.V., P.J.B.; Visualization: A.A.C., J.R.K., A.S., C.F., S.E.D.; Supervision: J.R.K., L.M., I.G.F., G.D.V., D.H.F., P.J.B.; Project Administration: J.R.K., L.M., I.G.F., D.H.F., P.J.B.; Funding: I.G.F., G.D.V., P.J.B.

## INCLUSION AND DIVERSITY

We support inclusive, diverse, and equitable conduct of research.

## STAR METHODS

### RESOURCE AVAILABILITY

#### Lead contact

Further information and requests for resources and reagents should be directed to and will be fulfilled by the lead contact: Pamela J. Bjorkman.

#### Materials availability

All materials generated in this study are available upon request through a Materials Transfer Agreement.

#### Data and code availability

Data are available in the main text, the supplementary information, and ELISA and neutralization assay raw data will be publicly available. DMS data are deposited on Github (https://github.com/bjorkmanlab/Mosaic-8b_prevax_fate_mapping) and raw sequencing data are available under NCBI SRA: NCBI SRA: Bioproject: PRJNA1067836 and Biosamples: SAMN39552627 and SAMN40463706. Materials are available upon request to the corresponding author with a signed material transfer agreement. Any additional information required to reanalyze the data reported in this paper is available from the lead contact upon request. This paper does not report original code. This work is licensed under a Creative Commons Attribution 4.0 International (CC BY 4.0) license, which permits unrestricted use, distribution, and reproduction in any medium, provided the original work is properly cited. To view a copy of this license, visit https://creativecommons.org/licenses/by/4.0/. This license does not apply to figures/photos/artwork or other content included in the article that is credited to a third party; obtain authorization from the rights holder before using such material.

### Key resources table

| REAGENT or RESOURCE | SOURCE | IDENTIFIER |
| --- | --- | --- |
| Antibodies |  |  |
| C102 IgG | Robbiani et al.<sup>70</sup> |  |
| C144 IgG | Robbiani et al.<sup>70</sup> |  |
| S309 IgG | Pinto et al.<sup>60</sup> |  |
| CR3022 IgG | Ter Meulen et al.<sup>76</sup> |  |
| C118 IgG | Robbiani et al.<sup>70</sup> |  |
| Goat Anti-Mouse IgG H&L (HRP) | Abcam | Cat #ab6789; RRID: AB_955439 |
| Mouse Anti-Monkey IgG-HRP (SB108a) | SouthernBiotech | Cat #4700-05; RRID: AB_2796069 |
| Mouse anti-IgY-Flag | Schiepers et al.<sup>21</sup> |  |
| Mouse anti-IgY-Strep | Schiepers et al.<sup>21</sup> |  |
| IgY | Gallus Immunotech | Cat #IgY-100 |
| Rabbit Anti-Flag Tag (DYKDDDDK Tag; D6W5B) HRP Conjugate | CellSignaling Technology | Cat #86861S, RRID:AB_2800094 |
| Mouse anti Strep-Tag Classic | BioRad | Cat #MCA2489P, RRID:AB_609796 |
| Alexa Fluor® 647 AffiniPure™ Goat Anti-Mouse IgG, Fcy fragment specific | Jackson ImmunoResearch | Cat #115-605-008, RRID:AB_2338904 |
| Allophycocyanin (APC) AffiniPure™ Goat Anti-Human IgG, Fcy fragment specific | Jackson ImmunoResearch | Cat #109-135-098, RRID:AB_2337690 |
| SureLight® APC Anti-DDDDK tag (FLAG® tag) antibody | Abcam | Cat # ab72569, RRID:AB_1310127 |
| Bacterial and virus strains |  |  |
| SARS-CoV-2 pseudotyped reporter virus | BEI | Cat# NR-53817 |
| SARS-CoV-2 WA1 pseudotyped virus | Cohen et al.<sup>13</sup> |  |
| SARS-CoV-2 Beta pseudotyped virus | Scheid et al.<sup>77</sup> |  |
| SARS-CoV-2 Omicron BA.5 pseudotyped virus | Cohen et al.<sup>14</sup> |  |
| SARS-CoV-2 Omicron XBB.1.5 pseudotyped virus | This paper |  |
| SHC014 pseudotyped virus | Cohen et al.<sup>13</sup> |  |
| WIV1 pseudotyped virus | Cohen et al.<sup>13</sup> |  |
| SARS-CoV-1 (SARS1) pseudotyped virus | Cohen et al.<sup>13</sup> |  |
| Khosta-2 chimera pseudotyped virus | Cohen et al.<sup>14</sup> |  |
| BtKY72 chimera pseudotyped virus | Cohen et al.<sup>14</sup> |  |
| *Bacillus subtilis* strain CVD175 (168 ΔaprE ΔnprE Δvpr Δbpr ΔnprB Δmpr Δepr ΔhtrA ΔwprA spoIIIE::kan hag::ery) | This paper; Ingenza, Ltd |  |
| Biological samples |  |  |
| Immunized non-human primate serum | This paper; University of Washington |  |
| Immunized mouse serum | This paper; Labcorp |  |
| Immunized mouse serum | This paper; The Rockefeller University |  |
| Chemicals, peptides, and recombinant proteins |  |  |
| Heat-Inactivated Fetal Bovine Serum, Optima | Bio-Techne | Cat# S12450H |
| Addavax | InvivoGen | Cat# vac-adx-10 |
| LION | Erasmus et al.<sup>28</sup>; HDT Bio |  |
| Tamoxifen | Sigma | Cat# T5648 |
| 3,3',5,5'-Tetramethylbenzidine substrate | Sigma | Cat# T0565-100ML |
| Britelite Plus reagent | Revvity Health Sciences, Inc (formerly Perkin Elmer) | Cat# 6066769 |
| SuperSignal ELISA Femto Maximum Sensitivity Substrate | ThermoFisher | Cat #37074 |
| Streptavidin APC | eBioscience | Cat #17-4317-82 |
| Mosaic-8b RBD-mi3 nanoparticle vaccine | Cohen et al.<sup>14</sup> |  |
| Admix-8b RBD-mi3 nanoparticle vaccine | This paper |  |
| Homotypic SARS-2 RBD-mi3 nanoparticle vaccine | Cohen et al.<sup>14</sup> |  |
| SARS-CoV-2 WA1 6P Spike protein | Hsieh et al.<sup>84</sup> |  |
| SARS-CoV-2 WA1 RBD-Avi-His | Cohen et al.<sup>13</sup> |  |
| SARS-CoV-2 Beta RBD-Avi-His | Cohen et al.<sup>14</sup> |  |
| SARS-CoV-2 Delta RBD-Avi-His | Cohen et al.<sup>14</sup> |  |
| SARS-CoV-2 Omicron BA.5 RBD-Avi-His | Cohen et al.<sup>14</sup> |  |
| SARS-CoV-2 BA.5-S8 RBD-Avi-His | This paper |  |
| SARS-CoV-2 XBB.1.1 RBD-Avi-His | This paper |  |
| SARS-CoV-2 XBB.1.5 RBD-Avi-His | This paper |  |
| Pang17 RBD-Avi-His | Cohen et al.<sup>13</sup> |  |
| SHC014 RBD-Avi-His | Cohen et al.<sup>13</sup> |  |
| WIV1 RBD-Avi-His | Cohen et al.<sup>13</sup> |  |
| RF1 RBD-Avi-His | Cohen et al.<sup>13</sup> |  |
| SARS-1 RBD-Avi-His | Cohen et al.<sup>13</sup> |  |
| Khosta-2 RBD-Avi-His | Cohen et al.<sup>14</sup> |  |
| BtKY72 RBD-Avi-His | Cohen et al.<sup>13</sup> |  |
| Bm4831 RBD-Avi-His | Cohen et al.<sup>13</sup> |  |
| Yun11 RBD-Avi-His | Cohen et al.<sup>13</sup> |  |
| RsTTS200 RBD-Avi-His | Cohen et al.<sup>14</sup> |  |
| SARS-CoV-2 Beta RBD-His-ST003 | Cohen et al.<sup>14</sup> |  |
| RaTG13 RBD-His-ST003 | Cohen et al.<sup>14</sup> |  |
| Rs4081 RBD-His-ST003 | Cohen et al.<sup>14</sup> |  |
| SHC014 RBD-His-ST003 | Cohen et al.<sup>14</sup> |  |
| Pang17 RBD-His-ST003 | Cohen et al.<sup>14</sup> |  |
| WIV1 RBD-His-ST003 | Cohen et al.<sup>14</sup> |  |
| RF1 RBD-His-ST003 | Cohen et al.<sup>14</sup> |  |
| RmYN02 RBD-His-ST003 | Cohen et al.<sup>14</sup> |  |
| SARS-CoV-2 Beta RBD-ST001-C-tag | This paper, Ingenza Ltd. |  |
| RaTG13 RBD-ST001-C-tag | This paper, Ingenza Ltd. |  |
| Rs4081 RBD-ST001-C-tag | This paper, Ingenza Ltd. |  |
| SHC014 RBD-ST001-C-tag | This paper, Ingenza Ltd. |  |
| Pang17 RBD-ST001-C-tag | This paper, Ingenza Ltd. |  |
| WIV1 RBD-ST001-C-tag | This paper, Ingenza Ltd. |  |
| RF1 RBD-ST001-C-tag | This paper, Ingenza Ltd. |  |
| RmYN02 RBD-ST001-C-tag | This paper, Ingenza Ltd. |  |
| SpyCatcher003-mi3 | Bruun et al.<sup>78</sup>; Ingenza Ltd. |  |
| Critical commercial assays |  |  |
| Expi293 Expression System Kit | ThermoFisher | Cat# A14635 |
| Deposited data |  |  |
| Deep mutational scanning processed data | This paper; GitHub | https://github.com/bjorkmanlab/Mosaic-8b_prevax_fate_mapping |
| Deep mutational scanning sequencing data | This paper; NCBI SRA: Bioproject: PRJNA1067836 and Biosamples: SAMN39552627 SAMN40463706 |  |
| ELISA data | This paper |  |
| Neutralization data | This paper |  |
| Experimental models: Cell lines |  |  |
| Expi293F cells | ThermoFisher | RRID:CVCL_D615 |
| HEK293T cells | Pear et al.<sup>79</sup> | Cat# CCLV-RIE<br>1018 RRID:<br>CVCL_0063 |
| HEK293T-hACE2 | BEI | Cat# NR-52511;<br>RRID:CVCL_A7UK |
| HEK-293T cells expressing high levels of hACE2<br>(consensus Kozak) | Kenneth Matreyek,<br>Case Western<br>Reserve University |  |
| *Pichia pastoris* strain IGZ0038 ($\Delta och1$) | This paper; Ingenza,<br>Ltd | |
| WA1 RBD DMS yeast library | Starr et al.<sup>80</sup> |  |
| XBB.1.5 RBD DMS yeast library | Taylor et al.<sup>81</sup> |  |
| SARS-1 RBD DMS yeast library | Lee et al.<sup>82</sup> |  |
| RmYN02 RBD DMS yeast library | Lee et al.<sup>82</sup> |  |
| PRD-0038 DMS yeast library | Lee et al.<sup>82</sup> |  |
| AWY101 yeast | Wentz and Shusta<sup>83</sup> |  |
| Experimental models: Organisms/strains |  |  |
| cynomolgus macaques | This paper |  |
| BALB/c mice | Charles River<br>Laboratories | RRID:<br>IMSR_JAX:000664 |
| *Igk*<sup>Tag</sup> mice | Schiepers et al.<sup>21</sup>;<br>Jackson Laboratories | strain #038152;<br>RRID:<br>IMSR_JAX:038152 |
| S1pr2-cre<sup>ERT2</sup> mice | Shinnakasu et al.<sup>34</sup> |  |
| Oligonucleotides |  |  |
| primers for DMS Illumina sequencing | Starr et al.<sup>16</sup> | https://github.com/jbloomlab/SARS-CoV-2-RBD_DMS/blob/master/data/primers/primers.csv |
| Recombinant DNA |  |  |
| Expression vectors to produce Spytagged RBDs for<br>nanoparticle vaccine | Cohen et al.<sup>14</sup> | See Recombinant<br>protein section |
| Expression vectors to produce Spytagged-Ctag RBDs<br>for nanoparticle vaccine | This paper, Ingenza<br>Ltd | See Recombinant<br>protein section |
| Expression vectors to produce His-Avi RBDs for ELISAs | This paper, Cohen et<br>al.<sup>13,14</sup> | See Recombinant<br>protein section |
| pPPI4-SARS-CoV-2 S 6P | Hsieh et al.<sup>84</sup> | N/A |
| Expression vectors to produce Spytagged RBDs in<br>*Pichia pastoris* | This paper, Ingenza<br>Ltd | See Recombinant<br>protein section |
| pCVD148 SpyCatcher003-mi3 expression plasmid | This paper; Ingenza<br>Ltd | See Recombinant<br>protein section |
| Expression plasmids of sarbecovirus spikes for making<br>pseudotyped viruses | This paper, Cohen et<br>al.<sup>13,14</sup> | See Bacterial and<br>virus strains section |
| Software and algorithms |  |  |
| Graphpad Prism 10.1.0 | GraphPad | https://www.graphpad.com/ |
| AntibodyDatabase | West et al.<sup>85</sup> |  |
| Sony SH800 software v2.1.6 | Sony | https://www.sonybiotechnology.com/us/instruments/sh800s-cell-sorter/software/ |
| Adobe Illustrator 2024 28.0 | Adobe | https://www.adobe.com/products/illustrator/ |
| Deep mutational scanning processing steps | Greaney et al.<sup>33</sup> | https://github.com/jbloomlab/SARS-CoV-2_Bjorkman_pilot |
| Swift DMS | This paper |  |
| Other |  |  |
| self-amplifying replicon RNA (repRNA) SARS-2 WA1 full length prefusion spike vaccine | University of Washington |  |
| self-amplifying replicon RNA (repRNA) bivalent Beta/Delta SHARP vaccine | University of Washington |  |
| self-amplifying replicon RNA (repRNA) trivalent Beta/Delta/WA1 SHARP vaccine | University of Washington |  |
| self-amplifying replicon RNA (repRNA) bivalent WA1/BA.1 full length prefusion spike vaccine | Erasmus et al.<sup>28</sup>, HDT Bio |  |
| DNA SARS-2 WA1 full length prefusion spike vaccine | University of Washington |  |
| DNA bivalent Beta/Delta SHARP vaccine | University of Washington |  |
| DNA trivalent Beta/Delta/WA1 SHARP vaccine | University of Washington |  |
| Pfizer-like mRNA-LNP SARS-2 WA1 Spike vaccine | Helix Biotech |  |
| Pfizer-like mRNA-LNP SARS-2 BA.5 Spike vaccine | Helix Biotech |  |
| Moderna COVID-19 vaccine | Moderna |  |
| ChAdOx1 nCoV-19 vaccine | Viral Vector Core Facility, Oxford University |  |
| Nucleoside-modified mRNA-LNP WA1 Spike vaccine | Laczko et al.<sup>86</sup> | GenBank MN908947.3 |
| Nunc MaxiSorp 384-well plates | Sigma | Cat #P6491 |
| 96-Well Half Area Microplates | Greiner Bio-One | Cat #675061 |
| Capture Select™ C-tagXL Affinity Matrix | Thermo Fisher | Cat # 2943072050 |

### EXPERIMENTAL MODEL AND SUBJECT DETAILS

#### Mammalian cell lines

HEK293T cells for pseudovirus production (RRID:CVCL_0063) were cultured in Dulbecco’s modified Eagle’s medium (DMEM, Gibco) supplemented with 10% heat-inactivated fetal bovine serum (FBS, Bio-Techne) and 1% Penicillin/Streptomycin (Gibco) and 1% L-Glutamine (Gibco) at 37 °C and 5% CO_2_.

HEK293T-hACE2 (RRID:CVCL_A7UK) cells^87^ for pseudovirus neutralization assays were cultured in DMEM (Gibco) supplemented with 10% heat-inactivated FBS (Bio-Techne), 5 mg/ml gentamicin (Sigma-Aldrich), and 5 mg/mL blasticidin (Gibco) at 37 °C and 5% CO_2_.

Expi293 cells (RRID:CVCL_D615, ThermoFisher) for protein expression were maintained at 37 °C and 8% CO_2_ in ThermoFisher’s Expi293 expression medium. Transfections were done with ThermoFisher’s Expi293 Expression System Kit and maintained under shaking at 130 rpm.

293T-based cell lines were derived from female donors and not specially authenticated.

#### Yeast

*Pichia pastoris* strain IGZ0038 (Δ*och1;* Ingenza Ltd) was used as the recipient genetic background for expression of the 8 RBD-SpyTag001-Ctag sequences. *Pichia* strains were maintained on vegetarian YPD (1% [w/v] yeast extract [Oxoid], 2% (w/v) peptone from soybean [Millipore] and 2% (w/v) D-glucose [Alfa Aesar]) at 30°C.

#### Bacteria

Asporogenic and protease-deficient *Bacillus subtilis* strain CVD175 (168 ΔaprE ΔnprE Δvpr Δbpr ΔnprB Δmpr Δepr ΔhtrA ΔwprA spoIIE::kan hag::ery; Ingenza Ltd.) was used as genetic background for plasmid-borne expression of SpyCatcher003-mi3. The expression strain was maintained on Terrific Broth medium (Merck) supplemented with 10 μg/mL chloramphenicol, 10 μg/mL kanamycin, and 1% (v/v) glycerol at 37 °C.

#### Viruses

Pseudoviruses were generated by transfecting HEK293T cells (RRID:CVCL_0063) using Fugene HD (Promega) with a luciferase reporter construct and coronavirus pseudovirus constructs as described.^88^ Cells were co-transfected with HIV-1-based lentiviral plasmids, a luciferase reporter construct, and a coronavirus spike construct, leading to the production of lentivirus-based pseudovirions expressing the coronavirus spike protein at the pseudovirion surface. Supernatants were harvested 48-72 hours post-transfection, filtered, and stored at -80 °C until use. Pseudovirus infectivity was determined by titration on HEK293T_ACE2_ cells.

#### Non-human primates

Fourteen 5-year-old male and female cynomolgus macaques of Mauritian origin were utilized for this study. Animals were pair housed with full run through contact at the Washington National Primate Research Center (WaNPRC) accredited by AAALAC (American Association for the Accreditation of Laboratory Animal Care). Animal housing areas were cleaned daily and kept at 30-70% humidity and 72-82^◦^F. Animal diet consisted of fresh water on demand and commercial monkey chow supplemented with fruits and vegetables daily. Environmental enrichment included paired housing, food enrichment, rotating toys, music, movies, foraging opportunities, and all animals’ health and well-being were monitored daily by animal care staff. The University of Washington Institutional Animal Care and Use Committee (IACUC # 4266-02) reviewed and approved all animal protocols, which were also compliant with the U.S. Department of Health and Human Services *Guide for the Care and Use of Laboratory Animals*. The 14 pre-vaccinated NHPs were stratified into three groups 24 weeks after their last vaccination based on weight, age, sex, and Ab neutralization titers against SARS-2 WA1 D614G (Figure S2A).

#### Mice

Mouse procedures were approved by the Caltech, Labcorp, or Rockefeller Institutional Animal Care and Use Committees. Six- to 7-week-old female BALB/c mice were purchased from Charles River Laboratories (RRID: IMSR_JAX:000664) and housed at Labcorp Drug Development, Denver, PA for immunization studies. All animals were healthy upon receipt and were weighed and monitored during a 7 day acclimation period before the start of the study. Mice were randomly assigned to experimental groups of 10-14 animals. Up to 10 mice were cohoused together and cages were kept in a climate controlled room at 68 to 79 °C at 50±20% relative humidity. Mice were provided Rodent Diet #5001 (Purina Lab Diet) ad libitum.

*Igk*^Tag^ mice (Jackson Laboratories strain #038152, C57BL/6-*Igk*^em1Vic^/J, RRID: IMSR_JAX:038152), were generated on the C57BL/6 background at the Rockefeller University as described.^21^ *Igk*^Tag^ mice were bred with S1pr2-Cre^ERT2^ BAC-transgenic mice^34^ kindly provided by T. Kurosaki and T. Okada (U. Osaka, RIKEN-Yokohama), to obtain S1pr2*-*creERT2*.Igk*^Tag/Tag^ (S1pr2*-Igk*^Tag^) mice. S1pr2*-Igk*^Tag^ mice were held at the immunocore clean facility at Rockefeller under specific pathogen-free conditions.

### METHOD DETAILS

#### Reagents used for pre-vaccinations

NHPs were vaccinated over the course of 34 weeks with various nucleic acid vaccines prior to RBD-nanoparticle vaccination (Figure 2, Figure S2, Table S1). At weeks -64 and -58, immunizations comprised 20 ng of either self-amplifying replicon RNA (repRNA) or DNA encoding SARS-2 WA1 full length prefusion spike. At week -47, bivalent immunogens comprised 40 ng repRNA or DNA encoding SARS-2 B.1.351 (Beta) and B.1.617 (Delta) RBDs fused to the influenza H3N2 hemagglutinin HA2 stem domain (SHARP^27^; designed by A.I.R. and J.D.P.). At week -30, trivalent SHARP immunogens comprised 60 ng repRNA or DNA encoding the Beta, Delta, and WA1 RBDs fused to the influenza H3N2 hemagglutinin HA2 stem domain. Vaccines were formulated and administered into different animals through different modalities: n=4 macaques received a self-amplifying replicon RNA vaccine formulated with a cationic nanocarrier (repRNA-LION) and delivered IM as described;^28^ n=5 macaques received the same repRNA vaccine formulated on 1 micron gold particles and delivered by gene gun into the skin; and n = 5 macaques received DNA vaccine adjuvanted with a plasmid expressing IL-12 formulated at a 10:1 vaccine plasmid to adjuvant ratio on 1 micron gold particles and delivered by gene gun into the skin as described.^89^

Because we were unable to obtain licensed Pfizer-BioNTech or Moderna mRNA-LNP vaccines for research purposes from the respective companies, Pfizer-like mRNA-LNP formulations for WA1 and BA.5 were purchased from Helix Biotech and used for mRNA-LNP vaccinations (Figure 3; Figure S3; Table S1). Bivalent mRNA-LNP (Figure 3F-J; Figure S3F-N; Table S1) was prepared by mixing WA1 mRNA-LNP and BA.5 mRNA-LNP in a 1:1 ratio by mRNA mass. Moderna COVID-19 vaccine used as a boost in the ChAdOx1 study (Figure 4, 5; Figure S4, S5; Table S1) was the gift of Jeffrey Ravetch, Rockefeller University. Research grade ChAdOx1 nCoV-19 was kindly provided by the Viral Vector Core Facility, Oxford University (Figure 4, 5; Figure S4, S5; Table S1). Nucleoside-modified mRNA-LNP encoding SARS-2 spike protein WA1 strain (GenBank MN908947.3) used for S1pr2-*Igk*^Tag^ experiments (Figure 6; Table S1) was generated at the University of Pennsylvania and Acuitas as described.^86^

#### Immunization of NHPs

Mosaic-8b or homotypic SARS-2 Beta produced at Caltech as described^14^ were diluted with Dulbecco’s PBS and mixed 1:1 (v/v) with AddaVax for a final vaccine dose of 25 µg of RBD equivalents in 0.5 mL delivered IM. WA1/BA.1 spike repRNA was produced by HDT Bio as described.^28^ 25 µg WA1 and 25µg BA.1 repRNA were individually co-formulated with citrate buffer, water, sucrose, and LION^28^ and then mixed together for a final vaccine dose of 50 µg in 0.5 mL delivered IM. Prior to vaccine delivery, animals were sedated and hair over the injection site was clipped and wiped with ethanol.

#### Immunization of mice

mRNA-LNP and ChAdOx-1 pre-vaccinated mouse experiments were conducted using 6-7 week old female BALB/c mice (Charles River Laboratories) with 10-14 animals in each cohort.

Mice in the mRNA-LNP pre-vaccination study were vaccinated IM with 20 µL of WA1 mRNA-LNP at day -192 and day -171 containing 1 µg mRNA diluted in PBS (Figure 3, Figure S3, Table S1). Mice in groups that received a bivalent mRNA-LNP boost were additionally vaccinated IM with 20 µL of WA1/BA.5 mRNA-LNP (0.5 µg WA1 and 0.5 µg BA.5 mRNA) at day -73 (Figure 3F, Figure S3F). Mice were immunized IM with 5 µg of protein nanoparticle (RBD equivalents) in 100 µL containing 50% v/v AddaVax adjuvant on days 0 and 28 or received an additional dose of 1 µg WA1 mRNA-LNP at day 0.

Mice in the ChAdOx-1 study (Figure 4, Figure S4, Table S1) were vaccinated IM with 1×10^8^ IU of ChAdOx-1 on days -163 and -135. Mice were immunized IM with 5 µg of protein nanoparticle (RBD equivalents) in 100 µL containing 50% v/v AddaVax adjuvant on days 0 and 28 or received WA1 mRNA-LNP (1 µg mRNA in PBS) or 1×10^8^ IU of ChAdOx-1 at day 0. Mice in both studies were bled as indicated in Figure 3A and Figure 4A, including day 0 and day 28 via tail veins, and then euthanized at day 56 and bled through cardiac puncture. Blood samples were allowed to clot and serum was harvested and stored at -80 °C until use.

For Ab fate-mapping experiments, immune responses were induced in 7- to 12-week-old S1pr2-*Igk*^Tag^ mice (males and females) by IM immunization of right quadricep muscles with 3 µg WA1 mRNA-LNP (Figure 6, Figure S6, S7, Table S1). Mice were re-immunized using the same procedure 28 days later. These two primary immune responses were fate-mapped by oral gavage of 200 µl tamoxifen (Sigma) dissolved in corn oil at 50 mg/ml, on days 4, 8, 12 after the first and on days 6 and 12 after the second immunization. The third immunization was performed 28 days later with either mosaic-8b or homotypic nanoparticles (5 µg, adjuvanted 1:1 (v/v) with AddaVax) in the left quadricep muscles. The fourth immunization occurred 28 days after the third (month 3 of the experiment). Blood samples were collected at key time points throughout the experiment via cheek puncture into microtubes prepared with clotting activator serum gel (Sarstedt; catalog #41.1378.005).

#### Expression and purification of RBDs

Expression vectors encoding RBDs from spike proteins in SARS-2 Beta (GenBank QUT64557.1), SARS-2 Wuhan-Hu-1 (GenBank MN985325.1), RaTG13-CoV (GenBank QHR63300), SHC014-CoV (GenBank KC881005), Rs4081-CoV (GenBank KY417143), pangolin17-CoV (GenBank QIA48632), RmYN02-CoV (GSAID EPI_ISL_412977), Rf1-CoV (GenBank DQ412042), WIV1-CoV (GenBank KF367457), Yun11-CoV (GenBank JX993988), BM-4831-CoV (GenBank NC014470), BtKY72-CoV (GenBank KY352407), Khosta-2 CoV (QVN46569.1), RsSTT200-CoV (EPI_ISL_852605), LYRa3 (AHX37569.1) and SARS-CoV S (GenBank AAP13441.1) were constructed as described^13,14,90^ to include a C-terminal hexahistidine tag (6xHis; G-HHHHHH) and SpyTag003 (RGVPHIVMVDAYKRYK)^24^ (for coupling to SpyCatcher003-mi3 for SARS-2 Beta, RaTG13, SHC014, Rs4081, pangolin17-CoV, RmYN02, Rf1, and WIV1 RBDs) or a 15-residue Avi-tag (GLNDIFEAQKIEWHE) followed by a 6xHis tag (for ELISAs). RBDs were purified from transiently-transfected Expi293F cell (ThermoScientific) supernatants as described.^14^ A soluble SARS-2 trimer with 6P stabilizing mutations^84^ and monoclonal IgGs were produced as described.^17,18,58^

SpyTagged RBDs for RBD-nanoparticles used in mRNA-LNP pre-vaccinated mice were produced in *Pichia pastoris* strain IGZ0038 (Δ*och1;* Ingenza Ltd). Expression cassettes were constructed to encode RBD residues for SARS-2 Beta, RaTG13, SHC014, Rs4081, pangolin17-CoV, RmYN02, Rf1, and WIV1 with a C-terminal Gly-Ser linker, SpyTag, and C-Tag for affinity purification (GGGSGGGSGGGSGSGAHIVMVDAYKPTKGSGEPEA).^91^ Vectors harboring expression cassettes were linearized with SwaI and then transformed into *P. pastoris* by electroporation for targeted integration at the *AOX1* locus. Transformants were selected on vegetarian Yeast Extract-Peptone-Dextrose (YPD) medium supplemented with 1 M D-sorbitol (Merck) and 250 *µ*g mL^-1^ Zeocin (InvivoGen). For RBD-SpyTag001-Ctag protein expression, transformed *Pichia pastoris* strains were cultured in a 3 L Fermac 320 bioreactor (Electrolab Biotech) using a defined medium fed-batch fermentation process supplemented with glucose. The inoculum was grown in a 500 mL Erlenmeyer flask at 30 °C and 200 rpm. Bioreactors were controlled at 30 °C, pH 5.5 and 30% dissolved oxygen prior to induction and 25 °C, pH 6 and 30% dissolved oxygen post-induction. Induction was performed using a methanol feed supplemented with glucose. RBDs were purified via CaptureSelect^TM^ C-tagXL affinity columns (ThermoFisher) according to the manufacturer’s instructions (2 M MgCl_2_ elution buffer), followed by dialysis against TBS, pH 8.0 (20 mM Tris-HCl, 150 mM NaCl, pH 8.0). RBDs were then concentrated and purified by size-exclusion chromatography on a Superdex 200 Increase column (Cytiva) in TBS, pH 8.0.

#### Expression and purification of SpyCatcher003-mi3 nanoparticles

SpyCatcher003-mi3 nanoparticles^78^ were expressed in *B. subtilis* to avoid endotoxins associated with production in *E. coli*. Strain CVD175 (*B. subtilis* 168 ΔaprE ΔnprE Δvpr Δbpr ΔnprB Δmpr Δepr ΔhtrA ΔwprA spoIIE::kan hag::ery; Ingenza Ltd) was transformed with high-copy plasmid pCVD148 encoding for IPTG-inducible expression and secretion of SpyCatcher003-mi3. The resulting strain CVD326 was cultured in Terrific Broth medium (Merck) supplemented with 10 μg/mL chloramphenicol, 10 μg/mL kanamycin and 1% (v/v) glycerol at 37 °C with 250 rpm agitation. When OD_600_ reached a value of 1.0, SpyCatcher003-mi3 expression was induced for 20 hours by addition of IPTG to a final concentration of 1 mM. The culture supernatant was harvested by centrifugation at 4424 x g for 45 minutes at 4°C. SpyCatcher003-mi3 particles were then precipitated by addition of ammonium sulfate at a final concentration of 20% (w/v) with incubation for 45 min at room temperature on a stirrer plate. Precipitated particles were pelleted by centrifugation at 4424 × g for 45 min at 4 °C. The obtained pellet was resuspended in TBS (20 mM Tris, 150 mM NaCl, pH 8.0) and filtered through a 0.22 µm syringe filter. Glycerol was added to a final concentration of 10% (v/v), and purified nanoparticles were stored at -80°C until use.

#### Preparation of RBD SpyCatcher-mi3 nanoparticles

Purified SpyCatcher003-mi3 was incubated with a 2-fold molar excess (total RBD to mi3 subunit) of SpyTagged RBD (an equimolar mixture of eight RBDs for mosaic-8b, seven RBDs (all RBDs in mosaic-8b except for the SARS-2 RBD), or only one RBD for homotypic nanoparticles) overnight at room temperature in TBS. Conjugated RBD-mi3 particles were separated from free RBDs by SEC on a Superose 6 10/300 column (GE Healthcare) equilibrated with PBS (20 mM sodium phosphate pH 7.5, 150 mM NaCl). RBD-mi3 nanoparticle conjugations were assessed via SDS-PAGE and western blot. Concentrations of conjugated mi3 particles are reported based on RBD content determined using a Bio-Rad Protein Assay. RBD-mi3 nanoparticles used for immunization of NHPs, ChAdOx nCoV-19-vaccinated mice, and fate mapping studies in mice were made from Expi293-expressed RBDs as previously described.^13,14^ RBD-mi3 nanoparticles used to immunize mice previously vaccinated with mRNA-LNP were produced from *Pichia*-expressed RBDs. *Pichia*-produced and Expi293-produced RBD-mi3 nanoparticles elicit comparable immune responses in immunized mice as do RBD-mi3 nanoparticles constructed using *Bacillus* (this study) or *E. coli* mi3 (previous studies^13,14^) (A.A.C., J.R.K., H.G., and P.J.B., unpublished results). RBD-mi3 nanoparticles were aliquoted and flash frozen in liquid nitrogen before being stored at -80 °C until use. RBD-nanoparticles were characterized by SEC and SDS-PAGE.

#### Serum ELISAs

Affinity purified His-tagged RBD produced by Expi293 transient transfection was diluted in 0.1 M NaHCO_3_ pH 9.8 (20 µL of a 2.5 µg/mL solution) and coated onto Nunc® MaxiSorp™ 384-well plates (Sigma) and incubated overnight at 4°C. Plates were blocked with 3% bovine serum albumin (BSA), 0.1% Tween 20 in TBS (TBS-T) for 1 hr at room temperature and then the blocking was removed by aspiration. 1:100 dilutions of NHP or mouse serum were serially diluted by 3.1-fold in TBS-T/3% BSA and then added to the plates for 3 hr at room temperature followed by washing with TBS-T. After incubating for one hour in a 1:100,000 dilution of HRP-conjugated anti-IgG secondary (goat anti-mouse IgG (Abcam; RRID: AB_955439) for mouse serum or mouse-anti-monkey IgG (SouthernBiotech; RRID: AB_2796069) for NHP serum), the plates were washed with TBS-T and SuperSignal™ ELISA Femto Maximum Sensitivity Substrate (ThermoFisher) was added following manufacturer’s instructions. Luminescence was read at 425 nm. We used Graphpad Prism 10.1.0 to plot and analyze curves assuming a one-site binding model with a Hill coefficient to obtain midpoint titers (ED_50_ values). For the WA1 mRNA-LNP pre-vaccinated mouse experiments (Figure 3; Figure S3), ED_50_s were normalized by dividing the ED_50_ response at day 28 or day 56 by the ED_50_ response at day 0 to control for differences in Ab binding response between groups from the pre-vaccination (Figure S3). Mean ED_50_s are reported for ELISA data in Figure 2,3, and 4 and associated supplementary figures, as previously described.^13,14^

FLAG/Strep ELISAs for molecular fate-mapping experiments were performed side-by-side and with internal standards on each 96-well plate as described^21^. Plates were incubated overnight at 4 °C with 2 µg of RBD in PBS, wells for standard curves were coated with 10 µg/ml purified IgY (Gallus Immunotech). Plates were washed with PBS-Tween (PBS + 0.05% Tween20; Sigma) and blocked with 2.5% BSA in PBS for 2 hours at room temperature. Serum samples were diluted 1:100 in PBS and serially diluted in 3-fold steps to determine endpoint titers. Mouse anti-IgY mAb-FLAG or mAb-Strep were also serially titrated in 3-fold dilutions. After 2 hours of incubation, followed by washing, rabbit anti-FLAG-HRP (clone D6W5B, CellSignalingTechnology #86861S, RRID:AB_2800094) or anti-Strep (clone Strep-tag II StrepMAB-Classic, Biorad #MCA2489P, RRID:AB_609796) detection Abs were added for 30 minutes. Dilutions of detection Abs were defined such that FLAG- and Strep-tagged mAb dilution curves overlapped. After washing with PBS-Tween, samples were incubated with 3,3′,5,5′-Tetramethylbenzidine substrate (slow kinetic form, Sigma) and the reaction was stopped with 1N HCl. Optical Density (OD) absorbance was measured at 450 nm on a Fisher Scientific accuSkan FC plate reader. To normalize FLAG and Strep endpoint titers, the serum titer dilution was calculated at which each sample passed the threshold OD value of its respective mAb at 20 ng/µl. Titers were calculated by logarithmic interpolation of the dilutions with readings immediately above and immediately below the mAb OD used. Antibody OD and endpoint titer levels were plotted in GraphPad Prism 10.1.0, titer values below 100 were set to 100, the top dilution and therefore the limit of detection (LOD). Median anti-RBD titers are reported in Figure 6 and associated supplementary figures, consistent with analyses of previous molecular fate-mapping experiments.^21^

#### Serum pseudovirus neutralization assays

Lentiviral-based pseudoviruses were prepared as described^70,88^ using genes encoding S protein sequences lacking C-terminal residues in the cytoplasmic tail: 21 residue (SARS-2 variants, WIV1, and SHC014) or 19 residue cytoplasmic tail deletions (SARS-CoV, Khosta-2-SARS-1 chimera, BtKY72-SARS-1 chimera). BtKY72 (containing K493Y/T498W substitutions) and Khosta-2 pseudoviruses were made with chimeric spikes in which the RBD from SARS-1 (residues 323-501) was substituted with the RBD from BtKY72 K493Y/T498W (residues 327-503) or Khosta-2 (residues 324-500) as described^92^. For neutralization assays, pseudovirus was incubated with three-fold serially diluted sera from immunized NHPs or mice for 1 hour at 37°C, then the serum/virus mixture was added to 293T_ACE2_ target cells or high-hACE2 HEK-293T cell line expressing hACE2 encoded with a consensus Kozak sequence (kindly provided by Kenneth Matreyek, Case Western Reserve University) for SHC014 assays and incubated for 48 hours at 37°C. Media was removed, cells were lysed with Britelite Plus reagent (Revvity Health Sciences), and luciferase activity was measured as relative luminesce units (RLUs). Relative RLUs were normalized to RLUs from cells infected with pseudotyped virus in the absence of antiserum. Half-maximal inhibitory dilutions (ID_50_ values) were derived using 4-parameter nonlinear regression in AntibodyDatabase.^85^

### DMS

#### Yeast library sorting to identify mutations that reduce binding to polyclonal serum and monoclonal Abs for epitope mapping

DMS studies used to map epitopes recognized by serum Abs were performed in biological duplicates using independent mutant libraries (WA1,^80^ XBB.1.5,^81^ SARS-1,^82^ RmYN02,^82^ and PRD-0038,^82^ generously provided by Tyler Starr, University of Utah) as described previously.^14,33^ Serum samples were heat inactivated for 30 min at 56 °C before DMS analysis. In order to remove non-RBD specific yeast-binding Abs, heat-inactivated sera were then incubated twice with 50 OD units of AWY101 yeast transformed with an empty vector to remove yeast-binding Abs. Monoclonal Abs for DMS analyses were not heat inactivated or yeast depleted. RBD libraries were induced for RBD expression in galactose-containing synthetic defined medium with casamino acids (6.7g/L Yeast Nitrogen Base, 5.0 g/L Casamino acids, 1.065 g/L MES acid, and 2% w/v galactose + 0.1% w/v dextrose). After inducing for 18 hours, cells were washed 2x and then incubated with serum or monoclonal Abs (dilutions chosen to give sub-saturating binding to RBDs; data posted in https://github.com/bjorkmanlab/Mosaic-8b_prevax_fate_mapping) for 1 hour at RT with gentle agitation after which cells were washed 2x and labeled for 1 hour with secondary Ab: either 1:200 Alexa Fluor-647-conjugated goat anti-mouse-IgG Fc-gamma, Jackson ImmunoResearch 115-605-008, RRID:AB_2338904) for serum mouse Abs, 1:200 Allophycocyanin-AffiniPure Goat Anti-Human IgG/Fcγ Fragment Specific (Jackson ImmunoResearch 109-135-098, RRID:AB_2337690) for human monoclonal Abs, Streptavidin-APC (eBioscience 17-4317-82) for Strep tag Abs, and SureLight® APC Anti-DDDDK tag (Abcam ab72569, RRID:AB_1310127) for Flag tag Abs.

Stained yeast cells were processed on a Sony SH800 cell sorter. Cells were gated to capture RBD mutants that had reduced antibody binding for a relatively high degree of RBD expression (data posted in https://github.com/bjorkmanlab/Mosaic-8b_prevax_fate_mapping). For each sample, cells were collected up until around 5 × 10^6^ RBD^+^ cells were processed (which corresponded to around 5 × 10^5^-1 × 10^6^ RBD^+^ Ab escaped cells; data posted in https://github.com/bjorkmanlab/Mosaic-8b_prevax_fate_mapping). Antibody-escaped cells were grown overnight in a synthetic defined medium with casamino acids (6.7 g/L Yeast Nitrogen Base, 5.0 g/L Casamino acids, 1.065 g/L MES acid, and 2% w/v dextrose + 100 U/mL penicillin + 100 µg/mL streptomycin) to expand cells prior to plasmid extraction. DNA extraction and Illumina sequencing were carried out as previously described.^93^ Raw sequencing data are available on the NCBI SRA under BioProject PRJNA1067836, BioSample SAMN40463706 (for serum and mAb DMS in Figure 5 and Figure S5) and SAMN39552627 (for molecular fate mapping in Figure 6 and Figure S6). Escape fractions were computed using processing steps described previously^33,93^ (https://github.com/jbloomlab/SARS-CoV-2_Bjorkman_pilot) and implemented using a new Swift DMS program (available from authors upon request). Escape scores were calculated with a filter to remove variants with deleterious mutations that escape binding due to poor expression, >1 amino acid mutation, or low sequencing counts as described.^93,94^

Static line plot visualizations of escape maps were created using Swift DMS.^93^ In all cases, the line height indicates the escape score for that amino acid mutation, calculated as previously described.^93^ In some visualizations, RBD sites are categorized based on epitope region,^17^ which we defined as: class 1 (pink) (residues 403, 405, 406, 417, 420, 421, 453, 455-460, 473-478, 486, 487, 489, 503, 504); class 2 (purple) (residues 472, 479, 483-485, 490-495), class 3 (blue) (residues 341, 345, 346, 354-357, 396, 437-452, 466-468, 496, 498-501,462), class 4 (orange) (residues 365-390, 408). For structural visualizations, an RBD surface (PDB 6M0J) was colored by the site-wise escape metric at each site, with magenta scaled to be the maximum used to scale the y-axis for serum and monoclonal Abs, red scaled to be the maximum for Strep tag Abs, and blue scaled to be the maximum for Flag tag responses, and gray indicating no escape. Residues that exhibited the greatest escape fractions were highlighted with their residue number and colored according to epitope class.

#### Quantification and statistical analyses

We used pairwise comparisons, a method to evaluate sets of mean binding or neutralization titers against individual viral strains for different immunization cohorts, to determine whether results from different cohorts were significantly different from each other. Statistically significant titer differences between immunized groups of NHPs and mice for ELISAs and neutralization assays were determined using analysis of variance (ANOVA) followed by Tukey’s multiple comparison post hoc tests with the Geisser-Greenhouse correction, with pairing by viral strain, of ED_50_s/ID_50_s (converted to log_10_ scale) calculated using GraphPad Prism 10.1.0. To address whether our data were normally distributed (a requirement for Tukey’s multiple comparison tests), we used quantile-quantile (Q-Q) probability plots to compare actual and predicted binding and neutralization titers. Most of the data were normally distributed. However, two circumstances resulted in deviations from normal distributions: (1) Setting of background binding or neutralization data points to limits of detection (titers of 1:50 or 1:100); and (2) Neutralization or binding titers against the WA1 spike antigen (D614G pseudovirus or WA1 spike) that was matched to the strain in the COVID vaccines used for pre-vaccination, which resulted in skewing the distributions because of high titers and inclusion of non-RBD Ab epitopes.

In Figure 6D, the linear regression of primary addiction index against antigenic distance score was calculated using Prism.

---

**Figure S1.**
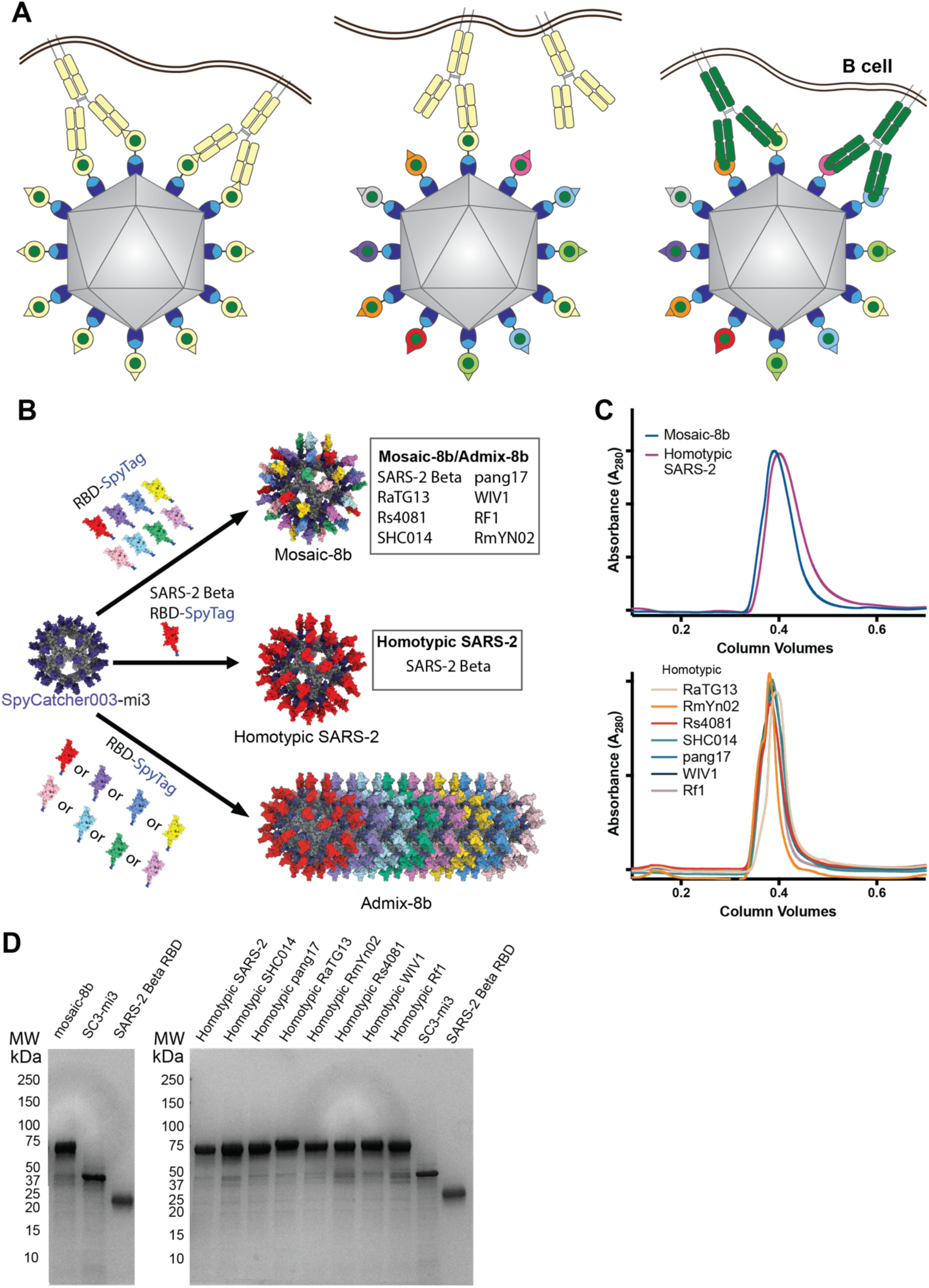
Preparation of RBD-mi3 nanoparticles, related to Figure 1. (A) Hypothesis illustrating potential mechanism for mosaic RBD-nanoparticle induction of cross-reactive Abs. Left: Both Fabs of a strain-specific membrane-bound BCR can bind to a strain-specific epitope (pale yellow triangle) on yellow antigens attached to a homotypic nanoparticle. Middle: Strain-specific BCRs can only bind with one Fab to a strain-specific epitope (triangle) on yellow antigen attached to a mosaic nanoparticle. Right: Cross-reactive BCRs can bind with both Fabs to a common epitope present on adjacent antigens (green circles) attached to a mosaic particle, but not to strain-specific epitopes (triangles). (B) Schematic of construction of mosaic-8b, homotypic SARS-2, and admix-8b RBD-mi3 nanoparticles made using models constructed with coordinates of an RBD (PDB 7BZ5), SpyCatcher (PDB 4MLI), and an i3-01 nanoparticle (PDB 7B3Y). (C) Superose 6 10/300 size exclusion chromatography profile after RBD conjugations to mi3 showing peaks for RBD-mi3 nanoparticles. (D) Coomassie-stained SDS-PAGE gel of RBD-coupled nanoparticles, unconjugated RBDs, and free SpyCatcher003-mi3 particles (SC3-mi3).

**Table S1.**
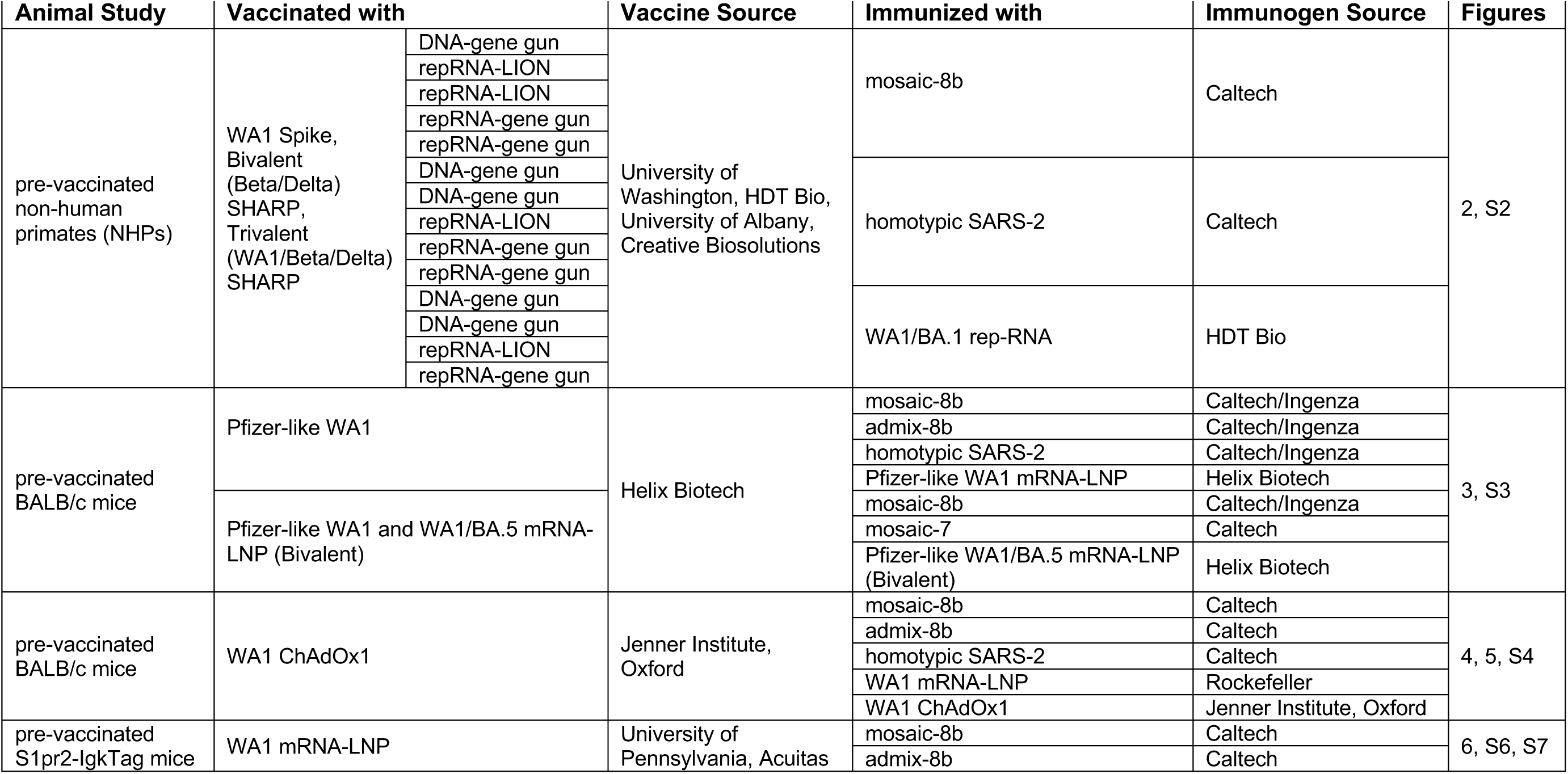
Summary of vaccines and immunogens, related to Figure 1.

**Figure S2.**
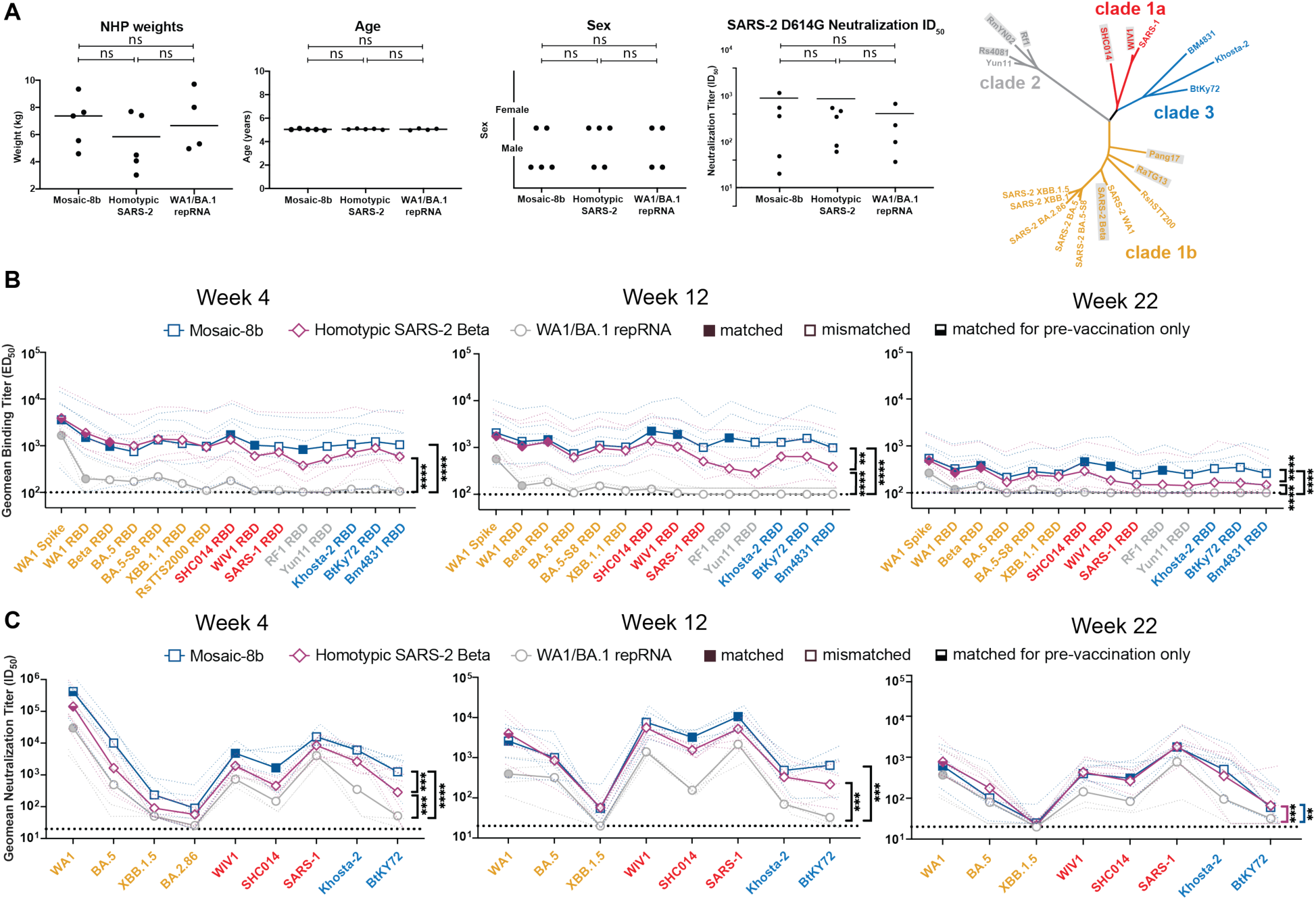
RBD-nanoparticle immunizations in previously-vaccinated NHPs elicit cross-reactive Ab responses, related to Figure 2. Data are shown for ELISA and neutralization analyses for serum samples from weeks 4, 12, and 22. (Data for samples from weeks 0, 2, 8, and 10 are in Figure 2). Geometric means of ED_50_ or ID_50_ values for all animals in each cohort are indicated by symbols connected by thick colored lines; results for individual animals are indicated by dotted lines. Mean titers against indicated viral antigens or pseudoviruses were compared pairwise across immunization cohorts by Tukey’s multiple comparison test with the Geisser-Greenhouse correction (as calculated by GraphPad Prism). Significant differences between cohorts linked by vertical lines are indicated by asterisks: p<0.05 = *, p<0.01 = **, p<0.001 = ***, p<0.0001 = ****. (A) Left: Stratification of pre-vaccinated NHPs into groups used for immunizations with mosaic-8b, homotypic SARS-2, and WA1/BA.1 repRNA based on weight, age, sex, and neutralization ID_50_ values (ns = no significant difference; Mann-Whitney test). Neutralization ID_50_s were derived from samples taken at week -8 in the vaccination/immunization regimen in Figure 2A. Right: Phylogenetic tree of selected sarbecoviruses calculated using a Jukes-Cantor generic distance model using Geneious Prime® 2023.1.2 based on amino acid sequences of RBDs aligned using Clustal Omega.^97^ RBDs included in mosaic-8b are highlighted in gray rectangles. (B) Geometric mean ELISA binding titers at the indicated weeks after immunization with mosaic-8b, homotypic SARS-2, or bivalent WA1/BA.1 repRNA against indicated viral antigens. (C) Geometric mean neutralization titers at the indicated weeks after immunization with mosaic-8b, homotypic SARS-2, or bivalent WA1/BA.1 repRNA against indicated sarbecovirus pseudoviruses.

**Figure S3.**
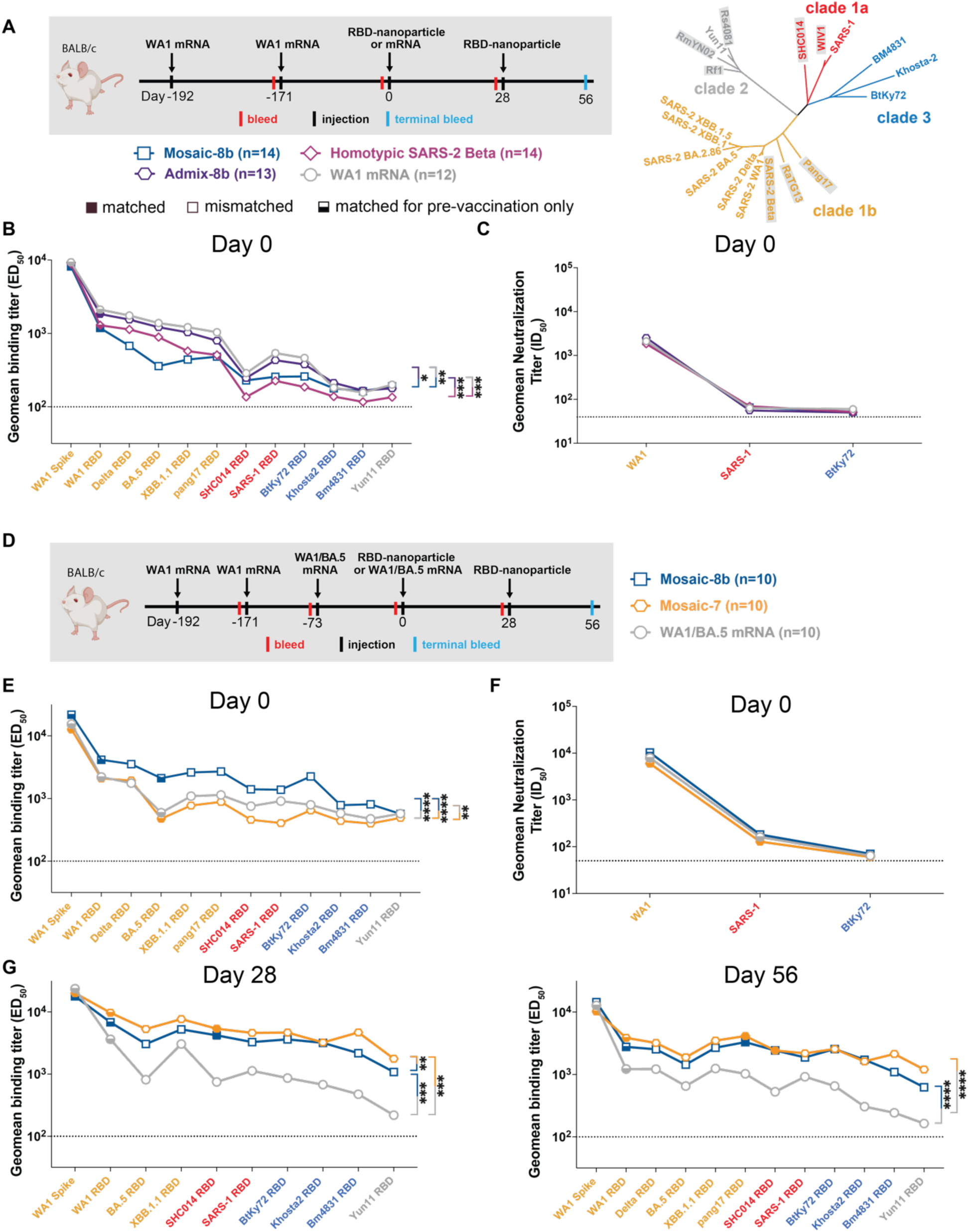
Cohorts of mRNA-LNP vaccinated mice showed significant differences in midpoint ED_50_ titers at day 0 (prior to RBD-nanoparticle immunizations), related to Figure 3. Geometric means of ED_50_ or ID_50_ values for all animals in each cohort are indicated by symbols connected by thick colored lines. Mean titers against indicated viral antigens or pseudoviruses were compared pairwise across immunization cohorts by Tukey’s multiple comparison test with the Geisser-Greenhouse correction (as calculated by GraphPad Prism). Significant differences between cohorts linked by vertical lines are indicated by asterisks: p<0.05 = *, p<0.01 = **, p<0.001 = ***, p<0.0001 = ****. (A) Left: Schematic of vaccination/immunization regimen for panels B and C. Mice were vaccinated at the indicated days prior to RBD-nanoparticle prime and boost immunizations at days 0 and 28 or mRNA-LNP prime immunization at day 0. Right: Phylogenetic tree of selected sarbecoviruses calculated using a Jukes-Cantor generic distance model using Geneious Prime® 2023.1.2 based on amino acid sequences of RBDs aligned using Clustal Omega.^97^ RBDs included in mosaic-8b are highlighted in gray rectangles. (B) Geometric mean ELISA binding titers of serum from mice assigned to each cohort at day 0 (192 and 171 days after the first and second mRNA-LNP vaccinations but prior to nanoparticle or additional mRNA-LNP immunizations) showing significant differences between cohorts prior to immunizations. (C) Geometric mean neutralization binding titers of serum from mice assigned to each cohort at day 0 (192 and 171 days after the first and second mRNA-LNP vaccinations but prior to nanoparticle or additional mRNA-LNP immunizations). (D) Schematic of vaccination/immunization regimen for panels E-G. Mice were vaccinated at the indicated days prior to RBD-nanoparticle prime and boost immunizations at days 0 and 28 or mRNA-LNP prime immunization at day 0. (E) Geometric mean ELISA binding titers of serum from mice assigned to each cohort at day 0 (192, 171, and 73 days after the first, second, and third mRNA-LNP vaccinations but prior to nanoparticle or additional mRNA-LNP immunizations) showing significant differences between cohorts prior to immunizations. (F) Geometric mean neutralization binding titers of serum from mice assigned to each cohort at day 0 (192, 171, and 73 days after the first, second, and third mRNA-LNP vaccinations but prior to nanoparticle or additional mRNA-LNP immunizations). (G) Non-baseline corrected geometric mean ELISA binding titers at the indicated days after immunization with mosaic-8b, mosaic-7, or WA1/BA.5 mRNA-LNP against indicated viral antigens. Compare with Figure 3E (baseline corrected geometric mean ELISA binding titers).

**Figure S4.**
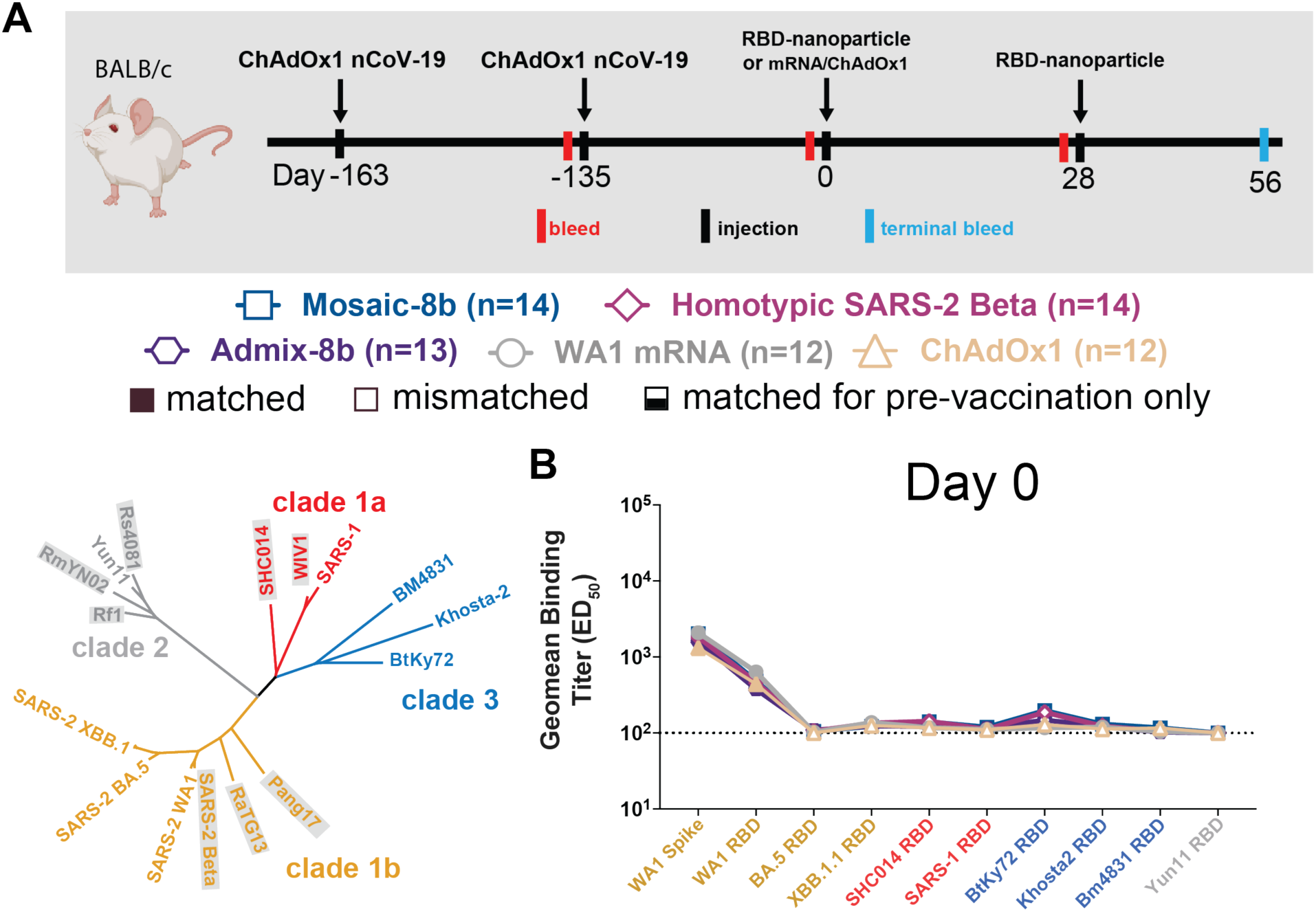
Cohorts of ChAdOx1 vaccinated mice showed no significant differences in midpoint ED_50_ titers at day 0, related to Figure 4. Geometric means of ED_50_ or ID_50_ values for all animals in each cohort are indicated by symbols connected by thick colored lines. Mean titers against indicated viral antigens or pseudoviruses were compared pairwise across immunization cohorts by Tukey’s multiple comparison test with the Geisser-Greenhouse correction (as calculated by GraphPad Prism). Significant differences between cohorts linked by vertical lines are indicated by asterisks: p<0.05 = *, p<0.01 = **, p<0.001 = ***, p<0.0001 = ****. (A) Top: Schematic of vaccination/immunization regimen. Mice were vaccinated at the indicated days prior to RBD-nanoparticle prime and boost immunizations at days 0 and 28 or mRNA-LNP or ChAdOx1 prime immunizations at day 0. Bottom: Phylogenetic tree of selected sarbecoviruses calculated using a Jukes-Cantor generic distance model using Geneious Prime® 2023.1.2 based on amino acid sequences of RBDs aligned using Clustal Omega.^97^ RBDs included in mosaic-8b are highlighted in gray rectangles. (B) Geometric mean ELISA binding titers of serum from mice assigned to each cohort at day 0 (163 and 134 days after the first and second ChAdOx1 vaccinations but prior to nanoparticle, mRNA-LNP, or additional ChAdOx1 immunizations). Binding titers are represented as mean ED_50_ values for serum IgG binding to RBD or spike proteins from the indicated sarbecovirus strains.

**Figure S5.**
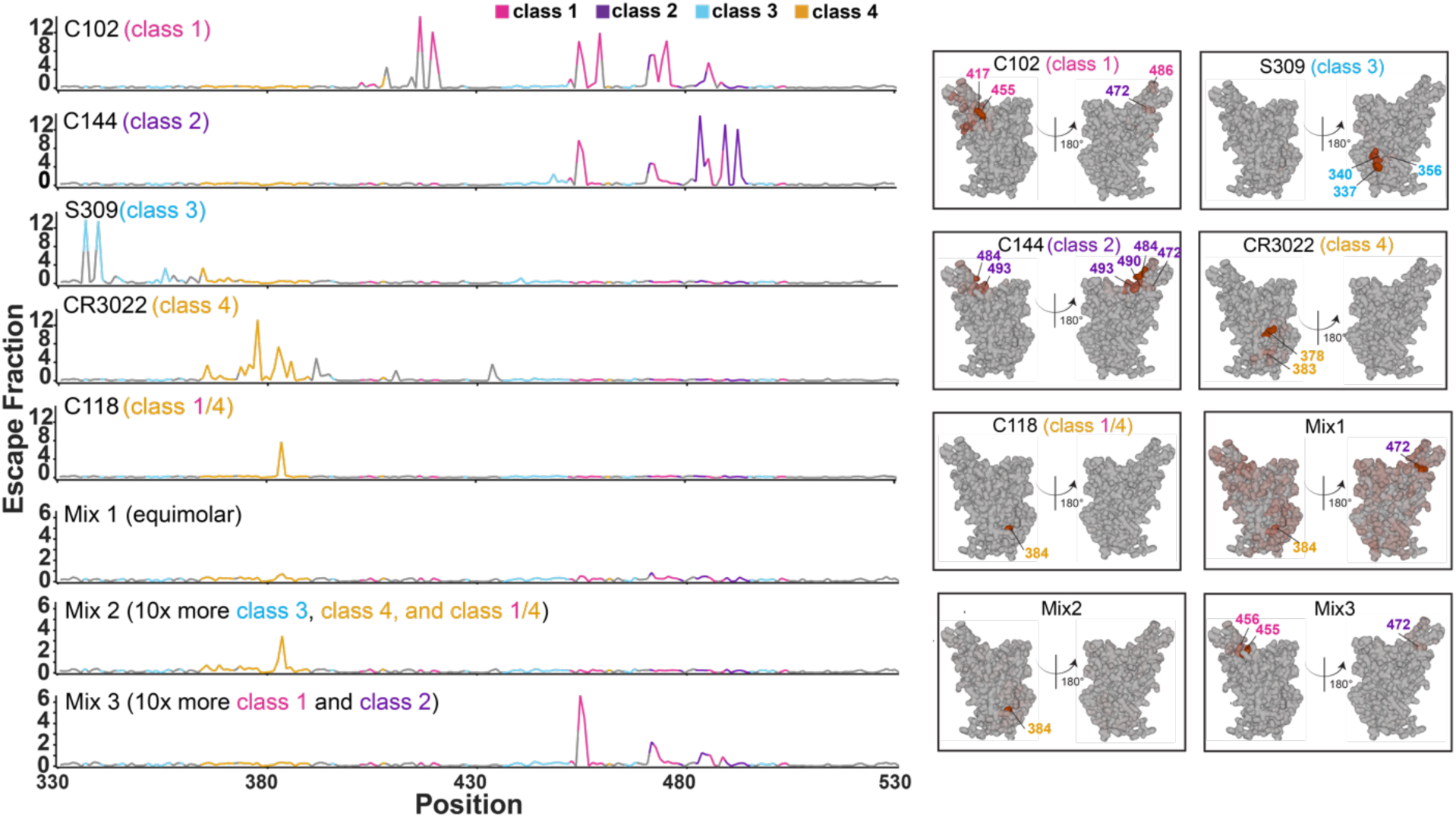
Comparison of DMS profiles of individual mAbs and mAb mixtures, related to Figure 5. Left: Line plots for DMS results from individual mAbs or the indicated mAb mixtures recognizing different RBD epitopes (epitopes defined in Figure 1B). C102, C118, C144, and S309 are mAbs that were derived from COVID-19 convalescent donors.^60,70^ CR3022 is an anti-SARS-CoV mAb that binds to SARS-2 RBD.^66^ Epitope assignments for these mAbs were previously described.^90^ DMS was conducted for the mAb reagents using a WA1 RBD library. The x-axis shows the RBD residue number and the y-axis shows the sum of the Ab escape of all mutations at a site (larger numbers indicating more Ab escape). Each line represents one antiserum with heavy lines showing the average across the n=3 sera in each group. Lines are colored differently for RBD epitopes within different classes^17^ (epitopes defined in Figure 1B); gray for residues not assigned to an epitope. Right: The average site-total Ab escape for the indicated mAbs and mAb mixtures (Mix 1 = equimolar mixture of all five mAbs; Mix 2 = 10-fold more S309, C118, and CR3022 than C102 and C144; Mix 3 = 10-fold more C102 and C144 than S309, C118, and CR3022) mapped to the surface of the WA1 RBD (PDB 6M0J). The locations of individual residues are highlighted on the RBD surfaces in colors corresponding to their epitope.

**Figure S6.**
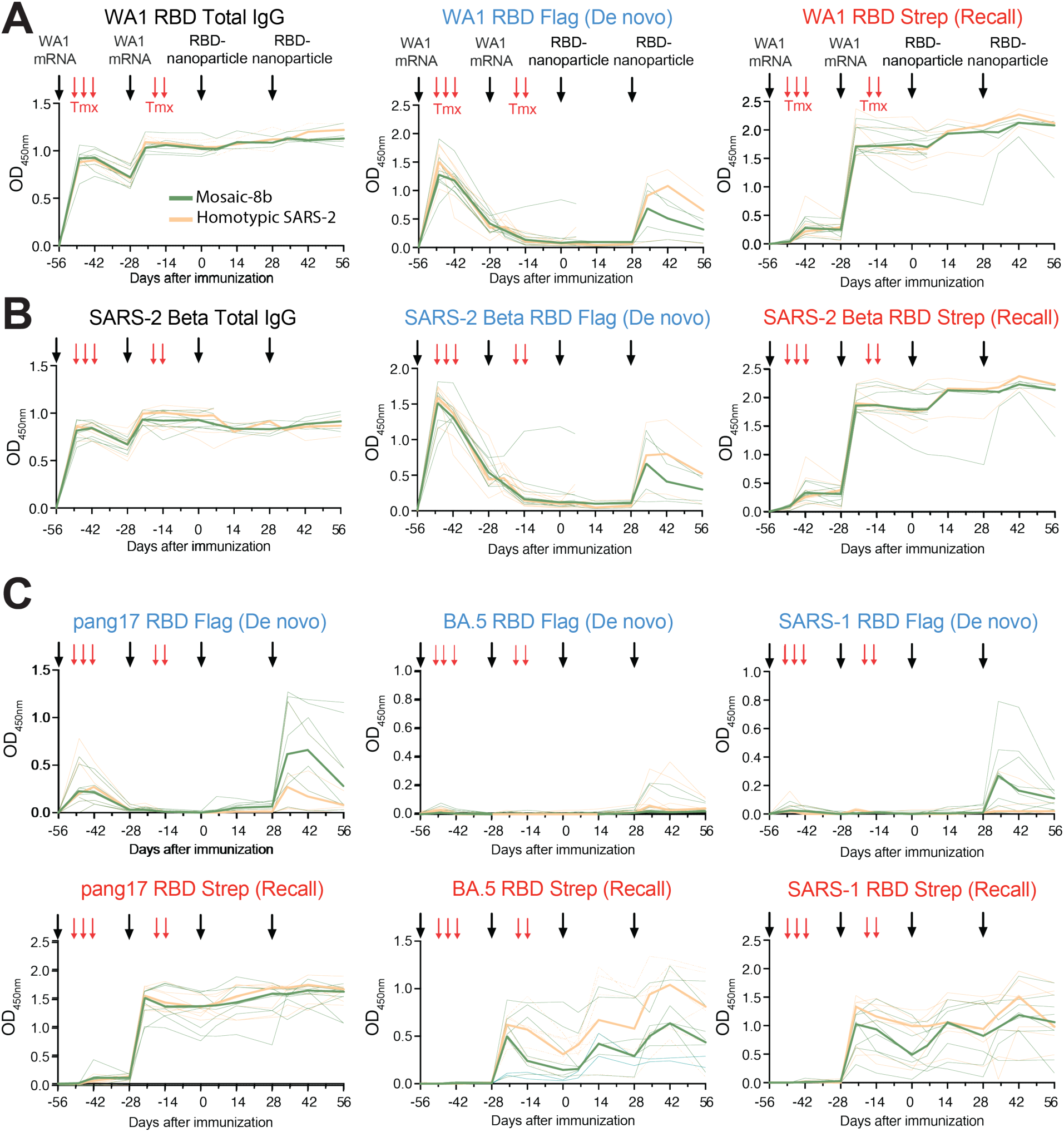
Time course of serum from fate mapping studies reveals differences in Ab responses before RBD-nanoparticle immunization, related to Figure 6. Time course of serum antibody responses. Flag^+^/de novo (blue) or Strep^+^/recall (red) Abs after immunization with RBD-nanoparticles. Red arrows indicate tamoxifen (Tmx) treatment in all panels; black arrows indicate vaccination with WA1 mRNA-LNP or the indicated RBD-nanoparticles, as denoted in the upper left panel. Mice immunized with mosaic-8b are indicated in green; mice immunized with homotypic SARS-2 Beta are indicated in orange. Individual mice are indicated by thin lines; median responses for groups are shown in thick lines. (A-B) Time course of levels of total IgG (left), Flag^+^/de novo Igκ (center), or Strep+/recall Igκ (right) Ab responses after immunization with mosaic-8b (green) or homotypic SARS-2 Beta (orange) RBD-nanoparticles measured against RBDs of (A) SARS-2 WA1, (B) SARS-2 Beta. (C) Time course of Flag^+^/de novo Igκ (top), or Strep+/recall Igκ (bottom) Ab responses after immunization with mosaic-8b (green) or homotypic SARS-2 Beta (orange) RBD-nanoparticles measured against RBDs of pang17, SARS-2 BA.5, or SARS-1. ELISA absorbance values (OD_450 nm_) are shown for serum samples diluted at 1:100 (the first dilution for endpoint titer ELISAs shown in Figure 6). Data up to day 0 are from the two independent experimental cohorts shown in Figure 6. Data from one of the experimental cohorts was collected up to day 56. For accurate comparisons, each datapoint within each plot is from samples analyzed together in one assay.

**Figure S7.**
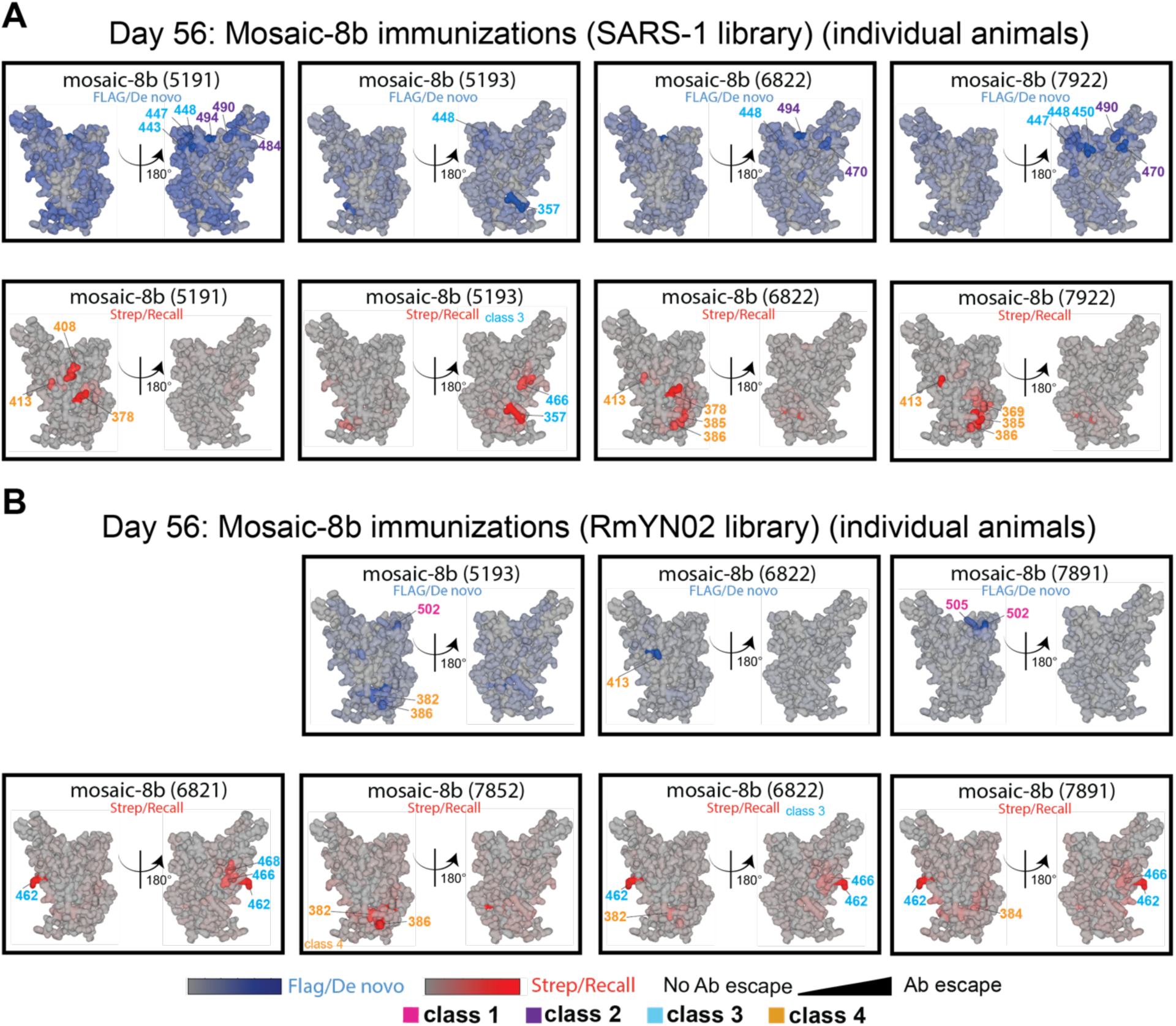
Individual animal DMS results, related to Figure 6. DMS analyses shown for individual animals for which we had sufficient sera and the sera exhibited detectable levels of Flag^+^ or Strep^+^ Abs. (A) DMS analyses for individual animals (indicated by 4-digit numbers) for compiled results shown in Figure 6F of day 56 serum from RBD-nanoparticle immunized mice using SARS-1 (antigenic distance score = 0.81) RBD mutant library. Ab binding sites are shaded according to degree of Ab escape, with blue for Flag/de novo responses and red for Strep/recall responses, on the surface of the WA1 RBD (PDB 6M0J). Comparisons are made for Flag/de novo and Strep/recall elicited by mosaic-8b and for Strep/recall elicited by homotypic SARS-2 (there were weak to no Flag/de novo responses after homotypic SARS-2 immunization). (B) DMS analyses for individual animals (indicated by 4-digit numbers) for compiled results shown in Figure 6F of day 56 serum from RBD-nanoparticle immunized mice using RmYN02 (antigenic distance score = 0.63) RBD mutant library. Ab binding sites are shaded according to degree of Ab escape, with blue for Flag/de novo responses and red for Strep/recall responses, on the surface of the WA1 RBD (PDB 6M0J). Comparisons are made for Flag/de novo and Strep/recall elicited by mosaic-8b and for Strep/recall elicited by homotypic SARS-2 (there were weak to no Flag/de novo responses after homotypic SARS-2 immunization). RmYN02 RBD is included on the mosaic-8b nanoparticle.

## Bibliography

1 Menachery, V. D., Yount, B. L., Debbink, K., Agnihothram, S., Gralinski, L. E., Plante, J. A., Graham, R. L., Scobey, T., Ge, X.-Y., Donaldson, E. F., Randell, S. H., Lanzavecchia, A., Marasco, W. A., Shi, Z.-L. & Baric, R. S. (2015). A SARS-like cluster of circulating bat coronaviruses shows potential for human emergence. Nature Medicine 21, 1508–1513.

2 Menachery, V. D., Yount, B. L., Sims, A. C., Debbink, K., Agnihothram, S. S., Gralinski, L. E., Graham, R. L., Scobey, T., Plante, J. A., Royal, S. R., Swanstrom, J., Sheahan, T. P., Pickles, R. J., Corti, D., Randell, S. H., Lanzavecchia, A., Marasco, W. A. & Baric, R. S. (2016). SARS-like WIV1-CoV poised for human emergence. Proceedings of the National Academy of Sciences 113, 3048–3053.

3 Zhou, H., Ji, J., Chen, X., Bi, Y., Li, J., Wang, Q., Hu, T., Song, H., Zhao, R., Chen, Y., Cui, M., Zhang, Y., Hughes, A. C., Holmes, E. C. & Shi, W. (2021). Identification of novel bat coronaviruses sheds light on the evolutionary origins of SARS-CoV-2 and related viruses. Cell 184, 4380–4391 e4314.

4 Davis, H. E., McCorkell, L., Vogel, J. M. & Topol, E. J. (2023). Long COVID: major findings, mechanisms and recommendations. Nature Reviews Microbiology 21, 133–146.

5 Planas, D., Bruel, T., Grzelak, L., Guivel-Benhassine, F., Staropoli, I., Porrot, F., Planchais, C., Buchrieser, J., Rajah, M. M., Bishop, E., Albert, M., Donati, F., Prot, M., Behillil, S., Enouf, V., Maquart, M., Smati-Lafarge, M., Varon, E., Schortgen, F., Yahyaoui, L., Gonzalez, M., De Seze, J., Pere, H., Veyer, D., Seve, A., Simon-Loriere, E., Fafi-Kremer, S., Stefic, K., Mouquet, H., Hocqueloux, L., van der Werf, S., Prazuck, T. & Schwartz, O. (2021). Sensitivity of infectious SARS-CoV-2 B.1.1.7 and B.1.351 variants to neutralizing antibodies. Nat Med 27, 917–924.

6 Washington, N. L., Gangavarapu, K., Zeller, M., Bolze, A., Cirulli, E. T., Schiabor Barrett, K. M., Larsen, B. B., Anderson, C., White, S., Cassens, T., Jacobs, S., Levan, G., Nguyen, J., Ramirez, J. M., 3rd, Rivera-Garcia, C., Sandoval, E., Wang, X., Wong, D., Spencer, E., Robles-Sikisaka, R., Kurzban, E., Hughes, L. D., Deng, X., Wang, C., Servellita, V., Valentine, H., De Hoff, P., Seaver, P., Sathe, S., Gietzen, K., Sickler, B., Antico, J., Hoon, K., Liu, J., Harding, A., Bakhtar, O., Basler, T., Austin, B., MacCannell, D., Isaksson, M., Febbo, P. G., Becker, D., Laurent, M., McDonald, E., Yeo, G. W., Knight, R., Laurent, L. C., de Feo, E., Worobey, M., Chiu, C. Y., Suchard, M. A., Lu, J. T., Lee, W. & Andersen, K. G. (2021). Emergence and rapid transmission of SARS-CoV-2 B.1.1.7 in the United States. Cell 184, 2587–2594 e2587.

7 Burki, T. K. (2021). Omicron variant and booster COVID-19 vaccines. The Lancet Respiratory Medicine.

8 Liu, L., Iketani, S., Guo, Y., Chan, J. F., Wang, M., Liu, L., Luo, Y., Chu, H., Huang, Y., Nair, M. S., Yu, J., Chik, K. K., Yuen, T. T., Yoon, C., To, K. K., Chen, H., Yin, M. T., Sobieszczyk, M. E., Huang, Y., Wang, H. H., Sheng, Z., Yuen, K. Y. & Ho, D. D. (2021). Striking Antibody Evasion Manifested by the Omicron Variant of SARS-CoV-2. Nature.

9 Kleanthous, H., Silverman, J. M., Makar, K. W., Yoon, I. K., Jackson, N. & Vaughn, D. W. (2021). Scientific rationale for developing potent RBD-based vaccines targeting COVID-19. NPJ Vaccines 6, 128.

10 Bowen, J. E., Addetia, A., Dang, H. V., Stewart, C., Brown, J. T., Sharkey, W. K., Sprouse, K. R., Walls, A. C., Mazzitelli, I. G., Logue, J. K., Franko, N. M., Czudnochowski, N., Powell, A. E., Dellota, E., Jr., Ahmed, K., Ansari, A. S., Cameroni, E., Gori, A., Bandera, A., Posavad, C. M., Dan, J. M., Zhang, Z., Weiskopf, D., Sette, A., Crotty, S., Iqbal, N. T., Corti, D., Geffner, J., Snell, G., Grifantini, R., Chu, H. Y. & Veesler, D. (2022). Omicron spike function and neutralizing activity elicited by a comprehensive panel of vaccines. Science 377, 890–894.

11 Starr, T. N., Czudnochowski, N., Liu, Z., Zatta, F., Park, Y. J., Addetia, A., Pinto, D., Beltramello, M., Hernandez, P., Greaney, A. J., Marzi, R., Glass, W. G., Zhang, I., Dingens, A. S., Bowen, J. E., Tortorici, M. A., Walls, A. C., Wojcechowskyj, J. A., De Marco, A., Rosen, L. E., Zhou, J., Montiel-Ruiz, M., Kaiser, H., Dillen, J. R., Tucker, H., Bassi, J., Silacci-Fregni, C., Housley, M. P., di Iulio, J., Lombardo, G., Agostini, M., Sprugasci, N., Culap, K., Jaconi, S., Meury, M., Dellota, E., Jr., Abdelnabi, R., Foo, S. C., Cameroni, E., Stumpf, S., Croll, T. I., Nix, J. C., Havenar-Daughton, C., Piccoli, L., Benigni, F., Neyts, J., Telenti, A., Lempp, F. A., Pizzuto, M. S., Chodera, J. D., Hebner, C. M., Virgin, H. W., Whelan, S. P. J., Veesler, D., Corti, D., Bloom, J. D. & Snell, G. (2021). SARS-CoV-2 RBD antibodies that maximize breadth and resistance to escape. Nature 597, 97–102.

12 Khoury, D. S., Docken, S. S., Subbarao, K., Kent, S. J., Davenport, M. P. & Cromer, D. (2023). Predicting the efficacy of variant-modified COVID-19 vaccine boosters. Nat Med 29, 574–578.

13 Cohen, A. A., Gnanapragasam, P. N. P., Lee, Y. E., Hoffman, P. R., Ou, S., Kakutani, L. M., Keeffe, J. R., Wu, H. J., Howarth, M., West, A. P., Barnes, C. O., Nussenzweig, M. C. & Bjorkman, P. J. (2021). Mosaic nanoparticles elicit cross-reactive immune responses to zoonotic coronaviruses in mice. Science 371, 735–741.

14 Cohen, A. A., van Doremalen, N., Greaney, A. J., Andersen, H., Sharma, A., Starr, T. N., Keeffe, J. R., Fan, C., Schulz, J. E., Gnanapragasam, P. N. P., Kakutani, L. M., West, A. P., Jr., Saturday, G., Lee, Y. E., Gao, H., Jette, C. A., Lewis, M. G., Tan, T. K., Townsend, A. R., Bloom, J. D., Munster, V. J. & Bjorkman, P. J. (2022). Mosaic RBD nanoparticles protect against challenge by diverse sarbecoviruses in animal models. Science 377, eabq0839.

15 Dong, W., Mead, H., Tian, L., Park, J. G., Garcia, J. I., Jaramillo, S., Barr, T., Kollath, D. S., Coyne, V. K., Stone, N. E., Jones, A., Zhang, J., Li, A., Wang, L. S., Milanes-Yearsley, M., Torrelles, J. B., Martinez-Sobrido, L., Keim, P. S., Barker, B. M., Caligiuri, M. A. & Yu, J. (2022). The K18-Human ACE2 Transgenic Mouse Model Recapitulates Non-severe and Severe COVID-19 in Response to an Infectious Dose of the SARS-CoV-2 Virus. J Virol 96, e0096421.

16 Starr, T. N., Greaney, A. J., Hilton, S. K., Ellis, D., Crawford, K. H. D., Dingens, A. S., Navarro, M. J., Bowen, J. E., Tortorici, M. A., Walls, A. C., King, N. P., Veesler, D. & Bloom, J. D. (2020). Deep Mutational Scanning of SARS-CoV-2 Receptor Binding Domain Reveals Constraints on Folding and ACE2 Binding. Cell 182, 1295–1310.e1220.

17 Barnes, C. O., Jette, C. A., Abernathy, M. E., Dam, K.-M. A., Esswein, S. R., Gristick, H. A. , Malyutin, A. G., Sharaf, N. G., Huey-Tubman, K. E., Lee, Y. E., Robbiani, D. F., Nussenzweig, M. C., West, A. P. & Bjorkman, P. J. (2020). SARS-CoV-2 neutralizing antibody structures inform therapeutic strategies. Nature 588, 682–687.

18 Fan, C., Cohen, A. A., Park, M., Hung, A. F., Keeffe, J. R., Gnanapragasam, P. N. P., Lee, Y. E., Gao, H., Kakutani, L. M., Wu, Z., Kleanthous, H., Malecek, K. E., Williams, J. & Bjorkman, P. J. (2022). Neutralizing monoclonal antibodies elicited by mosaic RBD nanoparticles bind conserved sarbecovirus epitopes. Immunity 55, 2419–2435 e2410.

19 Francis, T., Jr. (1960). On the Doctrine of Original Antigenic Sin. Proceedings of the American Philosophical Society 104, 572–578.

20 Cobey, S. & Hensley, S. E. (2017). Immune history and influenza virus susceptibility. Current opinion in virology 22, 105–111.

21 Schiepers, A., van ’t Wout, M. F. L., Greaney, A. J., Zang, T., Muramatsu, H., Lin, P. J. C., Tam, Y. K., Mesin, L., Starr, T. N., Bieniasz, P. D., Pardi, N., Bloom, J. D. & Victora, G. D. (2023). Molecular fate-mapping of serum antibody responses to repeat immunization. Nature 615, 482–489.

22 Brune, K. D., Leneghan, D. B., Brian, I. J., Ishizuka, A. S., Bachmann, M. F., Draper, S. J., Biswas, S. & Howarth, M. (2016). Plug-and-Display: decoration of Virus-Like Particles via isopeptide bonds for modular immunization. Scientific reports 6, 19234.

23 Zakeri, B., Fierer, J. O., Celik, E., Chittock, E. C., Schwarz-Linek, U., Moy, V. T. & Howarth, M. (2012). Peptide tag forming a rapid covalent bond to a protein, through engineering a bacterial adhesin. Proc Natl Acad Sci U S A 109, E690–697.

24 Keeble, A. H., Turkki, P., Stokes, S., Khairil Anuar, I. N. A., Rahikainen, R., Hytönen, V. P. & Howarth, M. (2019). Approaching infinite affinity through engineering of peptide– protein interaction. Proceedings of the National Academy of Sciences 116, 26523–26533.

25 Goldblatt, D., Alter, G., Crotty, S. & Plotkin, S. A. (2022). Correlates of protection against SARS-CoV-2 infection and COVID-19 disease. Immunol Rev 310, 6–26.

26 Dross, S., Fredericks, M. N., Maughan, M. D., UlrichLewis, J. T., Lewis, T. B., Dong, Y., Fuller, J. T., Wangari, S., Cohen, A. A., Bjorkman, P. J., Ramsingh, A. I., Pata, J., Erasmus, J., Bagley, K. C. & Fuller, D. H. in Conference on Retroviruses and Opportunistic Infections (Poster Session-D7, SARS-CoV-2 Vaccines and Prevention, Seattle, WA, 2023).

27 Erasmus, J. H., Archer, J., Randall, S., Pata, J. D., Ramsingh, A. I. & Fuller, D. H. (2024). A chimeric SARS-CoV-2 self-amplifying RNA vaccine induces robust receptor binding domain-focused antibody and T cell responses in naive and pre-immune mice. Ms. submitted.

28 Erasmus, J. H., Khandhar, A. P., O’Connor, M. A., Walls, A. C., Hemann, E. A., Murapa, P., Archer, J., Leventhal, S., Fuller, J. T., Lewis, T. B., Draves, K. E., Randall, S., Guerriero, K. A., Duthie, M. S., Carter, D., Reed, S. G., Hawman, D. W., Feldmann, H., Gale, M., Jr., Veesler, D., Berglund, P. & Fuller, D. H. (2020). An Alphavirus-derived replicon RNA vaccine induces SARS-CoV-2 neutralizing antibody and T cell responses in mice and nonhuman primates. Science translational medicine 12.

29 Franzese, M., Coppola, L., Silva, R., Santini, S. A., Cinquanta, L., Ottomano, C., Salvatore, M. & Incoronato, M. (2022). SARS-CoV-2 antibody responses before and after a third dose of the BNT162b2 vaccine in Italian healthcare workers aged </=60 years: One year of surveillance. Front Immunol 13, 947187.

30 Shen, X. (2022). Boosting immunity to Omicron. Nat Med 28, 445–446.

31 Zhang, Z., Mateus, J., Coelho, C. H., Dan, J. M., Moderbacher, C. R., Galvez, R. I., Cortes, F. H., Grifoni, A., Tarke, A., Chang, J., Escarrega, E. A., Kim, C., Goodwin, B., Bloom, N. I., Frazier, A., Weiskopf, D., Sette, A. & Crotty, S. (2022). Humoral and cellular immune memory to four COVID-19 vaccines. Cell 185, 2434–2451 e2417.

32 Greaney, A. J., Starr, T. N., Barnes, C. O., Weisblum, Y., Schmidt, F., Caskey, M., Gaebler, C., Cho, A., Agudelo, M., Finkin, S., Wang, Z., Poston, D., Muecksch, F., Hatziioannou, T., Bieniasz, P. D., Robbiani, D. F., Nussenzweig, M. C., Bjorkman, P. J. & Bloom, J. D. (2021). Mapping mutations to the SARS-CoV-2 RBD that escape binding by different classes of antibodies. Nat Commun 12, 4196.

33 Greaney, A. J., Starr, T. N., Gilchuk, P., Zost, S. J., Binshtein, E., Loes, A. N., Hilton, S. K., Huddleston, J., Eguia, R., Crawford, K. H. D., Dingens, A. S., Nargi, R. S., Sutton, R. E., Suryadevara, N., Rothlauf, P. W., Liu, Z., Whelan, S. P. J., Carnahan, R. H., Crowe, J. E., Jr. & Bloom, J. D. (2021). Complete Mapping of Mutations to the SARS-CoV-2 Spike Receptor-Binding Domain that Escape Antibody Recognition. Cell Host Microbe 29, 44–57 e49.

34 Shinnakasu, R., Inoue, T., Kometani, K., Moriyama, S., Adachi, Y., Nakayama, M., Takahashi, Y., Fukuyama, H., Okada, T. & Kurosaki, T. (2016). Regulated selection of germinal-center cells into the memory B cell compartment. Nat Immunol 17, 861–869.

35 Mesin, L., Schiepers, A., Ersching, J., Barbulescu, A., Cavazzoni, C. B., Angelini, A., Okada, T., Kurosaki, T. & Victora, G. D. (2020). Restricted Clonality and Limited Germinal Center Reentry Characterize Memory B Cell Reactivation by Boosting. Cell 180, 92–106 e111.

36 Heyman, B. (2000). Regulation of antibody responses via antibodies, complement, and Fc receptors. Annu Rev Immunol 18, 709–737.

37 McNamara, H. A., Idris, A. H., Sutton, H. J., Vistein, R., Flynn, B. J., Cai, Y., Wiehe, K., Lyke, K. E., Chatterjee, D., Kc, N., Chakravarty, S., Lee Sim, B. K., Hoffman, S. L., Bonsignori, M., Seder, R. A. & Cockburn, I. A. (2020). Antibody Feedback Limits the Expansion of B Cell Responses to Malaria Vaccination but Drives Diversification of the Humoral Response. Cell Host Microbe 28, 572–585 e577.

38 Tas, J. M. J., Koo, J. H., Lin, Y. C., Xie, Z., Steichen, J. M., Jackson, A. M., Hauser, B. M., Wang, X., Cottrell, C. A., Torres, J. L., Warner, J. E., Kirsch, K. H., Weldon, S. R., Groschel, B., Nogal, B., Ozorowski, G., Bangaru, S., Phelps, N., Adachi, Y., Eskandarzadeh, S., Kubitz, M., Burton, D. R., Lingwood, D., Schmidt, A. G., Nair, U., Ward, A. B., Schief, W. R. & Batista, F. D. (2022). Antibodies from primary humoral responses modulate the recruitment of naive B cells during secondary responses. Immunity 55, 1856–1871 e1856.

39 Schaefer-Babajew, D., Wang, Z., Muecksch, F., Cho, A., Loewe, M., Cipolla, M., Raspe, R., Johnson, B., Canis, M., DaSilva, J., Ramos, V., Turroja, M., Millard, K. G., Schmidt, F., Witte, L., Dizon, J., Shimeliovich, I., Yao, K. H., Oliveira, T. Y., Gazumyan, A., Gaebler, C., Bieniasz, P. D., Hatziioannou, T., Caskey, M. & Nussenzweig, M. C. (2022). Antibody feedback regulates immune memory after SARS-CoV-2 mRNA vaccination. Nature.

40 Schiepers, A., van ‘t Wout, M. F. L., Hobbs, A., Mesin, L. & Victora, G. D. (2023). Opposing effects of pre-existing antibody and memory T cell help on the dynamics of recall germinal centers. bioRxiv.

41 Bergeri, I., Whelan, M. G., Ware, H., Subissi, L., Nardone, A., Lewis, H. C., Li, Z., Ma, X., Valenciano, M., Cheng, B., Al Ariqi, L., Rashidian, A., Okeibunor, J., Azim, T., Wijesinghe, P., Le, L. V., Vaughan, A., Pebody, R., Vicari, A., Yan, T., Yanes-Lane, M., Cao, C., Clifton, D. A., Cheng, M. P., Papenburg, J., Buckeridge, D., Bobrovitz, N., Arora, R. K., Van Kerkhove, M. D. & Unity Studies Collaborator, G. (2022). Global SARS-CoV-2 seroprevalence from January 2020 to April 2022: A systematic review and meta-analysis of standardized population-based studies. PLoS Med 19, e1004107.

42 Azami, M., Moradi, Y., Moradkhani, A. & Aghaei, A. (2022). SARS-CoV-2 seroprevalence around the world: an updated systematic review and meta-analysis. Eur J Med Res 27, 81.

43 Corbett, K. S., Nason, M. C., Flach, B., Gagne, M., O’Connell, S., Johnston, T. S., Shah, S. N., Edara, V. V., Floyd, K., Lai, L., McDanal, C., Francica, J. R., Flynn, B., Wu, K., Choi, A., Koch, M., Abiona, O. M., Werner, A. P., Moliva, J. I., Andrew, S. F., Donaldson, M. M., Fintzi, J., Flebbe, D. R., Lamb, E., Noe, A. T., Nurmukhambetova, S. T., Provost, S. J., Cook, A., Dodson, A., Faudree, A., Greenhouse, J., Kar, S., Pessaint, L., Porto, M., Steingrebe, K., Valentin, D., Zouantcha, S., Bock, K. W., Minai, M., Nagata, B. M., van de Wetering, R., Boyoglu-Barnum, S., Leung, K., Shi, W., Yang, E. S., Zhang, Y., Todd, J. M., Wang, L., Alvarado, G. S., Andersen, H., Foulds, K. E., Edwards, D. K., Mascola, J. R., Moore, I. N., Lewis, M. G., Carfi, A., Montefiori, D., Suthar, M. S., McDermott, A., Roederer, M., Sullivan, N. J., Douek, D. C., Graham, B. S. & Seder, R. A. (2021). Immune correlates of protection by mRNA-1273 vaccine against SARS-CoV-2 in nonhuman primates. Science 373, eabj0299.

44 Gilbert, P. B., Montefiori, D. C., McDermott, A. B., Fong, Y., Benkeser, D., Deng, W., Zhou, H., Houchens, C. R., Martins, K., Jayashankar, L., Castellino, F., Flach, B., Lin, B. C., O’Connell, S., McDanal, C., Eaton, A., Sarzotti-Kelsoe, M., Lu, Y., Yu, C., Borate, B., van der Laan, L. W. P., Hejazi, N. S., Huynh, C., Miller, J., El Sahly, H. M., Baden, L. R., Baron, M., De La Cruz, L., Gay, C., Kalams, S., Kelley, C. F., Andrasik, M. P., Kublin, J. G., Corey, L., Neuzil, K. M., Carpp, L. N., Pajon, R., Follmann, D., Donis, R. O., Koup, R. A., Immune Assays Team section, s., Moderna, I. T. s. s., Coronavirus Vaccine Prevention Network /Coronavirus Efficacy Team section, s. & United States Government /Co, V. P. N. B. T. s. s. (2022). Immune correlates analysis of the mRNA-1273 COVID-19 vaccine efficacy clinical trial. Science 375, 43–50.

45 Fong, Y., Huang, Y., Benkeser, D., Carpp, L. N., Anez, G., Woo, W., McGarry, A., Dunkle, L. M., Cho, I., Houchens, C. R., Martins, K., Jayashankar, L., Castellino, F., Petropoulos, C. J., Leith, A., Haugaard, D., Webb, B., Lu, Y., Yu, C., Borate, B., van der Laan, L. W. P., Hejazi, N. S., Randhawa, A. K., Andrasik, M. P., Kublin, J. G., Hutter, J., Keshtkar-Jahromi, M., Beresnev, T. H., Corey, L., Neuzil, K. M., Follmann, D., Ake, J. A., Gay, C. L., Kotloff, K. L., Koup, R. A., Donis, R. O., Gilbert, P. B., Immune Assays, T., Coronavirus Vaccine Prevention Network /nCo, V. P. I., Study, T. & United States Government /Co, V. P. N. B. T. (2023). Immune correlates analysis of the PREVENT-19 COVID-19 vaccine efficacy clinical trial. Nat Commun 14, 331.

46 Joyce, M. G., Chen, W. H., Sankhala, R. S., Hajduczki, A., Thomas, P. V., Choe, M., Martinez, E. J., Chang, W. C., Peterson, C. E., Morrison, E. B., Smith, C., Chen, R. E., Ahmed, A., Wieczorek, L., Anderson, A., Case, J. B., Li, Y., Oertel, T., Rosado, L., Ganesh, A., Whalen, C., Carmen, J. M., Mendez-Rivera, L., Karch, C. P., Gohain, N., Villar, Z., McCurdy, D., Beck, Z., Kim, J., Shrivastava, S., Jobe, O., Dussupt, V., Molnar, S., Tran, U., Kannadka, C. B., Soman, S., Kuklis, C., Zemil, M., Khanh, H., Wu, W., Cole, M. A., Duso, D. K., Kummer, L. W., Lang, T. J., Muncil, S. E., Currier, J. R., Krebs, S. J., Polonis, V. R., Rajan, S., McTamney, P. M., Esser, M. T., Reiley, W. W., Rolland, M., de Val, N., Diamond, M. S., Gromowski, G. D., Matyas, G. R., Rao, M., Michael, N. L. & Modjarrad, K. (2021). SARS-CoV-2 ferritin nanoparticle vaccines elicit broad SARS coronavirus immunogenicity. Cell reports 37, 110143.

47 Saunders, K. O., Lee, E., Parks, R., Martinez, D. R., Li, D., Chen, H., Edwards, R. J., Gobeil, S., Barr, M., Mansouri, K., Alam, S. M., Sutherland, L. L., Cai, F., Sanzone, A. M., Berry, M., Manne, K., Bock, K. W., Minai, M., Nagata, B. M., Kapingidza, A. B., Azoitei, M., Tse, L. V., Scobey, T. D., Spreng, R. L., Rountree, R. W., DeMarco, C. T., Denny, T. N., Woods, C. W., Petzold, E. W., Tang, J., Oguin, T. H., 3rd, Sempowski, G. D., Gagne, M., Douek, D. C., Tomai, M. A., Fox, C. B., Seder, R., Wiehe, K., Weissman, D., Pardi, N., Golding, H., Khurana, S., Acharya, P., Andersen, H., Lewis, M. G., Moore, I. N., Montefiori, D. C., Baric, R. S. & Haynes, B. F. (2021). Neutralizing antibody vaccine for pandemic and pre-emergent coronaviruses. Nature 594, 553–559.

48 Tan, T. K., Rijal, P., Rahikainen, R., Keeble, A. H., Schimanski, L., Hussain, S., Harvey, R., Hayes, J. W. P., Edwards, J. C., McLean, R. K., Martini, V., Pedrera, M., Thakur, N., Conceicao, C., Dietrich, I., Shelton, H., Ludi, A., Wilsden, G., Browning, C., Zagrajek, A. K., Bialy, D., Bhat, S., Stevenson-Leggett, P., Hollinghurst, P., Tully, M., Moffat, K., Chiu, C., Waters, R., Gray, A., Azhar, M., Mioulet, V., Newman, J., Asfor, A. S., Burman, A., Crossley, S., Hammond, J. A., Tchilian, E., Charleston, B., Bailey, D., Tuthill, T. J., Graham, S. P., Duyvesteyn, H. M. E., Malinauskas, T., Huo, J., Tree, J. A., Buttigieg, K. R., Owens, R. J., Carroll, M. W., Daniels, R. S., McCauley, J. W., Stuart, D. I., Huang, K. A., Howarth, M. & Townsend, A. R. (2021). A COVID-19 vaccine candidate using SpyCatcher multimerization of the SARS-CoV-2 spike protein receptor-binding domain induces potent neutralising antibody responses. Nat Commun 12, 542.

49 Powell, A. E., Zhang, K., Sanyal, M., Tang, S., Weidenbacher, P. A., Li, S., Pham, T. D., Pak, J. E., Chiu, W. & Kim, P. S. (2021). A Single Immunization with Spike-Functionalized Ferritin Vaccines Elicits Neutralizing Antibody Responses against SARS-CoV-2 in Mice. ACS Cent Sci 7, 183–199.

50 Heath, P. T., Galiza, E. P., Baxter, D. N., Boffito, M., Browne, D., Burns, F., Chadwick, D. R., Clark, R., Cosgrove, C., Galloway, J., Goodman, A. L., Heer, A., Higham, A., Iyengar, S., Jamal, A., Jeanes, C., Kalra, P. A., Kyriakidou, C., McAuley, D. F., Meyrick, A., Minassian, A. M., Minton, J., Moore, P., Munsoor, I., Nicholls, H., Osanlou, O., Packham, J., Pretswell, C. H., San Francisco Ramos, A., Saralaya, D., Sheridan, R. P., Smith, R., Soiza, R. L., Swift, P. A., Thomson, E. C., Turner, J., Viljoen, M. E., Albert, G., Cho, I., Dubovsky, F., Glenn, G., Rivers, J., Robertson, A., Smith, K., Toback, S. & nCo, V. S. G. (2021). Safety and Efficacy of NVX-CoV2373 Covid-19 Vaccine. N Engl J Med 385, 1172–1183.

51 Wang, W., Huang, B., Zhu, Y., Tan, W. & Zhu, M. (2021). Ferritin nanoparticle-based SARS-CoV-2 RBD vaccine induces a persistent antibody response and long-term memory in mice. Cell Mol Immunol 18, 749–751.

52 Ma, X., Zou, F., Yu, F., Li, R., Yuan, Y., Zhang, Y., Zhang, X., Deng, J., Chen, T., Song, Z., Qiao, Y., Zhan, Y., Liu, J., Zhang, J., Zhang, X., Peng, Z., Li, Y., Lin, Y., Liang, L., Wang, G., Chen, Y., Chen, Q., Pan, T., He, X. & Zhang, H. (2020). Nanoparticle Vaccines Based on the Receptor Binding Domain (RBD) and Heptad Repeat (HR) of SARS-CoV-2 Elicit Robust Protective Immune Responses. Immunity 53, 1315–1330.e1319.

53 Geng, Q., Tai, W., Baxter, V. K., Shi, J., Wan, Y., Zhang, X., Montgomery, S. A., Taft-Benz, S. A., Anderson, E. J., Knight, A. C., Dinnon, K. H., 3rd, Leist, S. R., Baric, R. S., Shang, J., Hong, S. W., Drelich, A., Tseng, C. K., Jenkins, M., Heise, M., Du, L. & Li, F. (2021). Novel virus-like nanoparticle vaccine effectively protects animal model from SARS-CoV-2 infection. PLoS Pathog 17, e1009897.

54 Kang, Y. F., Sun, C., Zhuang, Z., Yuan, R. Y., Zheng, Q., Li, J. P., Zhou, P. P., Chen, X. C., Liu, Z., Zhang, X., Yu, X. H., Kong, X. W., Zhu, Q. Y., Zhong, Q., Xu, M., Zhong, N. S., Zeng, Y. X., Feng, G. K., Ke, C., Zhao, J. C. & Zeng, M. S. (2021). Rapid Development of SARS-CoV-2 Spike Protein Receptor-Binding Domain Self-Assembled Nanoparticle Vaccine Candidates. ACS Nano 15, 2738–2752.

55 Walls, A. C., Fiala, B., Schafer, A., Wrenn, S., Pham, M. N., Murphy, M., Tse, L. V., Shehata, L., O’Connor, M. A., Chen, C., Navarro, M. J., Miranda, M. C., Pettie, D., Ravichandran, R., Kraft, J. C., Ogohara, C., Palser, A., Chalk, S., Lee, E. C., Guerriero, K., Kepl, E., Chow, C. M., Sydeman, C., Hodge, E. A., Brown, B., Fuller, J. T., Dinnon, K. H., 3rd, Gralinski, L. E., Leist, S. R., Gully, K. L., Lewis, T. B., Guttman, M., Chu, H. Y., Lee, K. K., Fuller, D. H., Baric, R. S., Kellam, P., Carter, L., Pepper, M., Sheahan, T. P., Veesler, D. & King, N. P. (2020). Elicitation of Potent Neutralizing Antibody Responses by Designed Protein Nanoparticle Vaccines for SARS-CoV-2. Cell 183, 1367–1382 e1317.

56 Li, D., Martinez, D. R., Schafer, A., Chen, H., Barr, M., Sutherland, L. L., Lee, E., Parks, R., Mielke, D., Edwards, W., Newman, A., Bock, K. W., Minai, M., Nagata, B. M., Gagne, M., Douek, D., DeMarco, C. T., Denny, T. N., Oguin, T. H., Brown, A., Rountree, W., Wang, Y., Mansouri, K., Edwards, R. J., Ferrari, G., Sempowski, G. D., Eaton, A., Tang, J., Cain, D. W., Santra, S., Pardi, N., Weissman, D., Tomai, M., Fox, C., Moore, I. N., Andersen, H., Lewis, M. G., Golding, H., Khurana, S., Seder, R., Baric, R. S., Montefiori, D. C., Saunders, K. O. & Haynes, B. F. (2022). Breadth of SARS-CoV-2 Neutralization and Protection Induced by a Nanoparticle Vaccine. Nat Comm 13, 6309.

57 Dunkle, L. M., Kotloff, K. L., Gay, C. L., Anez, G., Adelglass, J. M., Barrat Hernandez, A. Q., Harper, W. L., Duncanson, D. M., McArthur, M. A., Florescu, D. F., McClelland, R. S., Garcia-Fragoso, V., Riesenberg, R. A., Musante, D. B., Fried, D. L., Safirstein, B. E., McKenzie, M., Jeanfreau, R. J., Kingsley, J. K., Henderson, J. A., Lane, D. C., Ruiz-Palacios, G. M., Corey, L., Neuzil, K. M., Coombs, R. W., Greninger, A. L., Hutter, J., Ake, J. A., Smith, K., Woo, W., Cho, I., Glenn, G. M., Dubovsky, F. & nCo, V. S. G. (2022). Efficacy and Safety of NVX-CoV2373 in Adults in the United States and Mexico. N Engl J Med 386, 531–543.

58 Jette, C. A., Cohen, A. A., Gnanapragasam, P. N. P., Muecksch, F., Lee, Y. E., Huey-Tubman, K. E., Schmidt, F., Hatziioannou, T., Bieniasz, P. D., Nussenzweig, M. C., West, A. P., Keeffe, J. R., Bjorkman, P. J. & Barnes, C. O. (2021). Broad cross-reactivity across sarbecoviruses exhibited by a subset of COVID-19 donor-derived neutralizing antibodies. Cell reports 36, 109760.

59. Liu, H., Wu, N. C., Yuan, M., Bangaru, S., Torres, J. L., Caniels, T. G., van Schooten, J., Zhu, X., Lee, C.-C. D., Brouwer, P. J. M., van Gils, M. J., Sanders, R. W., Ward, A. B. & Wilson, I. A. (2020). Cross-Neutralization of a SARS-CoV-2 Antibody to a Functionally Conserved Site Is Mediated by Avidity. Immunity 53, 1272–1280.e1275.

60 Pinto, D., Park, Y.-J., Beltramello, M., Walls, A. C., Tortorici, M. A., Bianchi, S., Jaconi, S., Culap, K., Zatta, F., De Marco, A., Peter, A., Guarino, B., Spreafico, R., Cameroni, E., Case, J. B., Chen, R. E., Havenar-Daughton, C., Snell, G., Telenti, A., Virgin, H. W., Lanzavecchia, A., Diamond, M. S., Fink, K., Veesler, D. & Corti, D. (2020). Cross-neutralization of SARS-CoV-2 by a human monoclonal SARS-CoV antibody. Nature 583, 290–295.

61 Sheward, D. J., Pushparaj, P., Das, H., Kim, C., Kim, S., Hanke, L., Dyrdak, R., McInerney, G., Albert, J., Murrell, B., Karlsson Hedestam, G. B. & Hällberg, B. M. (2022). Structural basis of Omicron neutralization by affinity-matured public antibodies. bioRxiv.

62 Burnett, D. L., Jackson, K. J. L., Langley, D. B., Aggrawal, A., Stella, A. O., Johansen, M. D., Balachandran, H., Lenthall, H., Rouet, R., Walker, G., Saunders, B. M., Singh, M., Li, H., Henry, J. Y., Jackson, J., Stewart, A. G., Witthauer, F., Spence, M. A., Hansbro, N. G., Jackson, C., Schofield, P., Milthorpe, C., Martinello, M., Schulz, S. R., Roth, E., Kelleher, A., Emery, S., Britton, W. J., Rawlinson, W. D., Karl, R., Schafer, S., Winkler, T. H., Brink, R., Bull, R. A., Hansbro, P. M., Jack, H. M., Turville, S., Christ, D. & Goodnow, C. C. (2021). Immunizations with diverse sarbecovirus receptor-binding domains elicit SARS-CoV-2 neutralizing antibodies against a conserved site of vulnerability. Immunity 54, 2908–2921 e2906.

63 Westendorf, K., Žentelis, S., Wang, L., Foster, D., Vaillancourt, P., Wiggin, M., Lovett, E., van der Lee, R., Hendle, J., Pustilnik, A., Sauder, J. M., Kraft, L., Hwang, Y., Siegel, R. W., Chen, J., Heinz, B. A., Higgs, R. E., Kallewaard, N. L., Jepson, K., Goya, R., Smith, M. A., Collins, D. W., Pellacani, D., Xiang, P., de Puyraimond, V., Ricicova, M., Devorkin, L., Pritchard, C., O’Neill, A., Dalal, K., Panwar, P., Dhupar, H., Garces, F. A., Cohen, C. A., Dye, J. M., Huie, K. E., Badger, C. V., Kobasa, D., Audet, J., Freitas, J. J., Hassanali, S., Hughes, I., Munoz, L., Palma, H. C., Ramamurthy, B., Cross, R. W., Geisbert, T. W., Menacherry, V., Lokugamage, K., Borisevich, V., Lanz, I., Anderson, L., Sipahimalani, P., Corbett, K. S., Yang, E. S., Zhang, Y., Shi, W., Zhou, T., Choe, M., Misasi, J., Kwong, P. D., Sullivan, N. J., Graham, B. S., Fernandez, T. L., Hansen, C. L., Falconer, E., Mascola, J. R., Jones, B. E. & Barnhart, B. C. (2022). LY-CoV1404 (bebtelovimab) potently neutralizes SARS-CoV-2 variants. Cell reports.

64 Greaney, A. J., Loes, A. N., Crawford, K. H. D., Starr, T. N., Malone, K. D., Chu, H. Y. & Bloom, J. D. (2021). Comprehensive mapping of mutations in the SARS-CoV-2 receptor-binding domain that affect recognition by polyclonal human plasma antibodies. Cell Host Microbe 29, 463–476 e466.

65 Piccoli, L., Park, Y.-J., Tortorici, M. A., Czudnochowski, N., Walls, A. C., Beltramello, M., Silacci-Fregni, C., Pinto, D., Rosen, L. E., Bowen, J. E., Acton, O. J., Jaconi, S., Guarino, B., Minola, A., Zatta, F., Sprugasci, N., Bassi, J., Peter, A., De Marco, A., Nix, J. C., Mele, F., Jovic, S., Rodriguez, B. F., Gupta, S. V., Jin, F., Piumatti, G., Lo Presti, G., Pellanda, A. F., Biggiogero, M., Tarkowski, M., Pizzuto, M. S., Cameroni, E., Havenar-Daughton, C., Smithey, M., Hong, D., Lepori, V., Albanese, E., Ceschi, A., Bernasconi, E., Elzi, L., Ferrari, P., Garzoni, C., Riva, A., Snell, G., Sallusto, F., Fink, K., Virgin, H. W., Lanzavecchia, A., Corti, D. & Veesler, D. (2020). Mapping neutralizing and immunodominant sites on the SARS-CoV-2 spike receptor-binding domain by structure-guided high-resolution serology. Cell.

66 Yuan, M., Liu, H., Wu, N. C., Lee, C.-C. D., Zhu, X., Zhao, F., Huang, D., Yu, W., Hua, Y., Tien, H., Rogers, T. F., Landais, E., Sok, D., Jardine, J. G., Burton, D. R. & Wilson, I. A. (2020). Structural basis of a shared antibody response to SARS-CoV-2. Science, eabd2321.

67 Yuan, M., Wu, N. C., Zhu, X., Lee, C.-C. D., So, R. T. Y., Lv, H., Mok, C. K. P. & Wilson, I. A. (2020). A highly conserved cryptic epitope in the receptor binding domains of SARS-CoV-2 and SARS-CoV. Science 368, 630–633.

68 Huo, J., Zhao, Y., Ren, J., Zhou, D., Duyvesteyn, H. M. E., Ginn, H. M., Carrique, L., Malinauskas, T., Ruza, R. R., Shah, P. N. M., Tan, T. K., Rijal, P., Coombes, N., Bewley, K. R., Tree, J. A., Radecke, J., Paterson, N. G., Supasa, P., Mongkolsapaya, J., Screaton, G. R., Carroll, M., Townsend, A., Fry, E. E., Owens, R. J. & Stuart, D. I. (2020). Neutralization of SARS-CoV-2 by Destruction of the Prefusion Spike. Cell Host & Microbe 28, 445–454.e446.

69 Zhou, D., Duyvesteyn, H. M. E., Chen, C.-P., Huang, C.-G., Chen, T.-H., Shih, S.-R., Lin, Y.-C., Cheng, C.-Y., Cheng, S.-H., Huang, Y.-C., Lin, T.-Y., Ma, C., Huo, J., Carrique, L., Malinauskas, T., Ruza, R. R., Shah, P. N. M., Tan, T. K., Rijal, P., Donat, R. F., Godwin, K., Buttigieg, K. R., Tree, J. A., Radecke, J., Paterson, N. G., Supasa, P., Mongkolsapaya, J., Screaton, G. R., Carroll, M. W., Gilbert-Jaramillo, J., Knight, M. L., James, W., Owens, R. J., Naismith, J. H., Townsend, A. R., Fry, E. E., Zhao, Y., Ren, J., Stuart, D. I. & Huang, K.-Y. A. (2020). Structural basis for the neutralization of SARS-CoV-2 by an antibody from a convalescent patient. Nature Structural & Molecular Biology 27, 950–958.

70 Robbiani, D. F., Gaebler, C., Muecksch, F., Lorenzi, J. C. C., Wang, Z., Cho, A., Agudelo, M., Barnes, C. O., Gazumyan, A., Finkin, S., Hagglof, T., Oliveira, T. Y., Viant, C., Hurley, A., Hoffmann, H. H., Millard, K. G., Kost, R. G., Cipolla, M., Gordon, K., Bianchini, F., Chen, S. T., Ramos, V., Patel, R., Dizon, J., Shimeliovich, I., Mendoza, P., Hartweger, H., Nogueira, L., Pack, M., Horowitz, J., Schmidt, F., Weisblum, Y., Michailidis, E., Ashbrook, A. W., Waltari, E., Pak, J. E., Huey-Tubman, K. E., Koranda, N., Hoffman, P. R., West, A. P., Jr., Rice, C. M., Hatziioannou, T., Bjorkman, P. J., Bieniasz, P. D., Caskey, M. & Nussenzweig, M. C. (2020). Convergent antibody responses to SARS-CoV-2 in convalescent individuals. Nature 584, 437–442.

71 Brouwer, P. J. M., Caniels, T. G., van der Straten, K., Snitselaar, J. L., Aldon, Y., Bangaru, S., Torres, J. L., Okba, N. M. A., Claireaux, M., Kerster, G., Bentlage, A. E. H., van Haaren, M. M., Guerra, D., Burger, J. A., Schermer, E. E., Verheul, K. D., van der Velde, N., van der Kooi, A., van Schooten, J., van Breemen, M. J., Bijl, T. P. L., Sliepen, K., Aartse, A., Derking, R., Bontjer, I., Kootstra, N. A., Wiersinga, W. J., Vidarsson, G., Haagmans, B. L., Ward, A. B., de Bree, G. J., Sanders, R. W. & van Gils, M. J. (2020). Potent neutralizing antibodies from COVID-19 patients define multiple targets of vulnerability. Science 369, 643–650.

72 Wang, E., Cohen, A. A., Caldera, L. F., Keeffe, J. R., Rorick, A. V., Aida, Y. M., Gnanapragasam, P. N. P., Bjorkman, P. J. & Chakraborty, A. K. (2024). Designed mosaic nanoparticles enhance cross-reactive immune responses in mice. bioRxiv.

73 Wheatley, A. K., Fox, A., Tan, H. X., Juno, J. A., Davenport, M. P., Subbarao, K. & Kent, S. J. (2021). Immune imprinting and SARS-CoV-2 vaccine design. Trends Immunol 42, 956–959.

74 Aguilar-Bretones, M., Fouchier, R. A., Koopmans, M. P. & van Nierop, G. P. (2023). Impact of antigenic evolution and original antigenic sin on SARS-CoV-2 immunity. J Clin Invest 133.

75 Arevalo, C. P., Bolton, M. J., Le Sage, V., Ye, N., Furey, C., Muramatsu, H., Alameh, M. G., Pardi, N., Drapeau, E. M., Parkhouse, K., Garretson, T., Morris, J. S., Moncla, L. H., Tam, Y. K., Fan, S. H. Y., Lakdawala, S. S., Weissman, D. & Hensley, S. E. (2022). A multivalent nucleoside-modified mRNA vaccine against all known influenza virus subtypes. Science 378, 899–904.

76 ter Meulen, J., van den Brink, E. N., Poon, L. L. M., Marissen, W. E., Leung, C. S. W., Cox, F., Cheung, C. Y., Bakker, A. Q., Bogaards, J. A., van Deventer, E., Preiser, W., Doerr, H. W., Chow, V. T., de Kruif, J., Peiris, J. S. M. & Goudsmit, J. (2006). Human Monoclonal Antibody Combination against SARS Coronavirus: Synergy and Coverage of Escape Mutants. PLoS Medicine 3, e237.

77 Scheid, J. F., Barnes, C. O., Eraslan, B., Hudak, A., Keeffe, J. R., Cosimi, L. A., Brown, E. M., Muecksch, F., Weisblum, Y., Zhang, S., Delorey, T., Woolley, A. E., Ghantous, F., Park, S. M., Phillips, D., Tusi, B., Huey-Tubman, K. E., Cohen, A. A., Gnanapragasam, P. N. P., Rzasa, K., Hatziioanno, T., Durney, M. A., Gu, X., Tada, T., Landau, N. R., West, A. P., Jr., Rozenblatt-Rosen, O., Seaman, M. S., Baden, L. R., Graham, D. B., Deguine, J., Bieniasz, P. D., Regev, A., Hung, D., Bjorkman, P. J. & Xavier, R. J. (2021). B cell genomics behind cross-neutralization of SARS-CoV-2 variants and SARS-CoV. Cell 184, 3205–3221 e3224.

78 Bruun, T. U. J., Andersson, A. C., Draper, S. J. & Howarth, M. (2018). Engineering a Rugged Nanoscaffold To Enhance Plug-and-Display Vaccination. ACS Nano 12, 8855–8866.

79 Pear, W. S., Nolan, G. P., Scott, M. L. & Baltimore, D. (1993). Production of high-titer helper-free retroviruses by transient transfection. Proc Natl Acad Sci U S A 90, 8392–8396.

80 Starr, T. N., Greaney, A. J., Hannon, W. W., Loes, A. N., Hauser, K., Dillen, J. R., Ferri, E., Farrell, A. G., Dadonaite, B., McCallum, M., Matreyek, K. A., Corti, D., Veesler, D., Snell, G. & Bloom, J. D. (2022). Shifting mutational constraints in the SARS-CoV-2 receptor-binding domain during viral evolution. Science 377, 420–424.

81 Taylor, A. L. & Starr, T. N. (2023). Deep mutational scans of XBB.1.5 and BQ.1.1 reveal ongoing epistatic drift during SARS-CoV-2 evolution. PLoS Pathog 19, e1011901.

82 Lee, J., Zepeda, S. K., Park, Y. J., Taylor, A. L., Quispe, J., Stewart, C., Leaf, E. M., Treichel, C., Corti, D., King, N. P., Starr, T. N. & Veesler, D. (2023). Broad receptor tropism and immunogenicity of a clade 3 sarbecovirus. Cell Host Microbe 31, 1961–1973 e1911.

83 Wentz, A. E. & Shusta, E. V. (2007). A novel high-throughput screen reveals yeast genes that increase secretion of heterologous proteins. Appl Environ Microbiol 73, 1189–1198.

84 Hsieh, C. L., Goldsmith, J. A., Schaub, J. M., DiVenere, A. M., Kuo, H. C., Javanmardi, K., Le, K. C., Wrapp, D., Lee, A. G., Liu, Y., Chou, C. W., Byrne, P. O., Hjorth, C. K., Johnson, N. V., Ludes-Meyers, J., Nguyen, A. W., Park, J., Wang, N., Amengor, D., Lavinder, J. J., Ippolito, G. C., Maynard, J. A., Finkelstein, I. J. & McLellan, J. S. (2020). Structure-based design of prefusion-stabilized SARS-CoV-2 spikes. Science 369, 1501–1505.

85 West, A. P., Jr., Scharf, L., Horwitz, J., Klein, F., Nussenzweig, M. C. & Bjorkman, P. J. (2013). Computational analysis of anti-HIV-1 antibody neutralization panel data to identify potential functional epitope residues. Proc Natl Acad Sci U S A 110, 10598–10603.

86 Laczko, D., Hogan, M. J., Toulmin, S. A., Hicks, P., Lederer, K., Gaudette, B. T., Castano, D., Amanat, F., Muramatsu, H., Oguin, T. H., 3rd, Ojha, A., Zhang, L., Mu, Z., Parks, R., Manzoni, T. B., Roper, B., Strohmeier, S., Tombacz, I., Arwood, L., Nachbagauer, R., Kariko, K., Greenhouse, J., Pessaint, L., Porto, M., Putman-Taylor, T., Strasbaugh, A., Campbell, T. A., Lin, P. J. C., Tam, Y. K., Sempowski, G. D., Farzan, M., Choe, H., Saunders, K. O., Haynes, B. F., Andersen, H., Eisenlohr, L. C., Weissman, D., Krammer, F., Bates, P., Allman, D., Locci, M. & Pardi, N. (2020). A Single Immunization with Nucleoside-Modified mRNA Vaccines Elicits Strong Cellular and Humoral Immune Responses against SARS-CoV-2 in Mice. Immunity 53, 724–732 e727.

87 Starr, T. N., Greaney, A. J., Addetia, A., Hannon, W. W., Choudhary, M. C., Dingens, A. S., Li, J. Z. & Bloom, J. D. (2021). Prospective mapping of viral mutations that escape antibodies used to treat COVID-19. Science 371, 850–854.

88 Crawford, K. H. D., Eguia, R., Dingens, A. S., Loes, A. N., Malone, K. D., Wolf, C. R., Chu, H. Y., Tortorici, M. A., Veesler, D., Murphy, M., Pettie, D., King, N. P., Balazs, A. B. & Bloom, J. D. (2020). Protocol and Reagents for Pseudotyping Lentiviral Particles with SARS-CoV-2 Spike Protein for Neutralization Assays. Viruses 12.

89 Koday, M. T., Leonard, J. A., Munson, P., Forero, A., Koday, M., Bratt, D. L., Fuller, J. T., Murnane, R., Qin, S. L., Reinhart, T. A., Duus, K., Messaoudi, I., Hartman, A. L., Stefano-Cole, K., Morrison, J., Katze, M. G. & Fuller, D. H. (2017). Multigenic DNA vaccine induces protective cross-reactive T cell responses against heterologous influenza virus in nonhuman primates. Plos One 12.

90 Barnes, C. O., West, A. P., Jr., Huey-Tubman, K. E., Hoffmann, M. A. G., Sharaf, N. G., Hoffman, P. R., Koranda, N., Gristick, H. B., Gaebler, C., Muecksch, F., Lorenzi, J. C. C., Finkin, S., Hagglof, T., Hurley, A., Millard, K. G., Weisblum, Y., Schmidt, F., Hatziioannou, T., Bieniasz, P. D., Caskey, M., Robbiani, D. F., Nussenzweig, M. C. & Bjorkman, P. J. (2020). Structures of Human Antibodies Bound to SARS-CoV-2 Spike Reveal Common Epitopes and Recurrent Features of Antibodies. Cell 182, 828–842 e816.

91 De Genst, E. J., Guilliams, T., Wellens, J., O’Day, E. M., Waudby, C. A., Meehan, S., Dumoulin, M., Hsu, S. T., Cremades, N., Verschueren, K. H., Pardon, E., Wyns, L., Steyaert, J., Christodoulou, J. & Dobson, C. M. (2010). Structure and properties of a complex of alpha-synuclein and a single-domain camelid antibody. J Mol Biol 402, 326–343.

92 Seifert, S. N., Bai, S., Fawcett, S., Norton, E. B., Zwezdaryk, K. J., Robinson, J., Gunn, B. & Letko, M. (2022). An ACE2-dependent Sarbecovirus in Russian bats is resistant to SARS-CoV-2 vaccines. PLoS Pathog 18, e1010828.

93 Hills, R. A., Tan, T. K., Cohen, A. A., Keeffe, J. R., Keeble, A. H., Gnanapragasam, P. N. P., Storm, K. N., Hill, M. L., Liu, S., Gilbert-Jaramillo, J., Afzal, M., Napier, A., James, W. S., Bjorkman, P. J., Townsend, A. R. & Howarth, M. (2023). Multiviral Quartet Nanocages Elicit Broad Anti-Coronavirus Responses for Proactive Vaccinology. bioRxiv.

94 Greaney, A. J., Starr, T. N., Eguia, R. T., Loes, A. N., Khan, K., Karim, F., Cele, S., Bowen, J. E., Logue, J. K., Corti, D., Veesler, D., Chu, H. Y., Sigal, A. & Bloom, J. D. (2022). A SARS-CoV-2 variant elicits an antibody response with a shifted immunodominance hierarchy. PLoS Pathog 18, e1010248.

95 Landau, M., Mayrose, I., Rosenberg, Y., Glaser, F., Martz, E., Pupko, T. & Ben-Tal, N. (2005). ConSurf 2005: the projection of evolutionary conservation scores of residues on protein structures. Nucleic Acids Res 33, W299–302.

96 Letko, M., Marzi, A. & Munster, V. (2020). Functional assessment of cell entry and receptor usage for SARS-CoV-2 and other lineage B betacoronaviruses. Nature Microbiology 5, 562–569.

97 Sievers, F., Wilm, A., Dineen, D., Gibson, T. J., Karplus, K., Li, W., Lopez, R., McWilliam, H., Remmert, M., Soding, J., Thompson, J. D. & Higgins, D. G. (2011). Fast, scalable generation of high-quality protein multiple sequence alignments using Clustal Omega. Mol Syst Biol 7, 539.

